# A language network in the individualized functional connectomes of over 1,000 human brains doing arbitrary tasks

**DOI:** 10.1101/2025.03.29.646067

**Authors:** Cory Shain, Evelina Fedorenko

## Abstract

A century and a half of neuroscience has yielded many divergent theories of the neurobiology of language. Two factors that likely contribute to this situation include (a) conceptual disagreement about language and its component processes, and (b) intrinsic inter-individual variability in the topography of language areas. Recent functional magnetic resonance imaging (fMRI) studies of small numbers of intensively scanned individuals have argued that a language-selective brain network emerges bottom-up from correlations (individualized functional connectomics, iFC) in task-free (e.g., rest) or task-regressed activation timecourses. Here we test this hypothesis at scale and evaluate its practical utility for task-agnostic language localization: we apply iFC separately to each of 1,957 (fMRI) scanning sessions (1,199 unique brains), each consisting of diverse tasks. We find that iFC indeed reveals a largely left-hemisphere-dominant frontotemporal network that is more stable within individuals than between them, robust to the granularity of the parcellation, and selective for language. These results support the hypothesis that this network is a key structure in the functional organization of the adult brain and show that it can be recovered retrospectively from arbitrary imaging data, with implications for neuroscience, neurosurgery, and neural engineering.

## Introduction

Language holds a special place in the study of the human mind and brain. It is a species-specific and species-universal acquired behavior of extraordinary complexity (1), it is intricately linked with human intelligence (2) and social coordination (3), and it is the inaugural domain of cognitive neuroscience (4). Language is also a strong candidate for neural specialization: it can be selectively impaired by developmental disorders (5), brain injury (6), or neurodegeneration (7), with far-reaching consequences for quality of life (8). But despite a century and a half of intensive research, substantial disagreement remains about which brain areas are implicated in language or its component processes and what computations those areas support (9–22).

Two factors likely contribute to this situation. The *first factor* is theoretical disagreement about the construct of language itself and its place in the mind and brain (1, 9, 23–30). It is difficult to synthesize across experiments that disagree fundamentally in how they construe their object of study, given that theoretical precommitments become embedded in task designs. For example, a commonly used task for operationalizing language processing is to compare responses to sentences and responses to lists of pronounceable nonwords (20, 31). However, critics of different theoretical persuasions have argued both that this task is too broad (e.g., that it fails to isolate syntax (32)) and too narrow (e.g., that it fails to include phonological processing (33); cf. (34)). We raise this example not to delve into the particulars of these debates, but to illustrate the difficulty of pursuing a cumulative neurobiology of language in the presence of core disagreements about how language should be conceptualized and studied. The *second factor* is growing evidence that language areas—similar to other areas in the association cortex—are weakly “tethered” to macroanatomical structure (e.g., sulcal/gyral landmarks or coordinates in a template brain) across individuals (10, 31, 35–38). The widely-targeted question of which brain regions support which language functions (12, 13, 15–17, 20, 39, 40) may therefore be ill-posed at the population level, which is the level that standard analytic approaches focus on. One solution to weak structure-function tethering is to identify language regions within individuals using an independent “localizer” task (31, 41–45). Over a decade of work across many labs has shown that diverse language localizers convergently identify a frontotemporal network with strikingly language-selective tuning and distributed sensitivity to a range of linguistic properties (for review, see Ref. (10)). But such studies rely even more fundamentally on the construct and test validity of the localizer task(s). Using a task to localize putative functions (task-based localization) thus accounts for individual variability but pushes functional assumptions deeper into the analysis.

A promising opportunity to skirt both of these issues in studies of the brain’s functional organization (46) comes from functional connectomics (FC), the study of structured covariation in activity between brain areas (47–50), typically using functional magnetic resonance imaging (fMRI). Historically, FC analyses have been performed at the group level (49, 51–54). Motivated in part by the concern above about individual variability in functional organization, recent FC research has moved from studying group-level functional connectomes to individualized ones (iFC) (55–63). In the process, an intriguing finding has emerged: iFC methods seem to recover a network with expected topography (in most people, falling primarily in previously documented frontal and temporal regions with a left-hemisphere bias) that is selective for language (64–67) and closely matches the topography of the language network previously identified and characterized using task-based localization (10).

However, to-date, iFC studies of language have involved relatively small numbers (*n* < 20) of intensively-scanned participants, and inferences about the population drawn from few individuals can be tenuous. Moreover, the bulk of this evidence is derived from a single task state (resting state)—though see Ref. (66). Therefore, claims from iFC about e.g., the existence, topography, and functional properties of a language network would benefit from a larger and more diverse sample of individuals and task states (68–70). Further, on the practical side, any form of data collection that is strictly for localization purposes (including resting state scanning) shortens the time available for the main experiment(s) in the scanning session, and, like localizer tasks, resting state iFC cannot be applied retrospectively to task-only datasets that have already been collected. These considerations limit the practical utility of iFC for functional localization. However, these limitations would be removed if the language network could be detected in *arbitrary* fMRI data, irrespective of task contents. Several lines of prior evidence support this possibility. First, iFC appears to be highly conserved across task states (71–73). Second, the language network as identified using task contrasts exhibits stable connectivity between rest and task (74, 75). And third, iFC studies have directly shown striking recovery of the same frontotemporal language network using task-based iFC (76) in samples of two (64) and fifteen (66) individuals.

In this study, we pushed this insight towards both scale and generality. Towards *scale*, we applied iFC methods to every fMRI session collected in the Fedorenko lab between 2007 and 2024 (excluding data from studies of people with brain lesions or an established diagnosis of a neurodevelopmental disorder, such as autism), a total of 1,957 sessions from 1,199 unique brains (see **Materials and Methods**). This dataset includes two orders of magnitude more individuals than any prior iFC study of language, thereby increasing our ability to make inferences about human brain organization. Towards *generality*, we explored whether a language network is detectable in iFC parcellations under the most general conditions we can simulate: only having access to fMRI timecourses, with no knowledge of the task, from a single scanning session, using fully automated analyses.

To foreshadow our key results, we found that iFC reliably recovers a network (LangFC) with strikingly similar properties to prior findings using task localization (10). In, particular, LangFC (a) shows expected topography, distinct from other brain networks, (b) is selective for language, (c) emerges across task states, (d) differs in function from nearby networks, (e) is robust to the granularity of the parcellation, (f) is stable within individuals and variable between them, (g) can be identified efficiently, and (h) reproduces localizer task-based results. This finding has implications for neuroscience, neurosurgery, and neural engineering (see **Discussion**).

## Results

Drawing on prior FC work that parcellates the brain into distributed networks of strongly functionally connected (covarying in activity over time) areas (51, 64, 65, 77), we developed a simple and fast automated procedure (schematized in **Fig 1**) for computing individualized probabilistic assignments of small-scale volumetric brain regions (2mm isotropic voxels) to functionally-labeled networks. The approach is based purely on low-frequency fluctuations over time in blood oxygen-level dependent (BOLD) measures recorded by fMRI, without reference to task contents. For maximum simplicity and generality, we did not apply task regression (i.e., residualizing the task structure out of the timecourses, see **Discussion**). Evidence from our study (**SI C**) and other recent work (66) suggests that a language network with qualitatively similar functional properties also emerges under task regression.

**Figure 1:**
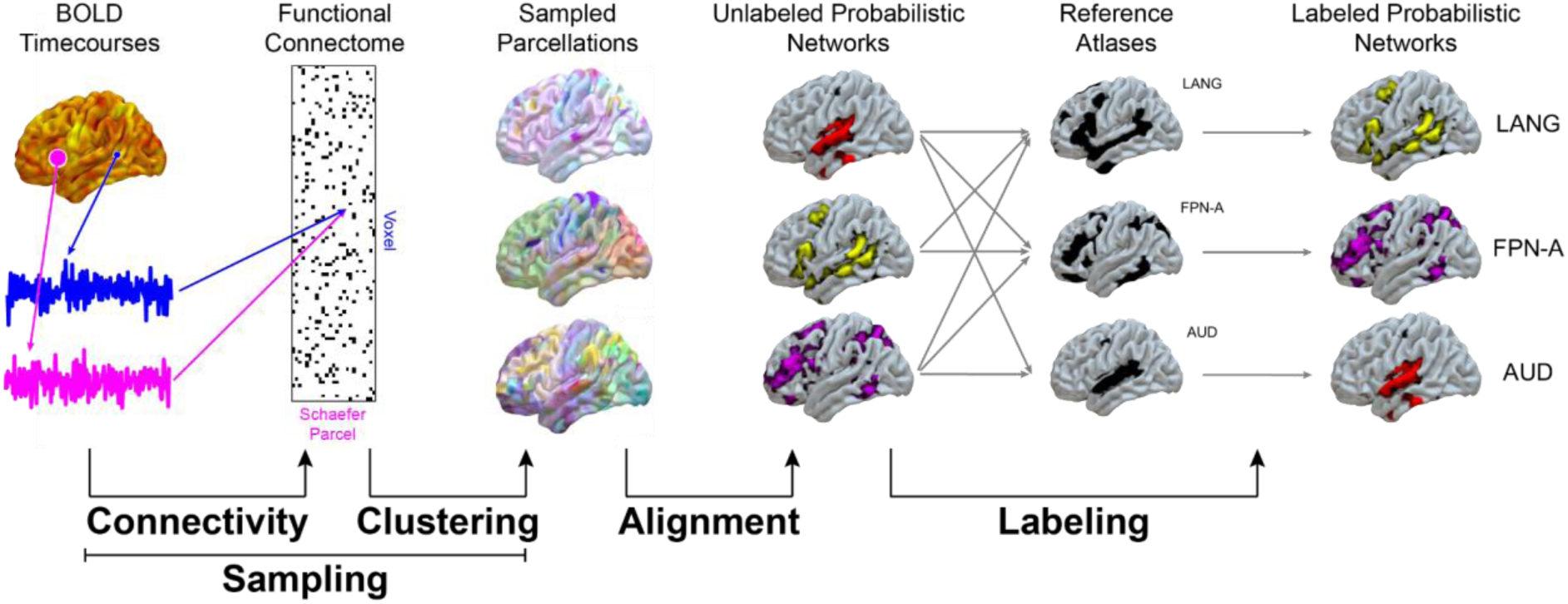
Schematic of our analysis procedure for a single participant. Voxelwise blood oxygen-level dependent (BOLD) timecourses are correlated with average timecourses in each of 1,000 anatomical parcels (53) to produce a correlation matrix (functional connectome), which is then binarized (in the schematic, black dots correspond to the 10% of the strongest voxel-to-voxel correlations). Voxel clusterings (parcellations) are sampled by repeatedly applying *k*-means to the voxelwise patterns of functional connectivity. These samples are then greedily aligned using linear sum assignment (78) and averaged to produce probabilistic assignments of voxels to (unlabeled) networks. Interpretable labels are then assigned to a subset of the networks in the parcellation by finding the most spatially correlated network in the parcellation to each of a set of reference atlases. For details, see **Materials and Methods**.

Our parcellation procedure for a single scanning session was as follows. *First*, we computed a binary connectivity matrix *R* per session (65). To do so, we computed a *V* x 1,000 correlation matrix between the timecourses (times of repetition or TRs, usually corresponding to 2s of data in this study) in *V* three-dimensional voxels in the gray matter volume and the average timecourses in 1,000 parcels of a prior atlas (53). *R* was then computed by binarizing against the 90th percentile across the whole matrix, following Ref. (65). Parcellation results were robust to many of these design choices (**SI D**), generalized to a qualitatively different template-matching method (**SI E**), and generalized across both rest- and task-based iFC in direct comparisons (**SI F**). *Second*, assuming a fixed number of networks *k*, we clustered the voxels into networks by applying *k*-means to *R*. Because *k*-means is sensitive to initial conditions, we derived a *probabilistic parcellation* by computing 100 different clusterings from 100 random initializations. These sampled clusterings are not commensurable: the network boundaries can differ, and even if boundaries are identical, the assignment of labels (1, …, *k*) to networks is random. To address this, in our *third* step, we greedily aligned the clusterings into a common space. To do so, we sorted the clusterings by their mean squared error (best to worst) and initialized the parcellation to the best clustering. We then incrementally modified the parcellation at each subsequent clustering by first using linear sum assignment (78) to optimize the spatial alignment between the clustering and the current state of the parcellation and then updating the parcellation with a moving average. This procedure yields a probabilistic whole-brain map for each of the *k* networks. In each map, the value in each gray matter voxel represents the probability of belonging to the corresponding network. *Fourth*, we extracted 16 functionally labeled networks from the parcellation based on spatial correlation to population-level reference atlases for 16 putative networks proposed by prior work, namely, the Language Atlas (LanA) derived from language localizer task contrasts in a large cohort (79), plus the spatial priors for 15 functional networks derived task-independently in a different sample of participants (65): language (LANG), frontoparietal A and B (FPN-A/B), default A and B (DN-A/B), cingulo-opercular (CG-OP), salience/parietal memory (SAL/PMN), dorsal attention A and B (dATN-A/B), auditory (AUD), premotor posterior-parietal rostral (PM-PPr), somatomotor A and B (SMOT-A/B), and visual central and peripheral (VIS-C/P).

Because the optimal *k* is unknown and may vary between individuals, we optimized *k* for each session by running the entire procedure above independently for *k* = 10, 20, …, 200 and selecting the value of *k* that maximized the average spatial correlation between the labeled iFC networks and their corresponding reference atlases. To avoid *post hoc* selection of results, the final parcellation for a session was computed by resampling a new parcellation at the chosen value of *k*. Importantly, we were not selecting for a particular functional profile for any network: the optimization of *k* is based only on the average spatial similarity to the 16 reference atlases and is therefore independent of any task contrasts available for that participant.

These iFC-derived functional networks were then analyzed with respect to diverse properties, including their responses to key contrasts (*t*-statistic) during a set of tasks that were frequently attested in the dataset and spanned a range of cognitive processes: language (reading and listening), theory of mind (verbal and nonverbal), executive functions (spatial working memory), numerical cognition, music perception, and high-level visual perception. Given our focus on language, a key contrast of interest was the *sentences vs. nonwords* (S-N) contrast in a heavily-studied and widely-used reading-based localizer task for language (31). Therefore, all runs used to estimate the S-N contrast for an individual were excluded from the parcellation procedure, ensuring data independence. iFC and task contrasts were compared using two quantitative measures: (a) Fisher-transformed spatial correlation between an iFC-derived network and the task contrast *t*-map—denoted z(r), and (b) the average *t*-value of the task contrast within an iFC-derived network, weighted by the voxelwise probability of network membership. These measures are complementary: z(r) captures overall *spatial similarity* to the task-evoked response (thus penalizing spatial deviation from the topography of a task contrast, including false negatives: task-responsive areas that fall outside the network), whereas the *t*-value captures the *strength* of the task-evoked response within the network (thus primarily penalizing false positives: task-unresponsive areas that fall into the network).

Detailed methods for parcellation and analysis are provided in **Materials and Methods**.

### LangFC Shows Expected Topography, Distinct from Other Brain Networks

We first compared the topography of our individualized LangFC estimates both to population-level expected topographies (reference atlases) of several key networks and to other iFC-derived networks in the same individual. Unsurprisingly, LangFC has a similar spatial distribution (**Fig 2A**) to the reference atlases used to label it (LANG and LanA, **Fig 2C**). More noteworthy is the fact that it has low or negative spatial correlation with the other fourteen reference atlases, indicating that LangFC was generally well separated from population-level expected topographies of networks that have been previously associated with nonlinguistic functions. When comparing LangFC to the set of iFC-derived networks in the same individual instead of reference atlases, this spatial separation is substantially enhanced (**Fig 2B**): LangFC is nearly identical within individuals whether labeled relative to LANG (65) or LanA (79), with weak or negative spatial correlation to other iFC-defined networks in the same individual. Note that, due to our probabilistic network definitions, different iFC networks can in principle overlap spatially; the fact that they generally do not is a contingent outcome. The finding of similar LangFC outcomes regardless of reference atlas (LANG or LanA) is noteworthy given that the LANG and LanA reference atlases differ meaningfully: although they are visually similar (**Fig 2B**), their z(r) to each other is only 0.37, and thus they could in principle systematically pick out different networks in the parcellation. But, in practice, LANG and LanA references picked out the same network in 58% of sessions in our study, and in the remaining sessions they identified networks with a similar degree of spatial similarity—average z(r)=0.39±0.01 SEM—to the similarity between the LANG and LanA reference atlases themselves.

**Figure 2:**
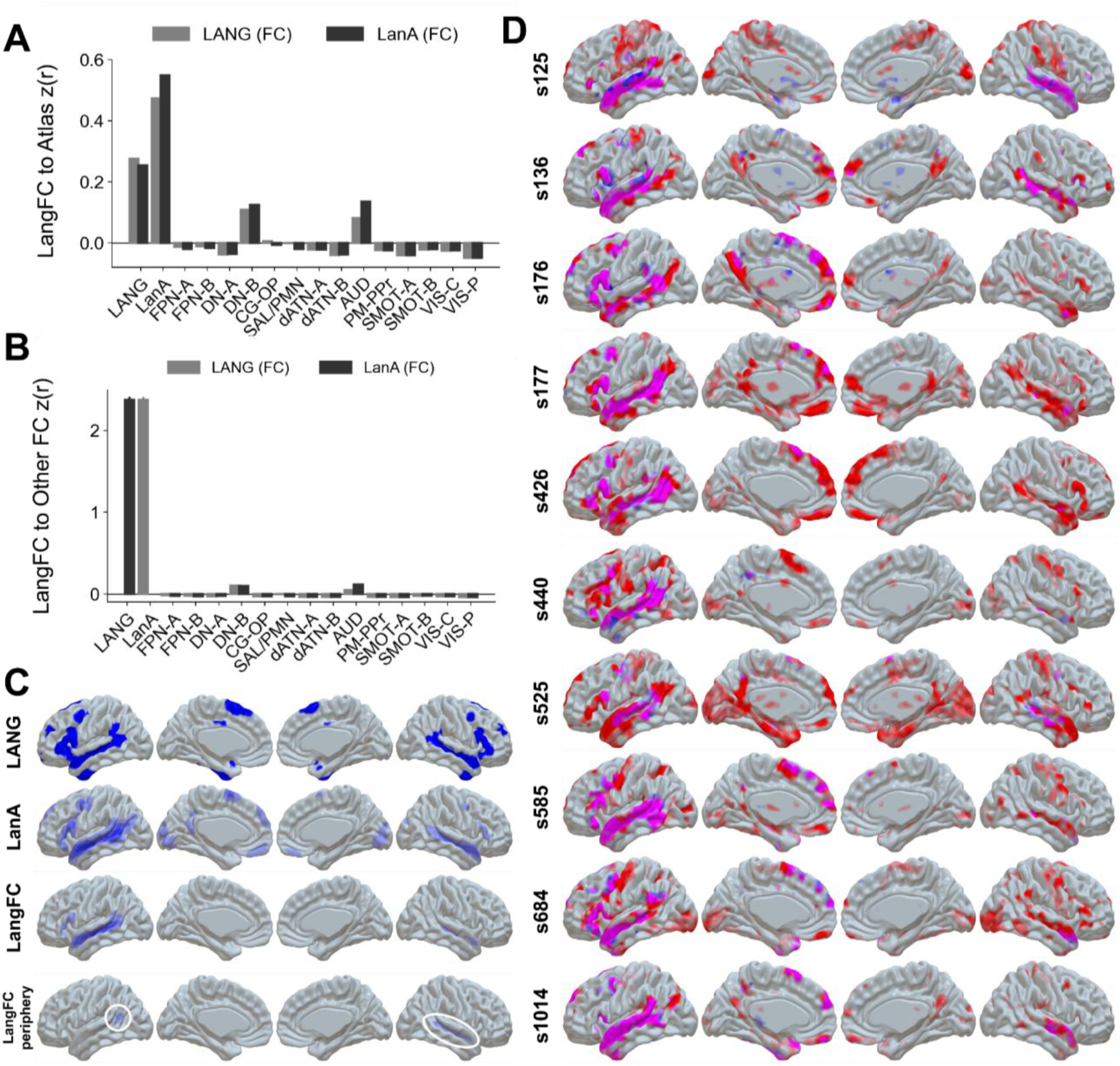
**A.** Individual-level LangFC networks that are labeled relative to LANG (light) and LanA (dark) are spatially similar to the LANG and LanA reference atlases (this is expected given that these atlases are used to label LangFC). Similarity to the other reference atlases is low, with weak similarity to two networks (DMN-B and AUD) that are known to abut language areas within individuals. **B.** Within individuals, LangFC is nearly identical whether labeled using LANG or LanA, and spatially dissimilar from other FC-derived networks. **A-B.** Error intervals represent standard errors of the mean (SEMs), which in some cases are too small to be visible given the size of our sample. **C.** The LANG and LanA reference atlases used to automatically label LangFC from the data-driven parcellation, alongside both the group average in our study of binarized (*p* > 0.5) LangFC and an estimate of LangFC’s “periphery”, with faint trends highlighted with white ellipses in the left temporoparietal area and the right superior temporal sulcus (see **Materials and Methods**). The LANG and LanA atlases are broadly similar, as is the group average of LangFC. The periphery is faint but tends to fall in right-hemisphere homotopes of language areas, as well as a left-hemisphere area posterior to temporal language areas, consistent with prior task-based evidence (80). **D.** LangFC (blue) vs. task *t*-maps (sentences vs. nonword lists or *S-N*, red) in the 10 participants with the highest S-N contrast stability between even and odd runs. Overlap is shown in magenta, and opacity reflects magnitude (0.2 < *p* < 0.8 for LangFC, 1 < *t* < 4 for S-N). LangFC overlaps substantially with the S-N contrast (and tends to be contained within it) and follows inter-individual variation in the topography of task activations.

The two most topographically similar networks to LangFC are AUD and DMN-B, both of which are known to abut temporal language regions within individuals (67, 81). In addition, when we consider an “extended” LangFC (82) composed of multiple iFC subnetworks whose union maximizes spatial similarity to the reference atlas (LANG or LanA, see **Materials and Methods**), we find (**Fig 2C**, bottom) that LangFC’s periphery tends to fall both in right hemisphere homotopes of language areas and in left-hemisphere areas posterior to temporal language areas. This finding is consistent with previous task-based reports about the distribution of the language network’s “periphery” (80, 82) or “weak shadow” (83).

Consistent with prior claims (64), LangFC (**Fig 2C**, bottom) systematically includes areas in the middle frontal / superior precentral gyri as well as dorsomedial frontal areas that fall outside of traditional theoretical models of language neurobiology (13), while also excluding temporoparietal areas that are often included in language neuroscience studies (20, 84). The inclusion of temporoparietal areas in language studies is likely because these areas often register responses to meaningful language, as reflected by their presence in the task-derived LanA atlas (**Fig 2B**, bottom). But prior work has reported that language-responsive temporoparietal areas differ functionally from the language network in several respects and are thus arguably not part of the network (64, 80), a claim that finds support from our iFC results.

LangFC also shows the left-hemisphere bias abundantly attested in prior work on language in the human brain (4, 10, 13, 40, 64, 85–91), as can be seen qualitatively in **Fig 2D** and **SI A**. We quantified this impression by computing a *laterality index* (LI) defined as the ratio of the interhemispheric difference to the interhemispheric sum in some measurement (92), thus ranging from -1 (fully right-lateralized) to 1 (fully left-lateralized). We found comparable degrees of strong left-hemisphere bias across participants (averaging across sessions within participants) in LangFC (number of voxels exceeding p=0.5 of belonging to LangFC in the parcellation, LI=0.61±0.01SEM) and the reading localizer task contrast (number of voxels exceeding t=5 in the sentences–nonwords contrast, LI=0.60±0.01SEM).

### LangFC Is Selective for Language

The individualized spatial distributions of LangFC and the S-N contrast—which are derived from independent data—are highly overlapping (especially in superior temporal, inferior and middle frontal, and dorsomedial frontal areas), as shown qualitatively in a sample of participants (**Fig 2D**, see **SI A** for exhaustive results) and quantitatively across participants (**Fig 3A**). LangFC is spatially correlated—high z(r), **Fig 3A** top—with contrasts evoked by language tasks but not tasks targeting theory of mind, executive functions, numerical cognition, music, or high-level visual processing. Likewise, the contrasts themselves—*t*-values, **Fig 3A** bottom—are large for language tasks but not the other tasks. The only nonlinguistic task that evokes a substantial response in LangFC is the verbal theory of mind task, but this task is reading-based (and thus mediated by language comprehension, unlike its non-verbal counterpart) and has been shown in prior work to be confounded with dimensions of linguistic complexity that are independently known to drive responses in language areas (80).

**Figure 3:**
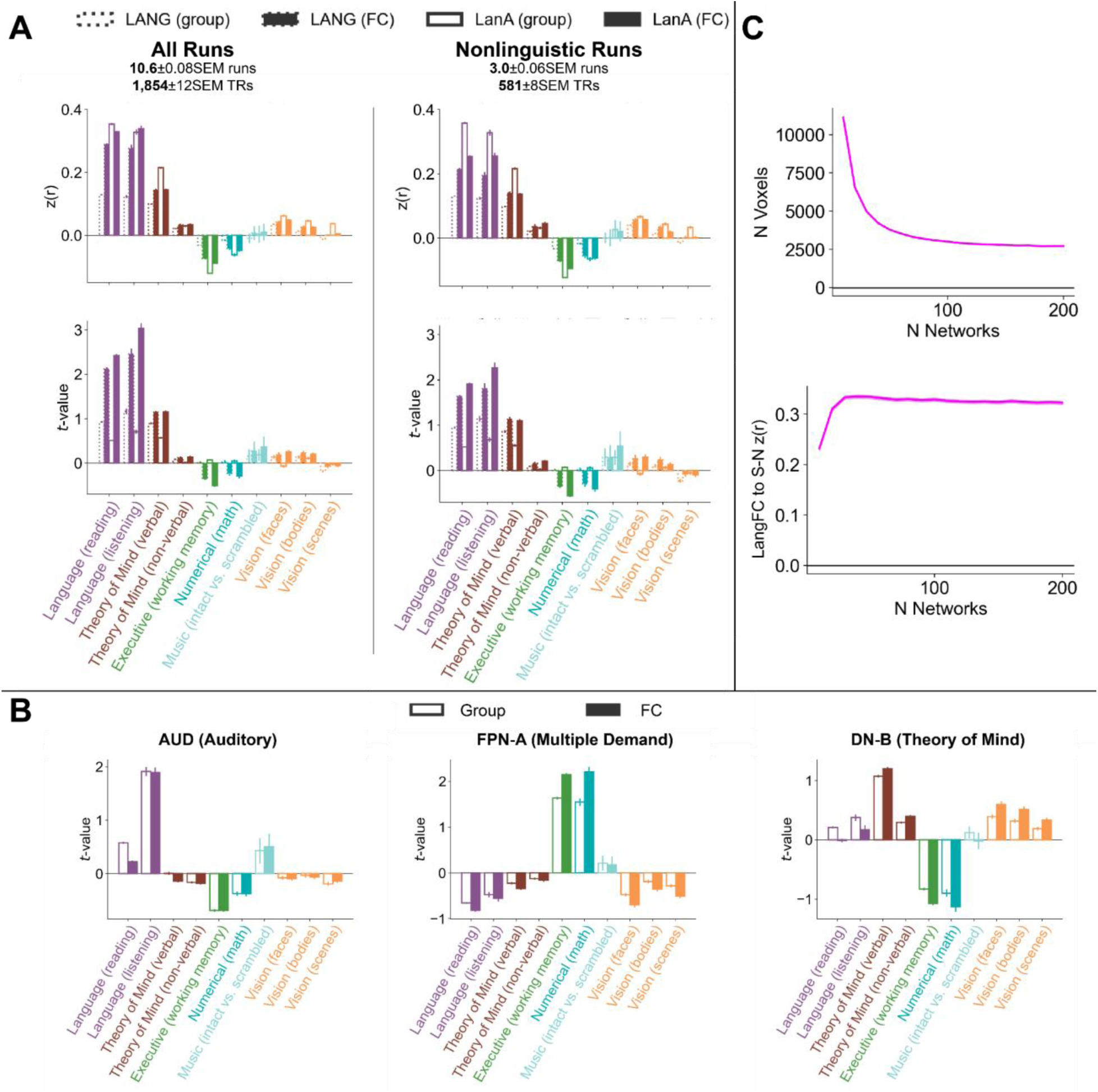
**A.** FCLang is more spatially similar to language vs. other contrasts (top) and likewise has a stronger average response to language vs. other contrasts (bottom), whether parcellated from all runs (left) or from nonlinguistic runs only (right). The larger z(r) and *t* values for verbal theory of mind can be explained by linguistic confounds in the task (80). **B.** Other iFC networks show a different functional profile. AUD only shows a strong response in an auditory task, FPN-A (which corresponds to the Multiple Demand (MD) network; (65)) doubly dissociates from LangFC (positive response to executive and numerical tasks and negative response to language tasks), and DN-B (which corresponds to the Theory of Mind (ToM) network; (65)) responds most strongly to theory of mind tasks and shows the default network’s signature task negativity (strong deactivation under demand during executive and numerical tasks). **C.** The language network is stable over a large range of parcellation granularities, with little decrease in the number of voxels (sum of LangFC probabilities) or change in spatial similarity to the reading localizer contrast map (S-N) beyond about 50 networks. **All panels:** Error intervals represent standard errors of the mean (SEMs), which in some cases are too small to be seen given the size of our sample.

LangFC is also more functionally selective than the reference atlases used to label it: it has higher z(r) and *t*-value for language tasks than LANG (filled vs. unfilled dotted bars in **Fig 2A**). Although LangFC has comparable z(r) in language tasks to the LANA atlas, it is more functionally selective (higher *t*-value, filled vs. unfilled solid bars in **Fig 2A**). LanA therefore tends to capture the spatial distribution of language task responses reasonably well (likely by virtue of being more continuously-valued and covering a greater spatial extent than LANG, **Fig 2C**), but it does so at the expense of functional selectivity. Only LangFC—which is individualized—both captures the spatial distribution of task responses and reliably excludes functionally irrelevant areas. Moreover, in supplementary analyses (**SI B**), we found that our approach to labeling LangFC from among the set of networks in the parcellation was on par with an “oracle” setting that directly selected the network most similar to the participant’s reading language localizer contrast map. This outcome suggests that our labeling approach nears the participant-specific optimum for this dataset. The importance of identifying language areas within individual brains evidenced here aligns with prior claims of weak structure-function tethering for language across individuals (35, 36, 38).

### LangFC Emerges Across Task States

**Fig 3A** shows evidence that functionally relevant stimulation (language) is not needed to identify LangFC: even when we parcellate only based on tasks that involved no language in input or response (e.g., a spatial working memory task or a face perception task; see **Materials and Methods** for details about the nonlinguistic tasks), we recover a variant of LangFC (“Nonlinguistic Runs”). The recovered network is qualitatively similar in spatial distribution and functional profile to LangFC derived from all runs to the exclusion of the S-N task contrast (“All Runs”), albeit with some degradation in the selectivity of the network’s correspondence to language task contrasts. Note that this is a conservative test: in the nonlinguistic condition, we are not only excluding functionally relevant data, but we are also parcellating on less than a third of the data on average (mean 581 timepoints, rather than 1,854), which can harm parcellation quality (see e.g., Ref. (65) and this study, **LangFC Can Be Identified Efficiently**). In supplementary analyses (**SI C**), we found similar estimates of LangFC regardless of whether we regressed out the task structure. These findings converge to suggest that LangFC connectivity is strong enough to enable good recovery of the network across diverse task states.

### LangFC Differs in Function from Nearby Networks

LangFC’s (language-selective) functional profile is different from the functional profiles of networks known to be spatially adjacent to language areas within individual brains (**Fig 3B**), including the auditory network (AUD), fronto-parietal network A (FPN-A, sometimes called the *multiple demand* (MD) network (93, 94)), and default network B (DN-B, sometimes called the *theory of mind* (ToM) network (95)). AUD only shows a substantial response to the listening-based language localizer, which contrasts intact speech with unintelligibly degraded speech (41). Although this contrast is generally effective for localizing high-level language areas, it does not perfectly match acoustic features across conditions, and therefore activates nonlinguistic auditory areas, including speech-selective areas (96, 97), to some extent (67). AUD shows only a weak response to a music task, plausibly because many auditory areas, including speech-processing areas, are not sensitive to music structure (e.g., Ref. (97)) and thus respond similarly in both the typical and structure-scrambled conditions. FPN-A shows a strong response to working memory and math tasks and weak or negative responses to all other tasks. Its response is anticorrelated with the language localizer task, which is a known property of this network, to the point that the reverse contrast (nonwords vs. sentences) can be used as an MD network localizer (98). FPN-A and LangFC thus doubly dissociate with respect to their language vs. executive/numerical task responses. DN-B shows little response to language tasks, is strongly activated by theory of mind tasks, and is strongly deactivated by executive/numerical tasks. All of these properties are expected based on prior characterization of DN-B’s function (65, 66, 99–101).

### LangFC Is Robust to the Granularity of the Parcellation

We found (**Fig 3C**) that the size and shape of LangFC converged to a stable configuration around a granularity of approximately 50 networks in the parcellation and remained stable over a range of up to 200 networks, considerably more than the 15 networks used in recent iFC studies of language (65, 66). Despite the increasing granularity of the parcellation, LangFC did not appreciably fractionate, but instead retained a stable count of about 2,000 voxels on average (computed as the sum of LangFC probabilities across the gray matter volume) and a stable degree of spatial similarity to the S-N contrast within individuals. This result reinforces prior claims that the language network is tightly functionally integrated, rather than showing sharp network-internal functional divides (10).

### LangFC Is Stable Within Individuals and Variable Between Them

We found (**Fig 4A**) that both LangFC and the *t*-maps for the S-N task contrast were stable within individuals across scanning sessions but variable between individuals, consistent with prior reports (36, 64). Moreover, within individuals, we found that LangFC was more stable than the S-N contrast when evaluated across the entire brain, suggesting that the iFC approach successfully “cleans up” some of the noisy parts of task-based maps (see **SI A**). However, when restricting the analysis to six left-hemisphere fronto-temporal regional masks used in prior work (36), we found that S-N contrasts became more stable than LangFC, although LangFC remained stable. Thus, relative to a localizer task, LangFC seems to provide less noisy localization at the whole brain level, but slightly noisier localization within brain areas likely to contain the “core” language network (82).

**Figure 4:**
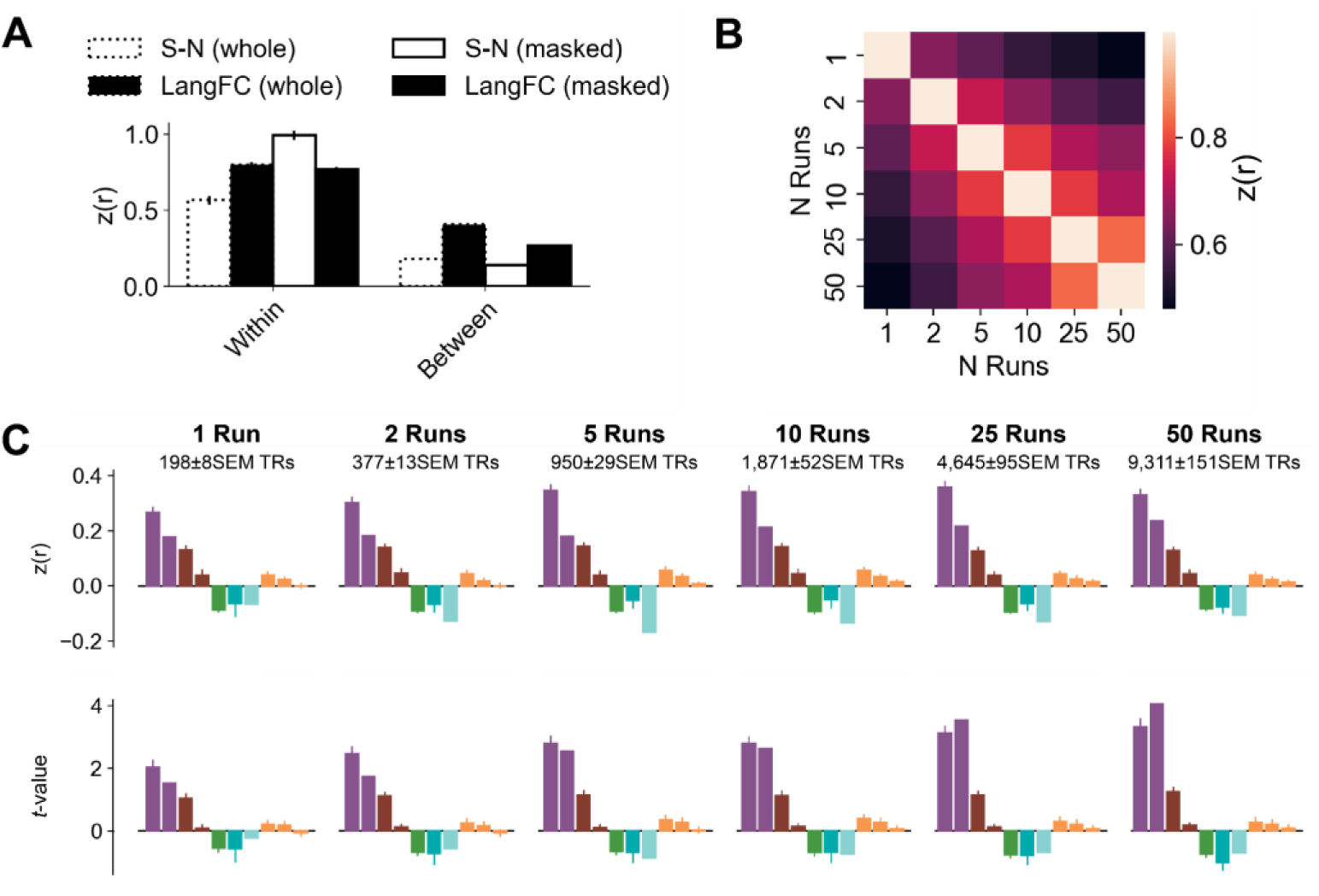
**A.** Both LangFC and the S-N contrast *t*-maps are more similar (a) between sessions within individuals vs. (b) between individuals, indicating substantial individual variability in structure-function tethering for language. LangFC is similarly stable whether considering the whole brain (whole) or only six left-hemisphere regional masks (masked), whereas S-N is more stable under masking, at least within individuals. **B.** The matrix of pairwise z(r) between LangFC as estimated from different numbers of runs (1, 2, 5, 10, 25, and 50) shows that LangFC stability increases with the number of runs used for parcellation. LangFC estimates from larger numbers of runs (especially 5 or more, bottom right of the plot) have greater Fisher-averaged correlation with each other. **C.** LangFC shows a language-selective profile of task responses (see Fig 2 for task labels) even when estimated from a single run (left), but its functional selectivity is enhanced by parcellating on more data (right), especially in the strength of the network’s language task response (*t*-value).

### LangFC Can Be Identified Efficiently

To assess how much data within an individual is needed to identify LangFC, we ran a secondary analysis of the subset of the participants in the dataset (n=53) who had at least 50 runs (on average, at least 5 hours of imaging data) in total across all scanning sessions. We parcellated each individual based only on the first *m* runs, *m* ∈ {1, 2, 5, 10, 25, 50} (when *m* = 50, mean 9,311±151SEM TRs). We found (**Fig 4B**) that increasing the number of runs helped stabilize the LangFC estimate, consistent with prior work (65): LangFC estimates based on larger numbers of runs (especially 5 or more) are more spatially similar to each other. Nonetheless, LangFC is already functionally selective even when estimated from a single run (mean 198±8SEM TRs, **Fig 4C**, left), although selectivity becomes more pronounced with more data (**Fig 4C**, right). These results suggest that LangFC can be identified to a reasonable approximation even with little data, but that parcellation quality will likely benefit from additional data when available. Importantly, if task-agnostic iFC is being applied retrospectively, it requires no data beyond the data already being collected for the main experiment(s).

### LangFC Reproduces Localizer Task-Based Results

We have confirmed that iFC recovers a language selective network with favorable characteristics across a range of measures of reliability, stability, and efficiency. These results suggest that iFC can be used instead of task-based localization to identify the language network within individuals. We put this suggestion to direct test by using LangFC as a drop-in replacement for the S-N *t*-map in a reanalysis of a prior study in the lab that used task-based localization (102). In brief, the experiment was a reading study in which participants read variable-length (1- to 12-word) “chunks” that formed either complete phrases (real-words condition), “syntactic prose” in which real words were replaced with plausible nonwords (Jabberwocky condition), or partial syntactic phrases (non-constituents condition; see the original study for additional details). In one analysis of the study, the top 10% of voxels with the largest S-N *t*-value within individuals were identified for each of the six regional parcels shown in **Fig 5A**, resulting in individualized *functional regions of interest* (fROIs). In the original study, it was found that task-based functional localization dramatically increased signal strength for the key effects (**Fig 5B**) relative to group-level ROIs (**Fig 5D**). When we instead functionally localized based on LangFC (**Fig 5C**), we largely recovered the favorable sensitivity properties of task-based localization, especially in (e.g., inferior frontal) regions where sensitivity suffers most in group-level analyses (103). Quantitatively, when we looked across the six regions in this analysis, we found a small but significant decrease in the slope of the length effect in the real-word conditions from iFC relative to direct task-based localization (mean=0.39, *t*=2.4, *p*=0.02) but a larger significant increase in this same slope from iFC relative to the anatomical group-level parcels (mean=0.60, *t*=5.1, *p*<0.001). This result supports the practical feasibility of using LangFC to find language areas in individual brains and study their response properties, including potentially in retrospective analyses of existing data and in participant populations for whom additional task-based localizer scans would be burdensome.

**Figure 5:**
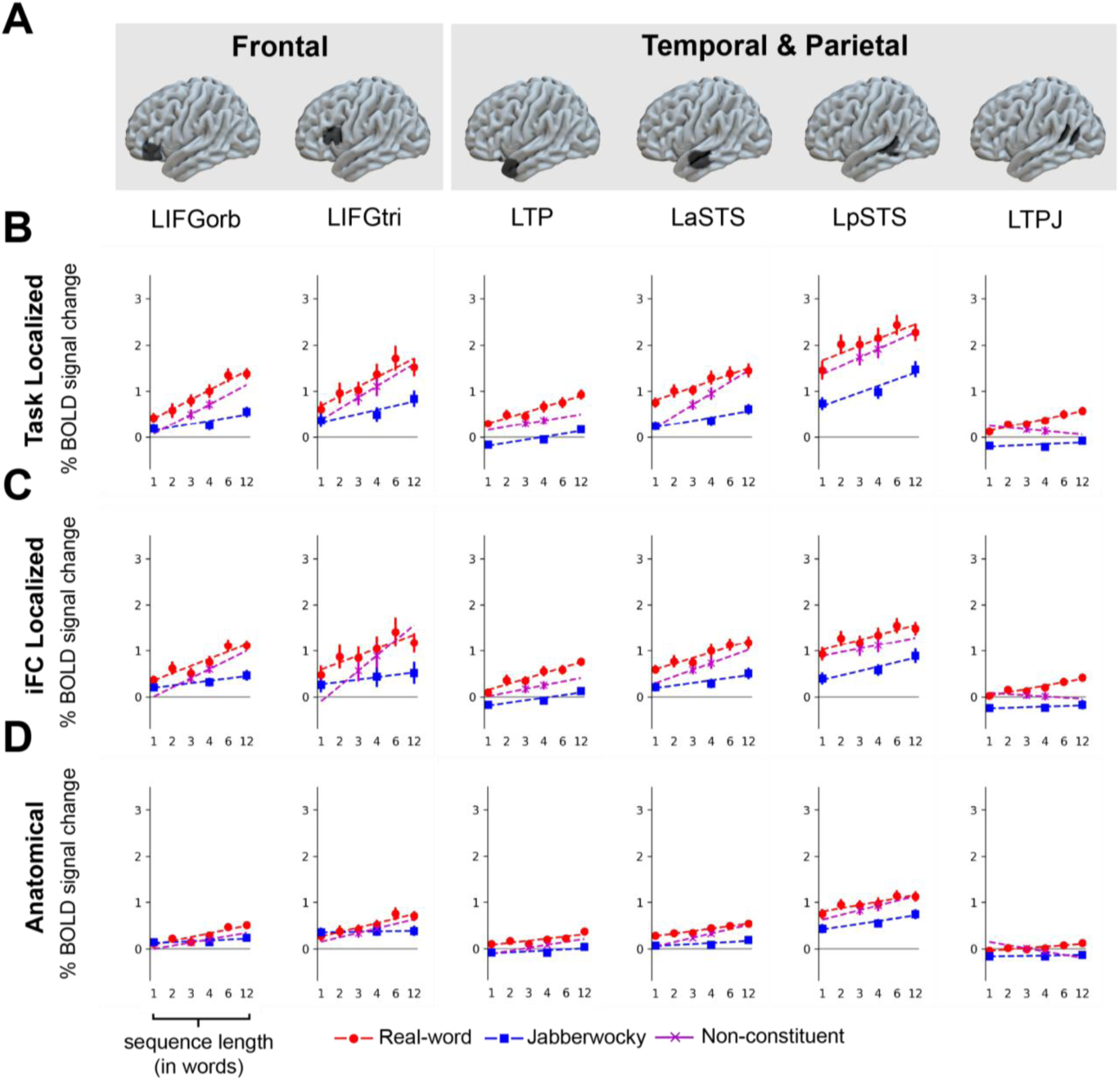
Reanalysis of a prior study (102) in six regional masks (**A**) using individualized task-based (S-N) localization (**B**) or iFC-based localization (**C**) within these masks, relative to averaging over the whole mask (**D**). Panels B and D are as reported in Ref. (102). The task is reading variable-length sequences of complete phrases (real-word), syntactic prose (Jabberwocky), and partial phrases (non-constituent). Even though this task is highly similar to the S-N localizer used in B, iFC is nearly as sensitive as task-based localization, and more sensitive than whole-mask averaging, especially in regions (e.g., IFG) where whole-mask averaging most attenuates the signal. **B-D.** Error intervals represent standard errors of the mean (SEMs).

## Discussion

We have argued that contradictory theoretical precommitments and widespread reliance on group-based functional mapping present barriers to progress in language neuroscience, and we have evaluated the potential of individualized functional connectomics (iFC) to skirt these issues and offer a path forward (46). In so doing, we have built on prior evidence to this effect (64–67) while going substantially beyond it in scale and generality, simulating widely-applicable conditions across diverse practical needs in functional neuroimaging (access only to arbitrary functional timecourses from a single scanning session) in the largest sample we could obtain (nearly 1,200 participants). Similar to other work (104, 105), we used an entirely automated process for parcellating individual brains into interpretably labeled networks with no task supervision. This design enabled our core contribution: we assessed at scale the extent to which recent claims that a network for language is a key unit of human brain organization (10)—which to date are based heavily on task fMRI, though see refs (64–67)—hold up under weaker assumptions. In this respect, our focus is primarily on the nature of human brain organization rather than on methodological approaches to studying this organization: we are not advocating a particular approach to functional brain parcellation and indeed expect, and show empirically, that the qualitative pattern generalizes across methods (**SI D** and **E**). Nonetheless, our claims have important practical implications because they support the use of iFC to identify the language network within individuals in a broad range of both prospective and retrospective settings, including the unfortunately common case where the task parameters associated with a given run (e.g., in an open dataset) are missing, incomplete, or poorly documented.

Specifically, we found that iFC systematically yielded a network with broadly similar topography to expectations established by prior work (in most brains, the network was left-lateralized and fell primarily in the temporal and frontal lobes). We further found that this network was selective for language, different in functional tuning (even doubly dissociating) from networks closely adjacent to it, robust to parcellation granularity (suggesting tight functional integration across the network), stable across scanning sessions within individuals, and variable between individuals. This picture of the neurobiology of language—which amounts to an iFC-based recapitulation of the picture advocated in a recent position paper based largely on task data (10)—diverges from standard theorizing in language neuroscience, which attempts to assign specific linguistic functions to macroanatomically defined regions or pathways in the brain (12, 13, 15, 19, 20, 106–110). Instead, we find iFC evidence convergent with prior claims (36) that the stable trait-level topography of language areas differs substantially between brains (**Fig 4A**). It follows from this that the question of which (macroanatomical) regions support which (linguistic) functions is ill-posed, at least for the population as a whole, which might partially underlie ongoing confusion about the basic neurobiology of language (11). We have emphasized the left-dominance and frontotemporal distribution of LangFC merely in order to highlight continuities between our findings and those of decades of prior research in language neurobiology. But in so doing, we do not intend to imply that the language network is exclusively left-lateral or exclusively frontotemporal. Even within our own cohort we find qualitative evidence (**SI A**) of both (a) substantial individual variability in lateralization of both task and LangFC topographies (90, 111) and (b) LangFC and task topographies that frequently extend into other parts of the brain, consistent with recent arguments (112–114). Quantifying and explaining these observations fall outside our present goals, and we leave them to future work.

Our finding that LangFC is highly selective for language (**Fig 3A**) is a truly contingent outcome of our analyses that converges both with recent iFC work (64–67) and with task-based studies (80, 98, 101, 115–117). We did not seek (and could not in principle have sought) this result in the design of our procedure, given that nothing constrains the functional connectome to show any particular task tuning. This finding is therefore strong independent confirmation of the hypothesis that an integrated frontotemporal network for language is a key unit of adult brain organization, independently of how language is construed theoretically or operationalized experimentally by researchers. In other words, iFC offers language neuroscientists a theory-external explanatory target in the brain, much as the discovery of the default network(s) has done for the functions that they support (47, 99, 100, 118). It follows from this that the term “language network” is demoted from definitional to descriptive: it refers to a brain network with independent existence that could instead be called e.g., “the frontotemporal network” without loss of accuracy. Such a label would belie the network’s strikingly language-selective tuning, and we therefore have continued to include “language” in our terminology. But our claims about this network would not be fundamentally altered by future nuances in our understanding of its function, such as recent evidence that this network is also engaged to some extent by nonverbal event semantics (119, 120).

Our findings also bear indirectly on influential claims that distinct linguistic subfunctions (e.g., processing of phonology, words, syntax, and sentence meaning) are implemented by different macroanatomically-defined (e.g., inferior frontal vs. posterior temporal) regions (12, 13, 20, 106). Any such sharp divisions have proven difficult to replicate within individuals (31, 102, 121–123), and our findings converge with recent iFC work (64–67) in suggesting a reason why: the entire network is tightly functionally integrated, as revealed by the fact that it is parcellated out of the functional connectome as a single unit. We furthermore have shown that this is not merely an artifact of choosing a particular parcellation granularity: even when the iFC approach has 200 networks to work with (and thus plenty of opportunity to fractionate the language network into component networks or regions, if they existed), it prefers to keep LangFC intact (**Fig 3C**). This result reduces the plausibility of any claims about sharp network-internal functional subdivisions at scales detectable using fMRI: any claimed subdivisions would need to explain how the activations across the putative divide are so highly correlated. This finding of course does not obviate questions about how the network implements different linguistic subprocesses. It simply encourages a reorientation away from macroanatomical divisions and toward other possible types of differentiation, including structure at finer spatial scales (124, 125), in time (126), in the core vs. periphery (82), in cortical gradients (127), and in distributed patterns of activation (128).

Beyond these findings and their bearing on theoretical questions about the neurobiology of language, we investigated the practical utility of iFC methods for language network identification in plausible real-world conditions in which individual-participant datasets are small or task information is unavailable or unreliable, leaving only the possibility of parcellating directly on timecourses with unknown task content, without the task regression (76) commonly used in other iFC work (64, 66, 72). Although task regression is known to improve the ability to estimate the resting state functional connectome from task data (76), the goal of iFC-based language localization is not to simulate rest but instead to identify the language network under minimal assumptions. We found that iFC showed favorable stability and efficiency characteristics that support its practical utility for functional localization: LangFC is topographically similar to language task contrasts in unseen data, is robust to the presence or absence of functionally relevant stimulation, and remains spatially and functionally similar with or without task regression in direct comparisons. Moreover, we found that LangFC can be identified to a good approximation even from only a single functional run, while also measurably benefiting from additional data when available. Finally, we showed that LangFC works well as a drop-in replacement for localizer task contrast maps in an existing functional localization toolchain (129, 130). Together, these results suggest a wide domain of practical applications for iFC as a theory-neutral language localization method. Retrospective iFC unlocks thousands of hours of existing fMRI datasets for individualized functional analysis, even if those datasets lack the localizer task contrasts that would otherwise be a prerequisite. It also could support such analyses in studies facing practical constraints on data quantity, such as participant populations with low tolerance for extended scanning (42). In addition, iFC has potential as a technique to identify eloquent cortex prior to surgical resection (131–134), to select sensor placements for neural speech prostheses (135–137), or to select sites for language-related brain stimulation (138).

Our study has several limitations, some of which are plausible targets for future work. *First*, our automated methods may introduce noise either in the mapping from the functional connectome to a parcellation or in the labeling of LangFC. Errors in parcellation may arise because we have used a parcellation method (*k*-means) that can converge to local optima (this issue also holds of other widely-used parcellation methods, including independent components analysis and dictionary learning). We have mitigated (but not eliminated) this issue by sampling and aligning many parcellations with different random seeds. Errors in labeling may arise if an individual’s language network is successfully parcellated but happens not to be the most spatially similar network to the reference atlas used to label LangFC. We have chosen to accept the possibility of such errors for this study in order to achieve scale and minimize researcher influence on results. But both of these steps can be replaced by manual expert checks of both the continuous functional connectome (seed-based analysis) and the topographic plausibility of candidate LangFC networks. Previous iFC studies have demonstrated that automatic and manual approaches to identifying the language network tend to align closely (64, 65). Nonetheless, in smaller scale studies, identification of LangFC based on expert review of the functional connectome is always available.

*Second*, we have only explored a single basic approach to FC parcellation (*k*-means), without engaging with significant recent efforts to increase the quality and sophistication of FC parcellation methods (104, 138–141). We have done so for strictly practical reasons (we needed fast methods to work at this scale of data), but we expect that our results provide a conservative picture of the utility of iFC for language localization that will be surpassed by integrating with ongoing technical advances in the field.

*Third*, although we have advocated the use of LangFC for task-agnostic language localization, we are not advocating a wholesale abandonment of localizer tasks. Even if iFC is used (rather than localizer tasks) for selective analysis, it is valuable to have localizer task data in order to validate LangFC estimates. Without such validation, there would be no direct confirmation of LangFC’s functional tuning within an individual, and results would need to be interpreted more cautiously. In other words, two lines of evidence are better than one: while we have shown that iFC is now essentially always available for language localization (given that it can be reliably estimated from arbitrary task states), additional task data when available can improve confidence in the localization by providing independent convergent evidence. Moreover, localizer tasks provide an estimate of the strength of response during language processing; such estimates can be useful for comparison with other experimental conditions, and also constitute a neural marker (reliable over time; Ref. (36)) for investigations of brain-behavior or brain-genetic relationships across individuals. Thus, for studies targeting language for which it is feasible to collect a localizer task, doing so can still have value, whereas for studies in which language localization is lower priority and/or running additional localizer tasks is infeasible, we have shown that iFC can usually serve as a strong stand-in.

*Fourth*, we have not engaged with (and cannot speak to) more nuanced questions about the functional connectome itself, including widespread interest in (a) its dynamics (142–152) and (b) its multivariate interactions (153–156). Our approach assumes based on prior evidence that much of the functional connectome is stationary (72) and that networks can be recovered from univariate voxel-level timecourses (65), and we leverage these assumptions to identify stable language-relevant brain areas within individuals. But we leave open the possibility that e.g., functional connectivity can vary over time and between task states in ways that meaningfully bear on the brain basis of language. The existence of reliable iFC methods for language neuroscience opens up a new window onto these questions for future work.

## Materials and Methods

### fMRI Data Acquisition and Preprocessing

All structural and functional data were collected on a whole-body 3 Tesla Siemens scanner with a 32-channel head coil at the Athinoula A. Martinos Imaging Center at the McGovern Institute for Brain Research at MIT. Because our dataset covers the entire fMRI acquisition history of a neuroscience lab (see **fMRI Session Selection**), it includes data collected over a long timespan for diverse projects with diverse goals (including unpublished pilot studies), and thus the acquisition parameters vary. There were two large-scale changes in equipment and/or lab-standard acquisition parameters, one in 2021, and one in 2022. Typical scan parameters in each of these three time periods are given in **Table 1**. Detailed scan parameters for each run in the dataset are provided in our Stanford Digital Repository dataset (**URL**).

**Table 1:** Typical MRI acquisition parameters used in this study. Counts in “number of runs” are provided for convenience and include near-matches to these typical configurations. Precise parameters for each run in the dataset are provided in our Stanford Digital Repository dataset (URL).

|  | ~2021 and before | ~2021 to ~2022 | ~2022 to present |
| --- | --- | --- | --- |
| Scanner Model | Siemens Trio | Siemens Prisma | Siemens Prisma |
| T1 (structural) parameters | 176 sagittal slices with 1 mm isotropic voxels (TR = 2,530 ms, TE = 3.3 ms, TI = 900 ms, flip = 9 degrees). | 208 sagittal slices with 0.85 mm isotropic voxels (TR = 1,800 ms, TE = 2.37 ms, TI = 900 ms, flip = 8 degrees) | 208 sagittal slices with 0.85 mm isotropic voxels (TR = 1,800 ms, TE = 2.37 ms, TI = 900 ms, flip = 8 degrees) |
| T2* (functional) parameters | EPI sequence with a 90° flip angle and using GRAPPA with an acceleration factor of 2; with the following parameters: thirty-three 4 mm thick near-axial slices acquired in an interleaved order (with 10% distance factor), with an in-plane resolution of 2.1 mm × 2.1 mm, FoV in the phase encoding (A >> P) direction 200 mm and matrix size 96 × 96, TR = 2000 ms and TE = 30 ms | SMS EPI sequence with a 90° flip angle and using a slice acceleration factor of 2, with the following acquisition parameters: fifty-two 2 mm thick near-axial slices acquired in the interleaved order (with 10% distance factor), 2 mm × 2 mm in-plane resolution, FoV in the phase encoding (A >> P) direction 208 mm and matrix size 104 × 104, TR = 2,000 ms, TE = 30 ms, and partial Fourier of 7/8 | SMS EPI sequence with a 90° flip angle and using a slice acceleration factor of 3, with the following acquisition parameters: seventy-two 2 mm thick near-axial slices acquired in the interleaved order (with 10% distance factor), 2 mm × 2 mm in-plane resolution, FoV in the phase encoding (F >> H) direction 208 mm and matrix size 104 × 104, TR = 2,000 ms, TE = 30 ms, and partial Fourier of 7/8 |
| Number of runs | 42,277 | 8,753 | 5,330 |

fMRI data were analyzed using SPM12 (release 7487), CONN EvLab module (release 19b), and other custom MATLAB scripts. Each participant’s functional and structural data were converted from DICOM to NIFTI format. All functional scans were coregistered and resampled using the 4th degree B-spline interpolation to the first scan of the first session (157). Potential outlier scans were identified from the resulting subject-motion estimates as well as from BOLD signal indicators using default thresholds in CONN preprocessing pipeline (5 standard deviations above the mean in global BOLD signal change, or framewise displacement values above 0.9 mm; (158)). Functional and structural data were independently normalized into a common space (the Montreal Neurological Institute [MNI] template; IXI549Space) using SPM12 unified segmentation and normalization procedure (159) with a reference functional image computed as the mean functional data after realignment across all timepoints omitting outlier scans. The output data were resampled to a common bounding box between MNI-space coordinates (-90, -126, -72) and (90, 90, 108), using 2mm isotropic voxels and 4th degree B-spline interpolation for the functional data, and 1mm isotropic voxels and trilinear interpolation for the structural data. BOLD signal in each voxel of each run was then denoised by residualizing against variables controlling for the effect of slow linear drifts, subject-motion parameters, signals originating in white matter and cerebrospinal fluid, and potential outlier scans. Last, the functional data were smoothed spatially using spatial convolution with a 4 mm FWHM Gaussian kernel.

### Tasks and Materials

The task fMRI runs in our dataset come from a large and diverse set of published and unpublished experiments. Since most of these tasks are not of interest (and documenting them fully would amount to an exhaustive history of research conducted in the Fedorenko lab), here we restrict our presentation to a subset of key tasks and materials: those that we directly use in this study either to compute task contrasts or to select or exclude runs for parcellation. These tasks were selected because they have been used relatively frequently and cover a spectrum of targeted cognitive processes (e.g., language, social cognition, and high-level vision). In our Stanford Digital Repository dataset (**URL**), we provide metadata on which of the following experimental tasks apply to which fMRI sessions.

In general, a given participant did not complete all of these tasks, so we only evaluated participants with respect to tasks they did complete. When possible, we used task contrasts derived from the same session used for parcellation. Otherwise, if a task contrast was available in a different session, we used those data (this is why identical task maps sometimes recur in participants with repeated scans in **Fig S1**). If no task contrast was available for a participant, they were omitted from evaluation on that contrast. For some tasks, our dataset includes minor variants in parameters like timing, item lists, and the presence of additional conditions not used in these analyses. For ease of comparison between task types, here we treat these variants as equivalent.

High-level descriptions of the key tasks used in this study are as follows (readers are referred to the citations for additional motivation and methods detail).

#### Language (reading)

Participants read sequences presented one word at a time on a screen in two conditions: a *sentences* condition in which the sequence formed a meaningful sentence, and a perceptually matched *nonwords* condition in which the sequence formed a list of pronounceable pseudowords (31).

#### Language (Listening)

Participants listened to audio recordings in two conditions: *intelligible* speech in a familiar language, and *unintelligible* degraded versions of these recordings (41).

#### Theory of Mind (Verbal)

Participants read vignettes presented one at a time in two conditions: a *false belief* condition and a *false photo* condition (95). The false belief items described a scenario in which a character has a false belief. The false photo items described previous states of the world that are no longer true (e.g., an object that has since been removed). Although careful attention was paid in the design of these items to match the conditions on multiple linguistic dimensions, subsequent work has found that the false belief items are higher in multiple measures of linguistic complexity that are known to modulate language regions, and thus that language network responses to this task do not necessarily reflect theory of mind processing (80).

#### Theory of Mind (Nonverbal)

Participants viewed a silent animated film (the Pixar short *Partly Cloudy*) which has been proposed as a naturalistic theory of mind localizer (160). The film was coded for time intervals that depict (a) mental state content, (b) physical events, (c) non-mentalizing social interactions, and (d) characters experiencing physical pain, and these interval codes were used as regressors to estimate the task contrast (response to *mental* segments over *physical* segments).

#### Executive (Working Memory)

Participants viewed sequences of marked squares on a grid and were tasked with keeping track of all marked locations, assessed at the end of the sequence using a two-alternative forced choice (98). Items occurred in two conditions: easy (4 locations) and hard (8 locations).

#### Numerical (Math)

Participants saw a number and were tasked with computing its cumulative sum with a sequence of three additional numbers (98). The addends occurred in two conditions: easy (magnitude 2-4) and hard (magnitude 6-8).

#### Music

Participants listened to tonal melodies in two conditions, an *intact* condition in which the melody followed typical Western tone sequences and rhythm, and a *scrambled* condition in which these melodies’ tones and rhythms were randomly scrambled (116).

#### High-Level Visual Perception

Participants viewed 3-second silent video clips in 5 conditions: faces, body parts, objects, scenes, and randomly scrambled objects (161).

### fMRI Session Selection

We parcellated every admissible fMRI scan of neuroanatomically typical individuals in the history of the Fedorenko lab as of August 27th, 2024. Inclusion was defined as generously as was feasible. A small percentage of sessions (26 out of 1,983) were excluded for the following reasons:

1. There were missing or corrupt data or metadata, preventing timecourse extraction (11 sessions).
2. There were no functional runs from an all-anatomical scan (1 session).
3. There were no functional runs remaining after exclusion of language localizer runs (14 sessions). This situation arose because we have a privileged interest in evaluating the parcellation against an extensively studied reading-based language localizer (sentences vs. nonword lists) contrast—which is often used to identify the language network (10, 31)—and are following a design in which runs from the localizer task are entirely held out from parcellation to ensure the independence of this critical task. This requirement is conservative and not strictly necessary; in fact, we found that LangFC is recovered from reading localizer runs even when the task structure is regressed out (**SI C**). Runs for the other experimental tasks we examined (described in **Tasks and Materials** below, e.g., the hard vs. easy contrast from the spatial working memory task) were not held out from parcellation, because holding out all contrasts requires either (a) substantial loss of source data for parcellation if removing all evaluation tasks simultaneously, since the evaluation tasks make up a large portion of the available data, or (b) re-parcellating the brain based on different subsets of data for each contrast, which is infeasible (given that we are searching over many parcellation granularities in a large dataset, each complete set of parcellations takes weeks on our compute cluster and requires terabytes of storage).

The remaining dataset contains 1,957 sessions—each typically containing about an hour (1,854±12SEM TRs, TR=2s) of fMRI data—recorded from a total participant population of 1,199 individuals (the difference between the number of sessions and the number of individuals is due to repeated scanning of some individuals). To avoid biasing results, no additional quality controls or filters were applied. Each session’s data was parcellated separately, even when the participant had completed other sessions. This allowed us to assess the stability of parcellations across sessions within an individual (see **LangFC Is Stable Within Individuals and Variable Between Them**) and ensured that our results were representative of the expected performance of our proposed method for a typical cognitive neuroscience experiment (in which each participant may only have one scanning session).

### Parcellation and Evaluation Procedure

We identified and functionally evaluated putative language regions using a five-step procedure within each session of fMRI data in our study: sampling, alignment, labeling, aggregation, and evaluation. The essence of this procedure is schematized in **Fig 1**. Our general purpose codebase for volumetric parcellation of fMRI timecourses in NIFTI format, largely built on the *nilearn* (162) and *scikit learn* (163) Python packages, is available open source: https://github.com/coryshain/parcellate. We have additionally released a separate reproduction codebase for all analyses specific to this study: https://github.com/coryshain/langlocFC. Although many of the design choices described below are motivated by considerations of performance, efficiency, or precedent set by prior work, we found in direct comparisons that most of them had little impact on results, supporting the robustness of the parcellations to diverse implementation details (**SI D**).

We have released all parcellation configuration files and results in the Stanford Digital Repository (**URL**), along with the preprocessed timecourses that were used as input to the parcellation.

#### Sampling

Gray matter voxels from all available functional runs after exclusions (see **fMRI Session Selection**) were individually bandpassed between 0.01 and 0.1Hz, detrended, and z-scored, voxelwise (64). The resulting timecourses were concatenated into a single sequence, and the time dimension was reduced to 200 principal components (PCs). Primarily for computational reasons, the resulting 200-dimensional vector for each of the V voxels was then correlated with the average vector within each of 1,000 parcels in a previously published atlas (53), resulting in a V × 1,000 connectivity matrix (in our case, V=134,713, so the use of parcels for spatial downsampling gives large memory savings without harming results, see **SI 3**). Following prior work (65), the connectivity matrix was then binarized at a quantile threshold of 0.9.

The resulting matrix served as input to an iterative parcellation procedure in which voxels’ connectivity patterns were clustered into *k* networks using *k*-means under 100 random initializations. This both mitigates the sensitivity of *k*-means to initial conditions and provides an opportunity to quantify uncertainty about voxel assignments to networks, in hopes of capturing core-periphery distinctions when relevant (164). This results in a sample of 100 clusterings (discrete parcellations) of the gray matter volume. However, aggregating over this sample is challenging due to the arbitrariness of the cluster labels assigned by *k*-means. Therefore, we include an alignment step, described below.

#### Alignment

We used a greedy approach to align the 100 sampled parcellations into a common set of networks. We first sorted the samples in ascending order (best to worst) according to the mean squared error of their cluster centroids to the data (a measure of clustering quality). We then initialized the probabilistic parcellation using the best sample. For each subsequent sample, we found the optimal spatial alignment (measured as Pearson correlation over the gray matter volume) to the current state of the probabilistic parcellation using linear sum assignment (78) and updated the probabilistic parcellation using a moving average with the newly aligned sample. The result is a set of *k* probabilistic (and potentially spatially overlapping) assignments of voxels to networks, with 0 indicating that a voxel was never assigned to the network and 1 indicating that the voxel was always assigned to the network. Each network in the final probabilistic parcellation was then min-max normalized.

#### Labeling

Interpretable labels were heuristically assigned to a subset of the networks in the probabilistic parcellation based on spatial similarity (Pearson correlation in the gray matter volume) to a set of (non-individualized) reference atlases (65, 79). For each reference atlas, the most spatially similar individualized network in the parcellation was selected and assigned the corresponding functional label (e.g., LANG, DN-B, FPN-A, etc.). The assignments could be many-to-one, in that the same network could be chosen by multiple reference atlases (in practice, for sufficiently granular parcellations, this was rare).

#### Aggregation

We wanted both to explore the influence of the choice of the number of networks *k* and allow *k* to vary as needed between individuals. Thus, for this study, we placed all three of the preceding steps (sampling, alignment, and labeling) under a loop over values of *k* from 10 to 200 in increments of 10, resulting in 20 parcellations (at different *k*) per scanning session. We defined the optimal *k* for a session to be the value that maximized the average spatial similarity between the reference atlases and their corresponding labeled iFC counterparts. We chose to select on correspondence to the reference atlases in order to avoid any bias toward a particular pattern of functional tuning (as would be the case if we e.g., selected on similarity to language task responses). In order to avoid *post hoc* selection of results, we refitted the parcellation using the chosen value of *k* to produce our final parcellation for the participant.

#### Evaluation

As described in **Results**, we applied two key analyses of functional tuning to all labeled networks in the parcellation, for all available tasks within our critical set for which we had data for the participant: z(r), a measure of spatial similarity between the network and the distribution of task responses, and mean *t*-value, a weighted sum across the gray matter volume of the voxelwise *t*-values for a given contrast, weighted by their iFC-estimated probability of belonging to the network in question.

### Additional Analyses

Here we describe all remaining methods used for secondary analyses in this study.

#### LangFC Periphery (Fig 2C)

To identify the LangFC “periphery” (visualized at the group level in **Fig 2C**), we developed a procedure for finding a union of subnetworks in our probabilistic parcellation that maximized overall spatial similarity (Pearson correlation) to the reference atlas. To do so, in each individual, we started with LangFC labeled relative to a reference atlas (LanA), and we used a greedy approach in which we iteratively identified the next most similar network to the reference atlas and added it to the union (by definition, the first of these networks is LangFC) as long as doing so increased spatial similarity between the union and the reference atlas. This greedy approach is computationally efficient at the expense of global optimality, since we did not exhaustively maximize spatial similarity over the power set of networks in the parcellation. Our stopping criterion guarantees that there will always be at least one network (LangFC) in the union, and possibly (but not necessarily) more. A voxel’s probability of belonging to the union is equal to its probability of belonging to at least one of the component subnetworks, that is, the probability of at least one success under a Poisson-binomial distribution (165) parameterized by the membership probabilities for each subnetwork in the union. Union membership was used to create a new probabilistic atlas representing the probability of belonging to an iFC-based extended language network (82). To identify the LangFC “periphery” in an individual, we discretized both LangFC and extended LangFC (p > 0.5) and selected all voxels belonging to extended LangFC but not LangFC. These boolean maps per participant were then averaged across all participants in stereotactic space to produce the probabilistic atlas visualized at the bottom of **Fig 2C**, which provides an iFC-based measure of the topographic tendencies in the language network’s periphery.

#### Nonlinguistic Runs (Fig 3A)

To produce the “nonlinguistic runs” analyses on the right side of **Fig 3A**, we re-parcellated all sessions that contained at least one of a number of tasks that we identified as nonlinguistic, in the sense that no language was present in the stimulus and no overt or covert language response was requested. These tasks included the spatial working memory task, nonverbal theory of mind task, music task, and visual perception tasks described in **Tasks and Materials**. They also included resting state scans, silent movie watching, various non-linguistic visual perception tasks (e.g., motion perception, action perception), abstract symbolic rule induction tasks, and nonspeech auditory perception tasks. We have uploaded a table of all nonlinguistic tasks and brief descriptions to the Stanford Digital Repository (**URL**). We cannot of course rule out the possibility that some participants e.g., engaged in inner speech during the task (166–168), except to note that some of these tasks (spatial working memory and math) occur frequently among our nonlinguistic runs and elicit weak or negative responses in language areas (**Fig 3A**), suggesting that covert language is not used pervasively in order to support these types of thought (24).

### Variability Between and Within Individuals (Fig 4A)

To measure within- and between-individual variation as reported in **Fig 4A**, we used the following procedures.

Between-individual variation was computed by taking both the LangFC topographies on the one hand and the reading localizer task maps on the other from all sessions in our main analysis, averaging these across sessions as needed in cases where the same person participated in multiple sessions. We then computed the between-individual spatial correlation by averaging across the lower triangle of the (1,199 x 1,199) participant-wise Fisher-transformed correlation matrix.

Within-individual variation was computed by finding all participants with at least two sessions (305 participants, 1,063 sessions, 3.5 sessions per participant on average) and computing session-wise Fisher-transformed spatial correlation matrices between LangFC topographies on the one hand and between reading localizer task maps on the other, independently for each individual. The resulting Fisher correlations were then averaged to provide an estimate of typical stability between runs within an an individual.

### Partial Reproduction of Shain et al. (2024)

For reliable comparison to task-based localization in Ref. (102), we exactly matched their procedure for defining functional regions of interest (fROIs) with one change: we replaced the S-N localizer task *t*-maps with LangFC probabilistic maps. The resulting procedure was as follows, for each participant:

1. Mask the brain volume with each of six regional parcels from a prior study (20): left pars orbitalis (LIFGorb), left pars triangularis (LIFGtri), left temporal pole (LTP), left anterior temporal sulcus (aSTS), left posterior temporal sulcus (pSTS), and left temporoparietal junction (LTPJ).
2. Within each regional mask, identify the subset of voxels with the top 10% probability of belonging to LangFC according to that participant’s probabilistic network estimate. This top-10% subset of voxels defines the fROI for each of the six regional masks.
3. Within the fROI, compute the average effect size for each contrast of interest—see Ref. (102) for contrast definitions.

The fact that LangFC defines valid network membership probabilities (unlike e.g., a task *t*-map) offers principled alternatives to the design above, such as taking a weighted average over the entire gray matter volume (as in **Fig 3A**) or selection by absolute thresholding (e.g., *p* > 0.5, i.e., more likely than not to belong to LangFC). Such approaches could have advantages, including the possibility of discovering areas that fall outside the regional masks, or application to participants with different brain anatomy relative to template spaces derived from typical adults (e.g., children or individuals with brain lesions). We leave validation of alternative localization criteria to future work.

## Acknowledgments

We would like to acknowledge the Athinoula A. Martinos Imaging Center at the McGovern Institute for Brain Research at MIT, and its support team (Steve Shannon and Atsushi Takahashi). We are also indebted to the many past and current members of the Fedorenko lab (including undergraduates, research assistants, graduate students, and postdocs) for the countless hours they collectively invested in gathering, preprocessing, and analyzing the data that made this study possible. We would also like to thank Anya Ivanova and Randy Buckner for their valuable input, and Atsushi Takahashi and Alex Fung for help in retrieving acquisition sequence details for this dataset. In addition, EF was supported by NIH awards DC016607 and DC016950 from NIDCD, and NS121471 from NINDS, and research funds from the McGovern Institute for Brain Research, the Department of Brain and Cognitive Sciences, the Simons Center for the Social Brain, and MIT’s Quest for Intelligence.

