## Supplementary Information for "A language network in the individualized functional connectomes of over 1,000 human brains doing arbitrary tasks"

### SI A: Individual Participant Visualizations of LangFC vs. Task Data

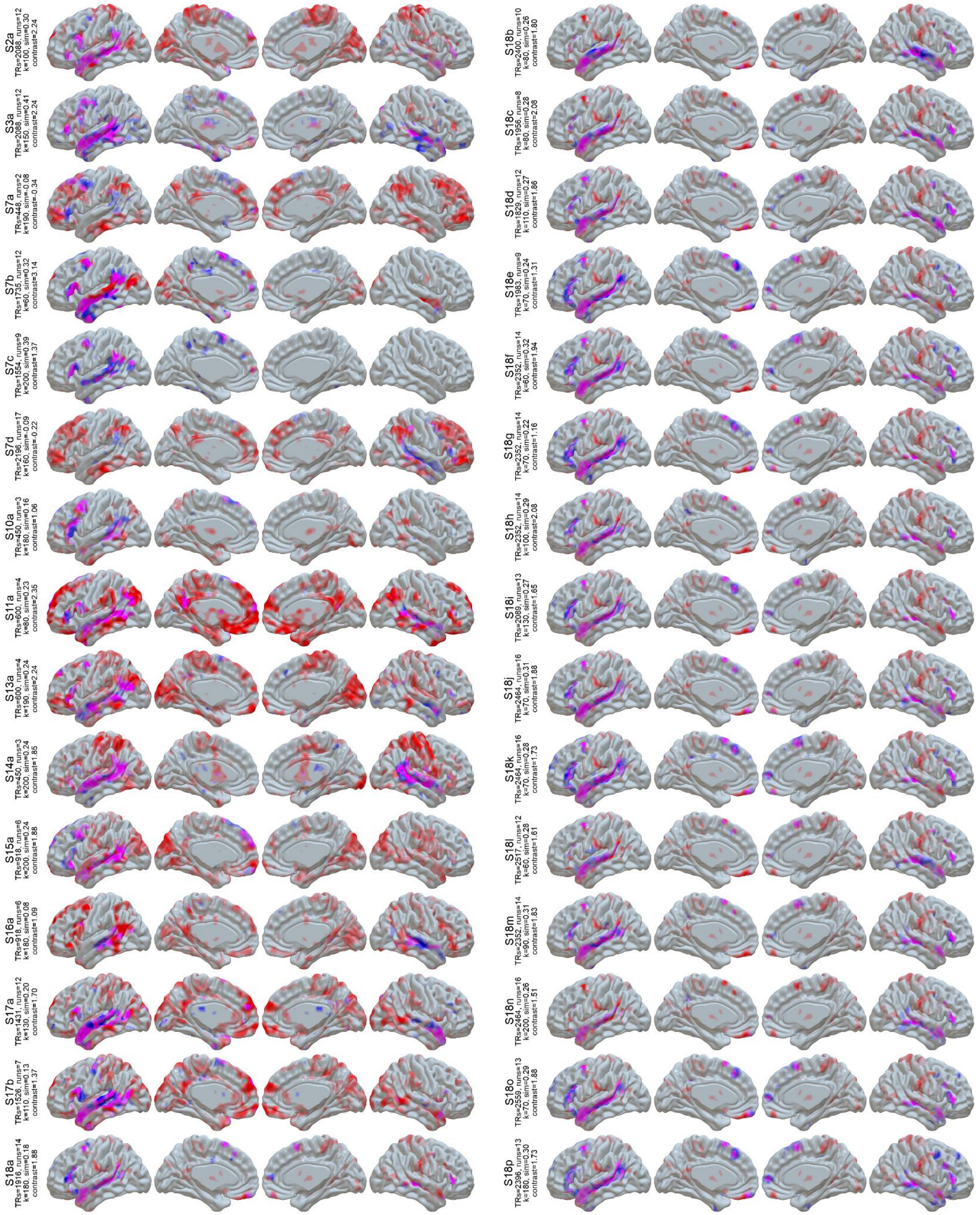


**Figure S1:** LangFC (blue) vs. task *t*-maps (sentences vs. nonword lists or *S-N*, red) in the 10 sessions with the highest S-N contrast stability between even and odd runs. Overlap is shown in magenta, and opacity reflects magnitude ($0.2 < p < 0.8$ for LangFC, $1 < t < 4$ for S-N).

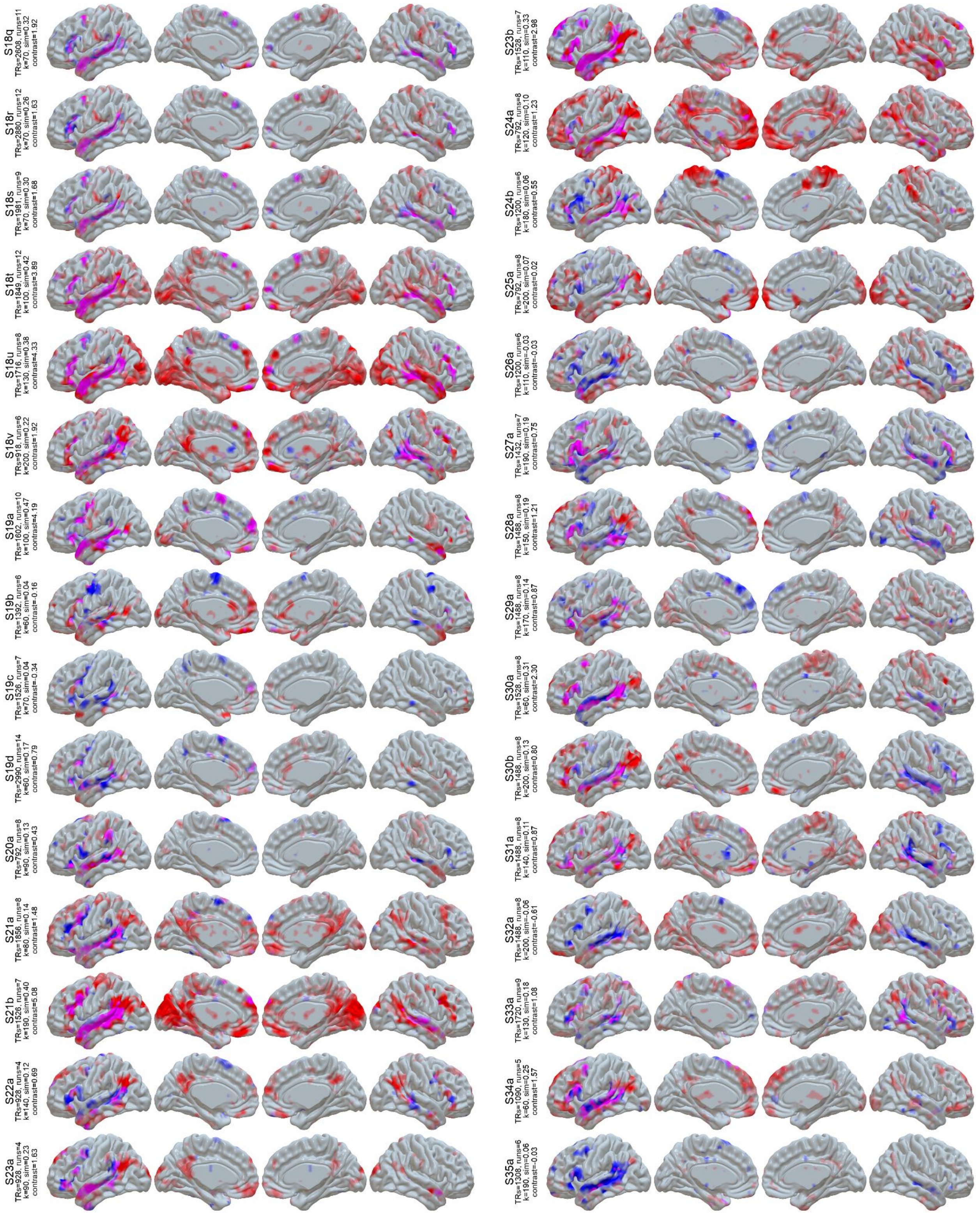


**Figure S1 (cont):** LangFC (blue) vs. task *t*-maps (sentences vs. nonword lists or *S-N*, red) in the 10 sessions with the highest S-N contrast stability between even and odd runs. Overlap is shown in magenta, and opacity reflects magnitude (0.2 < *p* < 0.8 for LangFC, 1 < *t* < 4 for S-N).

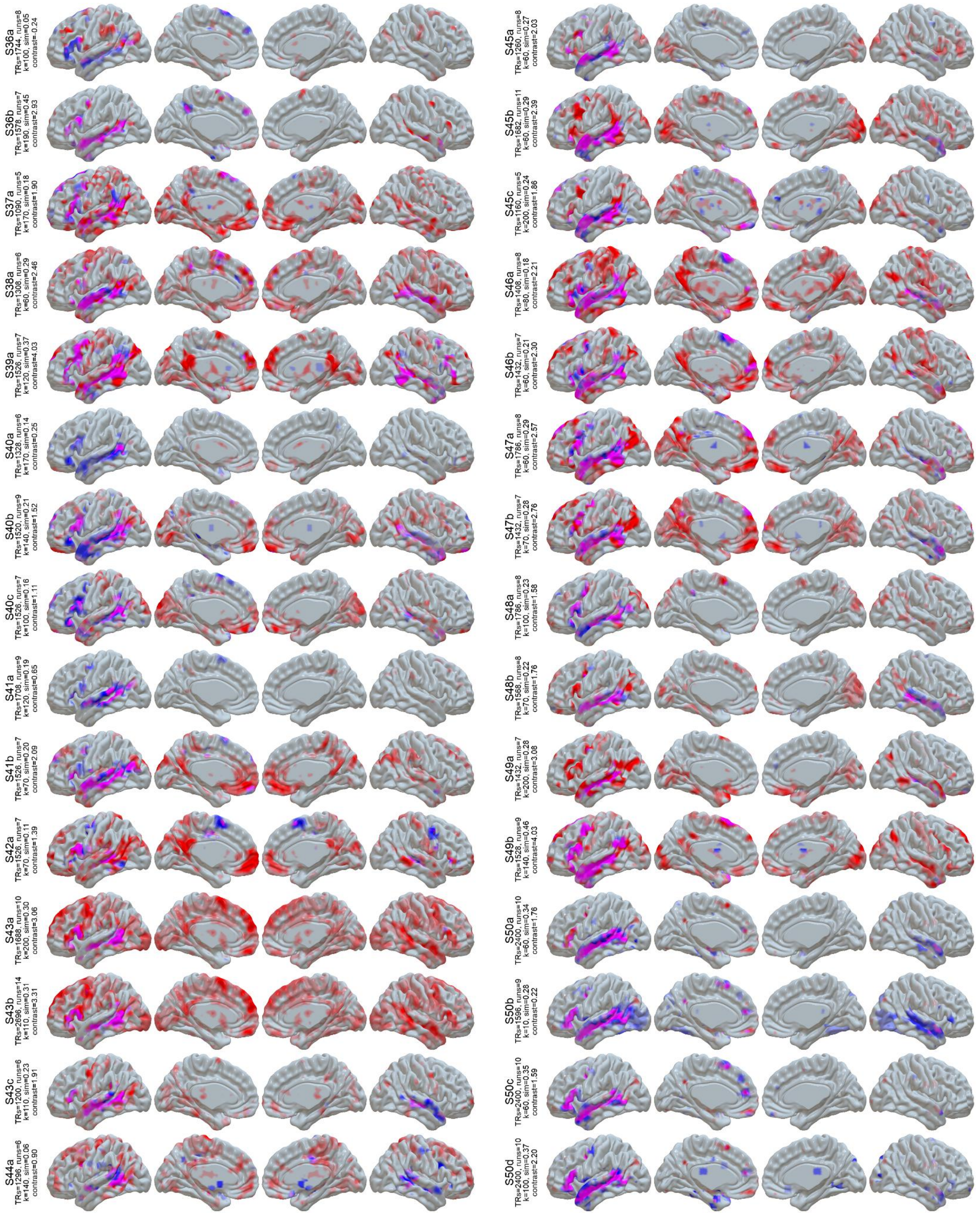


**Figure S1 (cont):** LangFC (blue) vs. task *t*-maps (sentences vs. nonword lists or *S-N*, red) in the 10 sessions with the highest S-N contrast stability between even and odd runs. Overlap is shown in magenta, and opacity reflects magnitude (0.2 < *p* < 0.8 for LangFC, 1 < *t* < 4 for S-N).

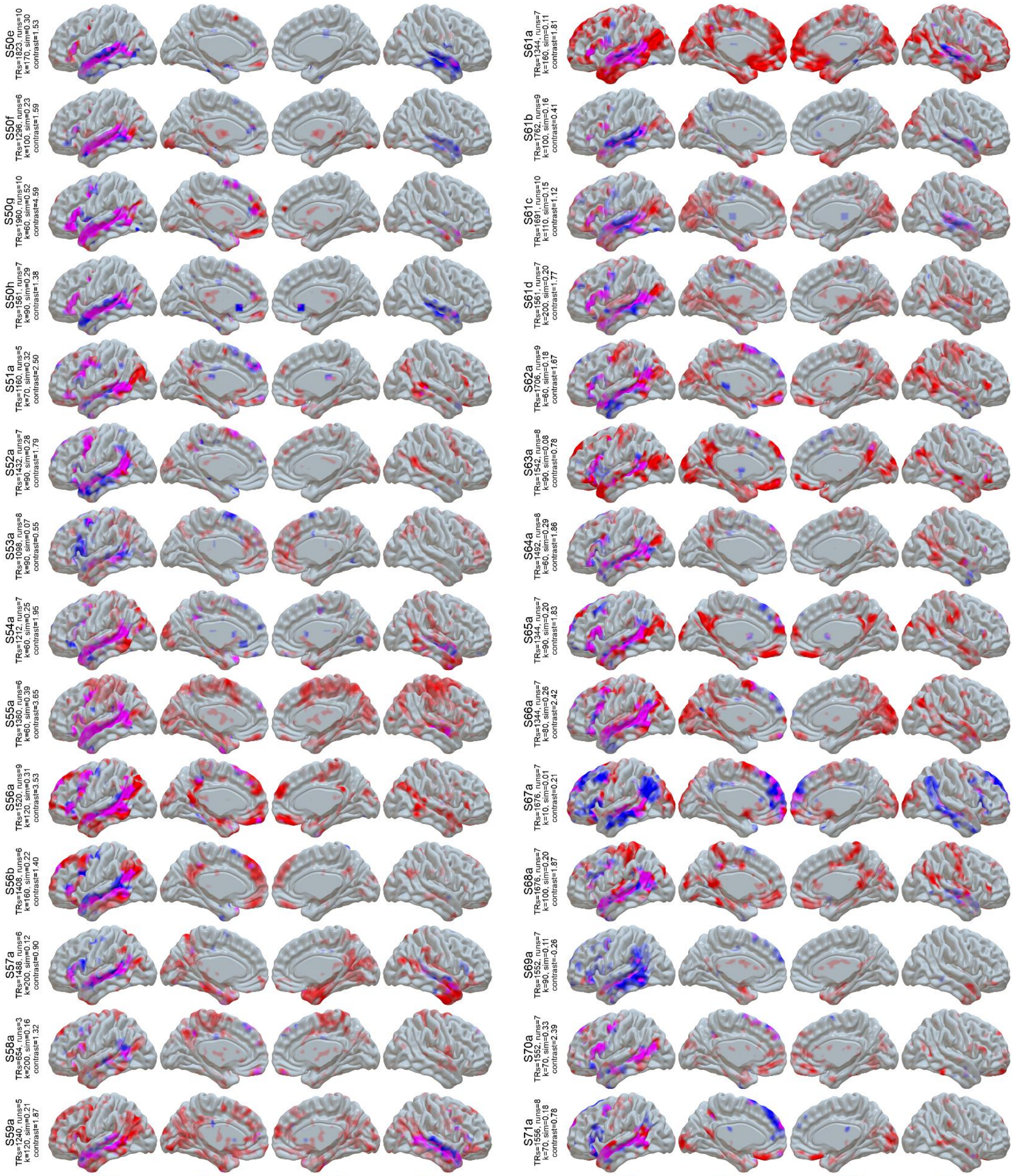


**Figure S1 (cont):** LangFC (blue) vs. task *t*-maps (sentences vs. nonword lists or *S-N*, red) in the 10 sessions with the highest S-N contrast stability between even and odd runs. Overlap is shown in magenta, and opacity reflects magnitude (0.2 < $p$ < 0.8 for LangFC, 1 < $t$ < 4 for S-N).

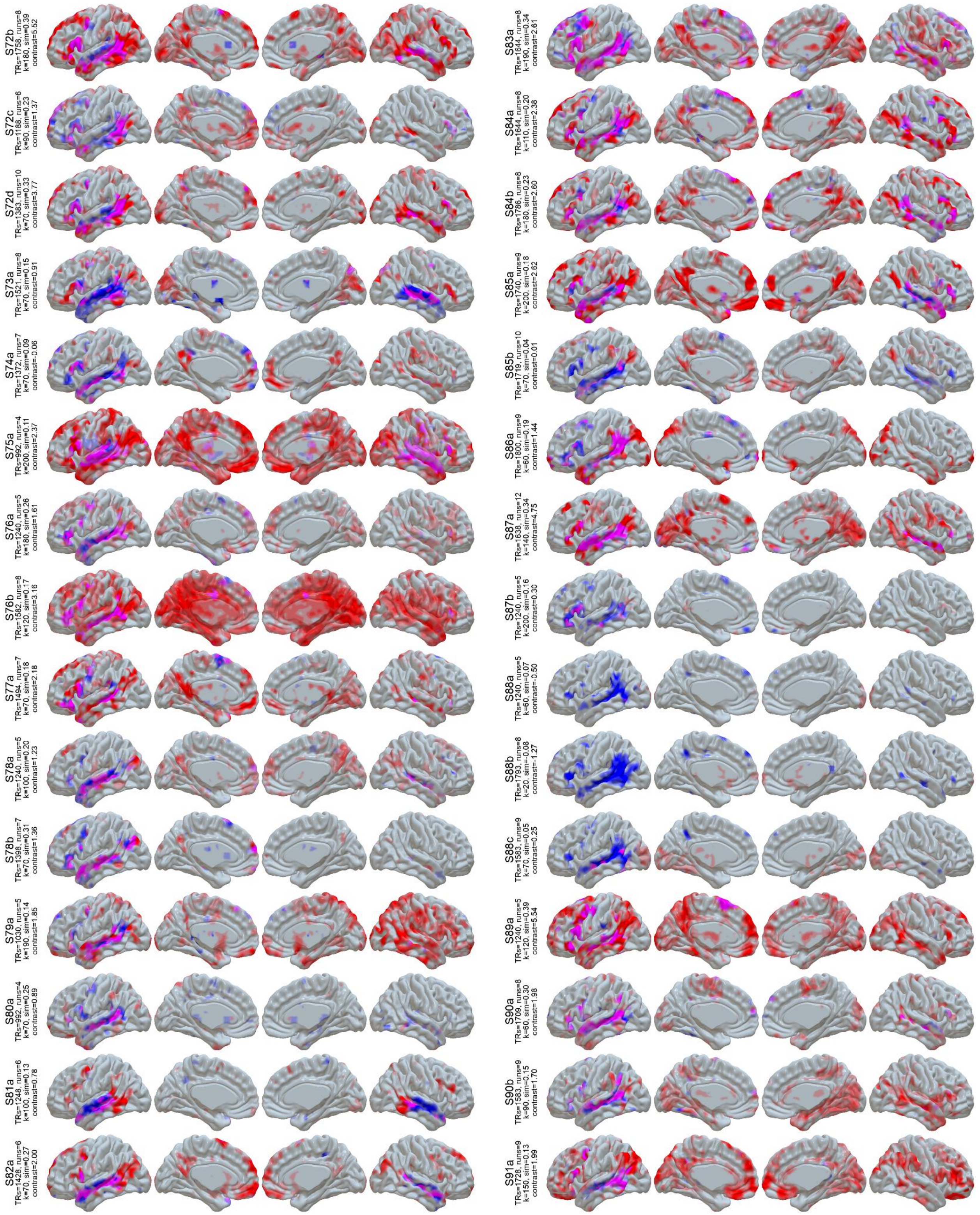


**Figure S1 (cont):** LangFC (blue) vs. task *t*-maps (sentences vs. nonword lists or *S-N*, red) in the 10 sessions with the highest S-N contrast stability between even and odd runs. Overlap is shown in magenta, and opacity reflects magnitude (0.2 < *p* < 0.8 for LangFC, 1 < *t* < 4 for S-N).

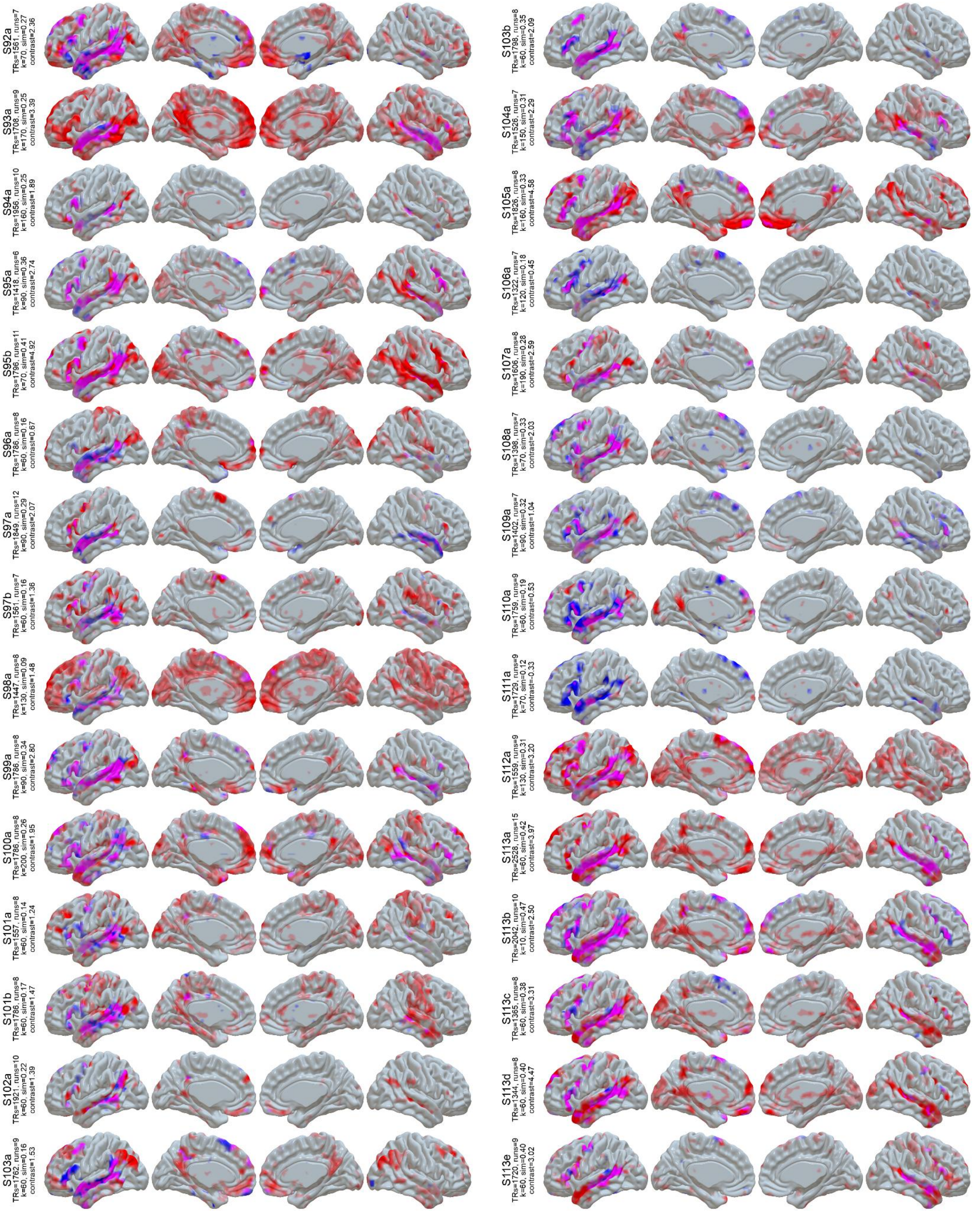


**Figure S1 (cont):** LangFC (blue) vs. task *t*-maps (sentences vs. nonword lists or *S-N*, red) in the 10 sessions with the highest S-N contrast stability between even and odd runs. Overlap is shown in magenta, and opacity reflects magnitude (0.2 < *p* < 0.8 for LangFC, 1 < *t* < 4 for S-N).

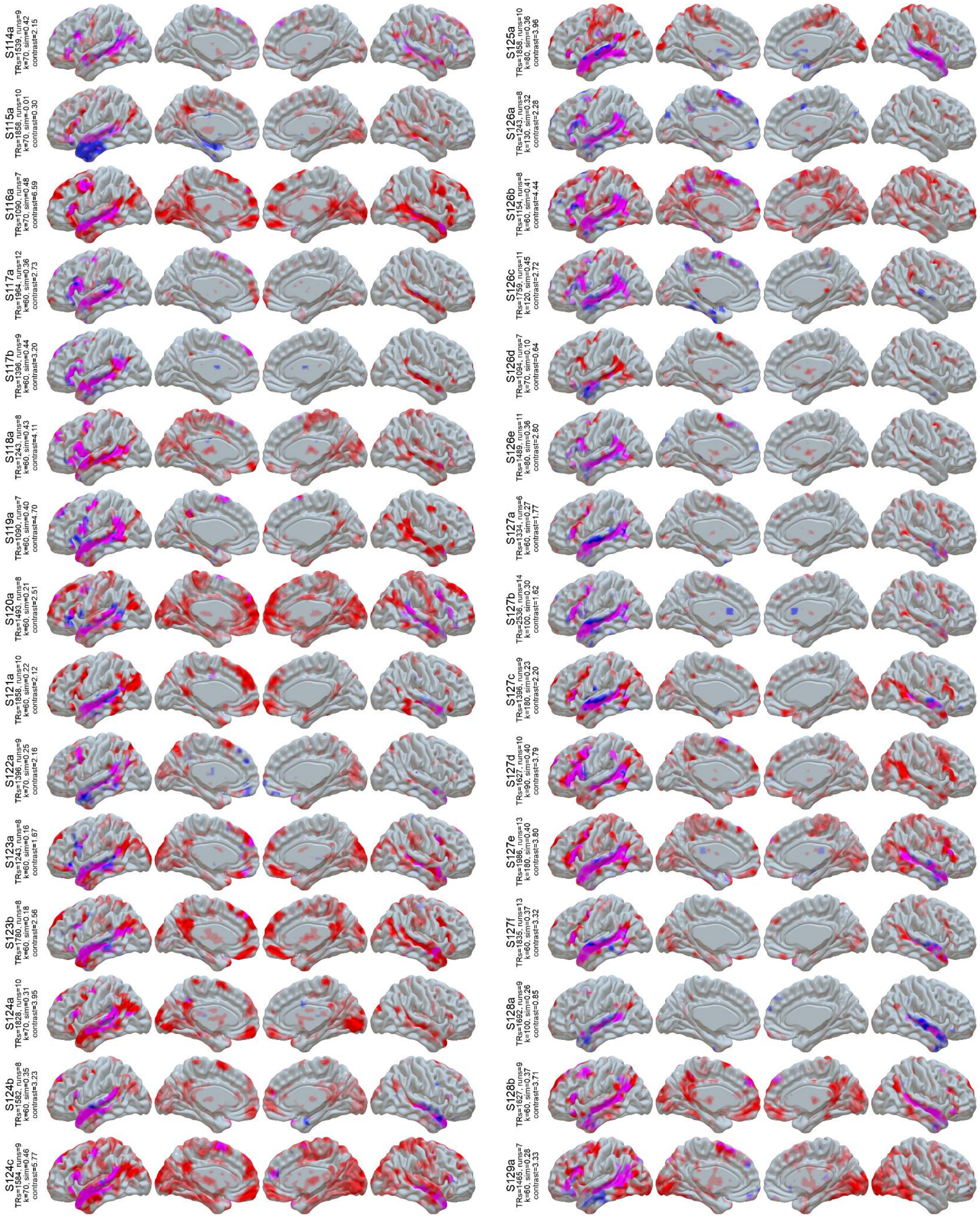


**Figure S1 (cont):** LangFC (blue) vs. task *t*-maps (sentences vs. nonword lists or *S-N*, red) in the 10 sessions with the highest S-N contrast stability between even and odd runs. Overlap is shown in magenta, and opacity reflects magnitude ($0.2 < p < 0.8$ for LangFC, $1 < t < 4$ for S-N).

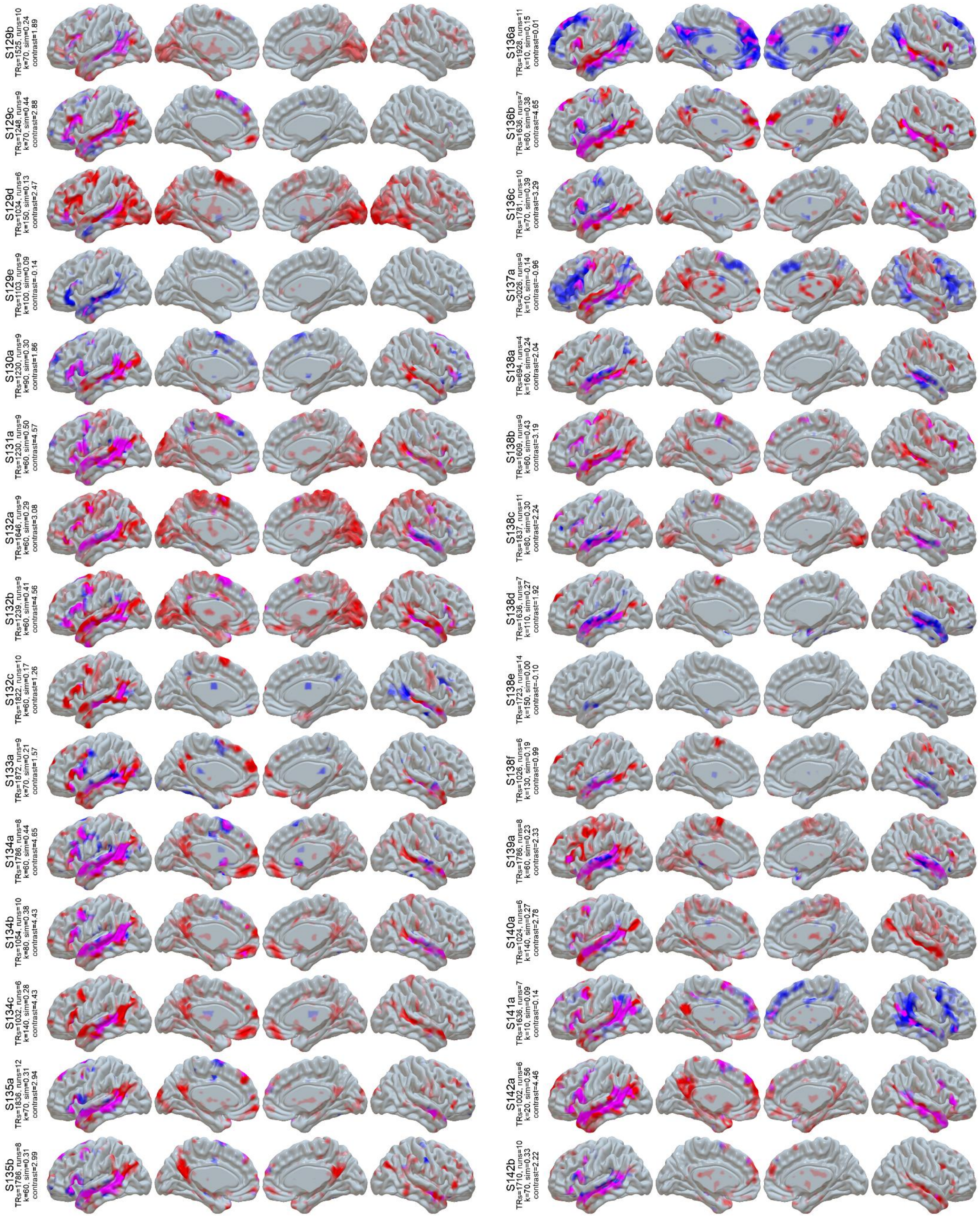


**Figure S1 (cont):** LangFC (blue) vs. task *t*-maps (sentences vs. nonword lists or *S-N*, red) in the 10 sessions with the highest S-N contrast stability between even and odd runs. Overlap is shown in magenta, and opacity reflects magnitude (0.2 < *p* < 0.8 for LangFC, 1 < *t* < 4 for S-N).

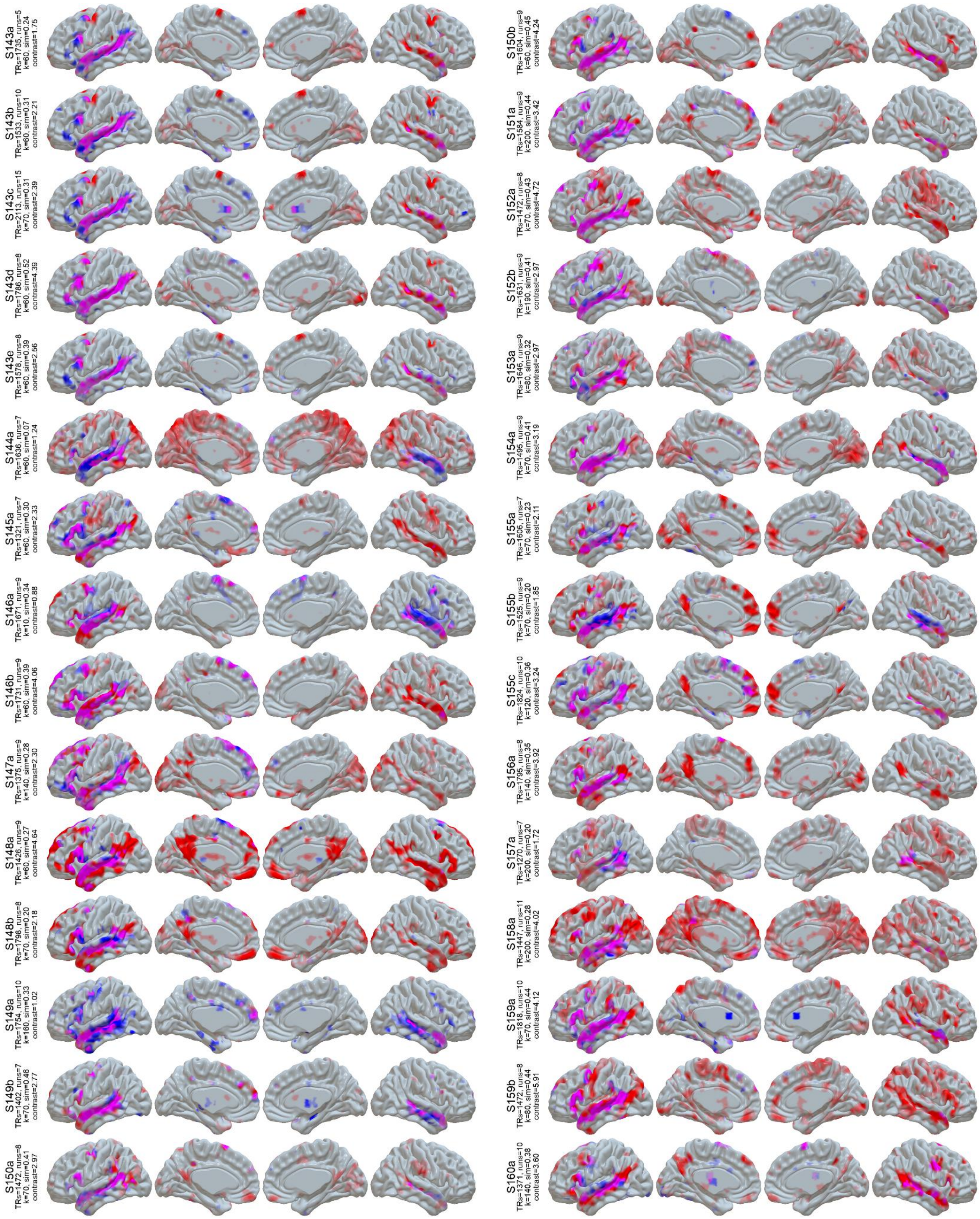


**Figure S1 (cont):** LangFC (blue) vs. task *t*-maps (sentences vs. nonword lists or *S-N*, red) in the 10 sessions with the highest S-N contrast stability between even and odd runs. Overlap is shown in magenta, and opacity reflects magnitude (0.2 < *p* < 0.8 for LangFC, 1 < *t* < 4 for S-N).

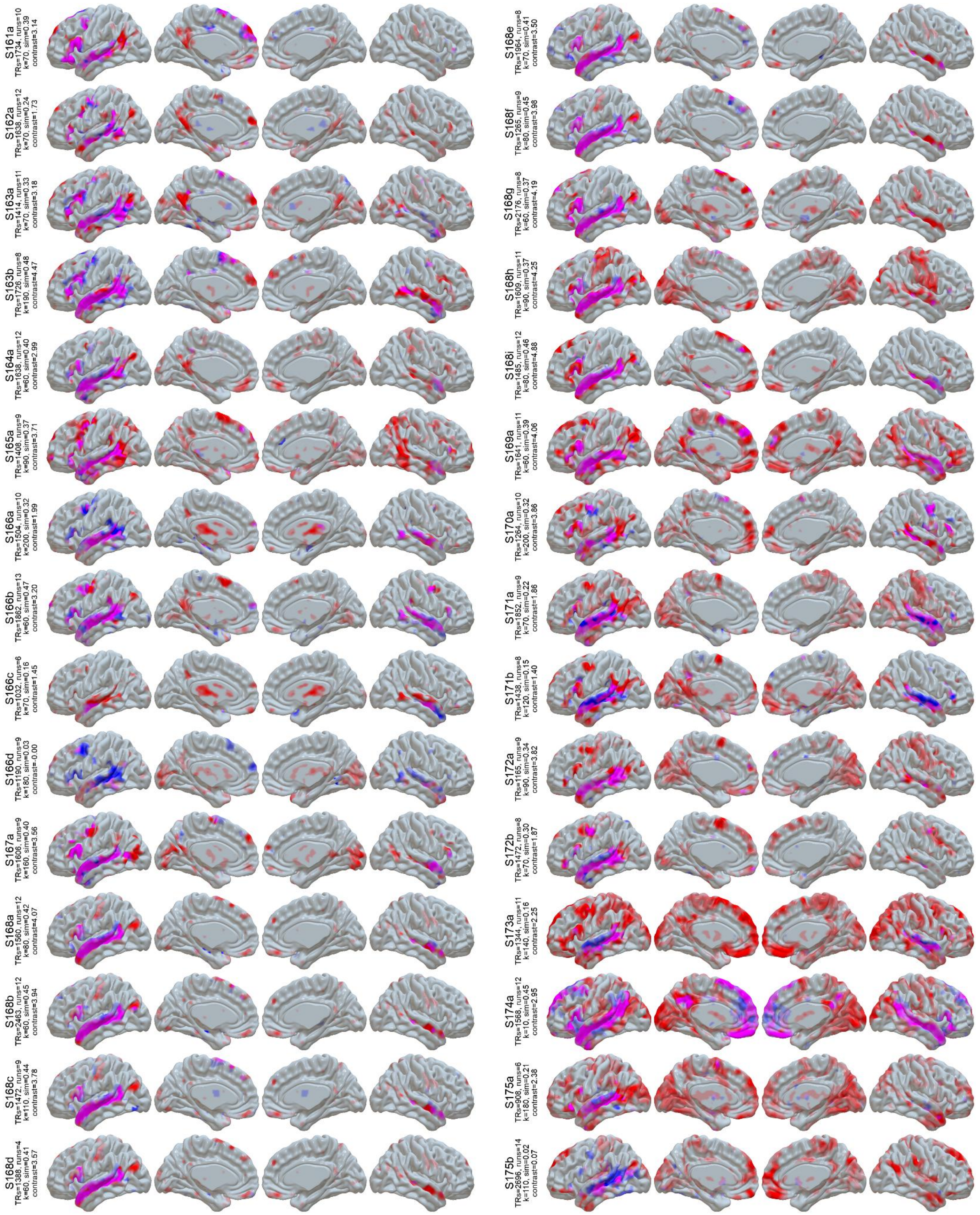


**Figure S1 (cont):** LangFC (blue) vs. task *t*-maps (sentences vs. nonword lists or *S-N*, red) in the 10 sessions with the highest S-N contrast stability between even and odd runs. Overlap is shown in magenta, and opacity reflects magnitude ($0.2 < p < 0.8$ for LangFC, $1 < t < 4$ for S-N).

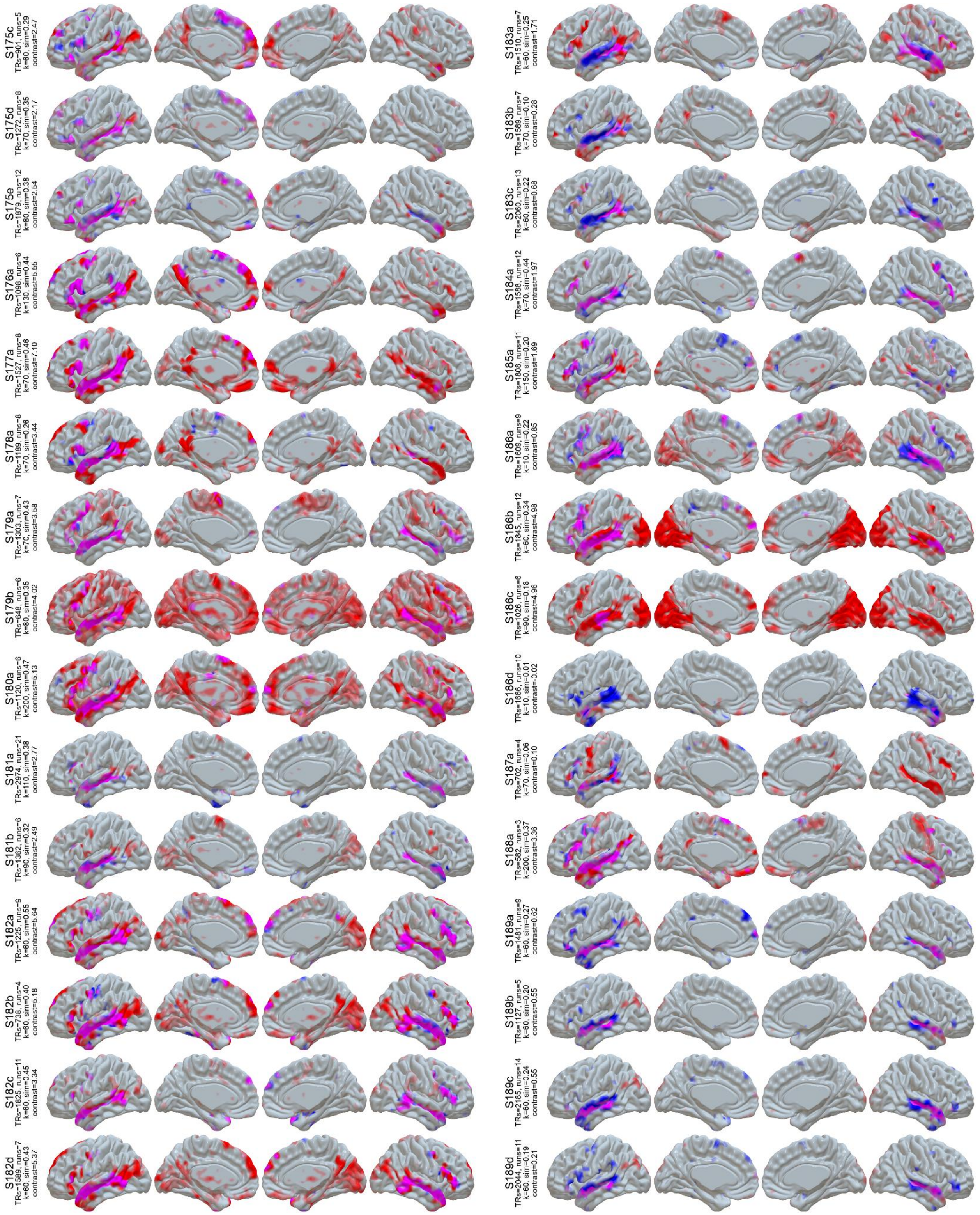

**Figure S1 (cont):** LangFC (blue) vs. task *t*-maps (sentences vs. nonword lists or *S-N*, red) in the 10 sessions with the highest S-N contrast stability between even and odd runs. Overlap is shown in magenta, and opacity reflects magnitude (0.2 < $p$ < 0.8 for LangFC, 1 < $t$ < 4 for S-N).

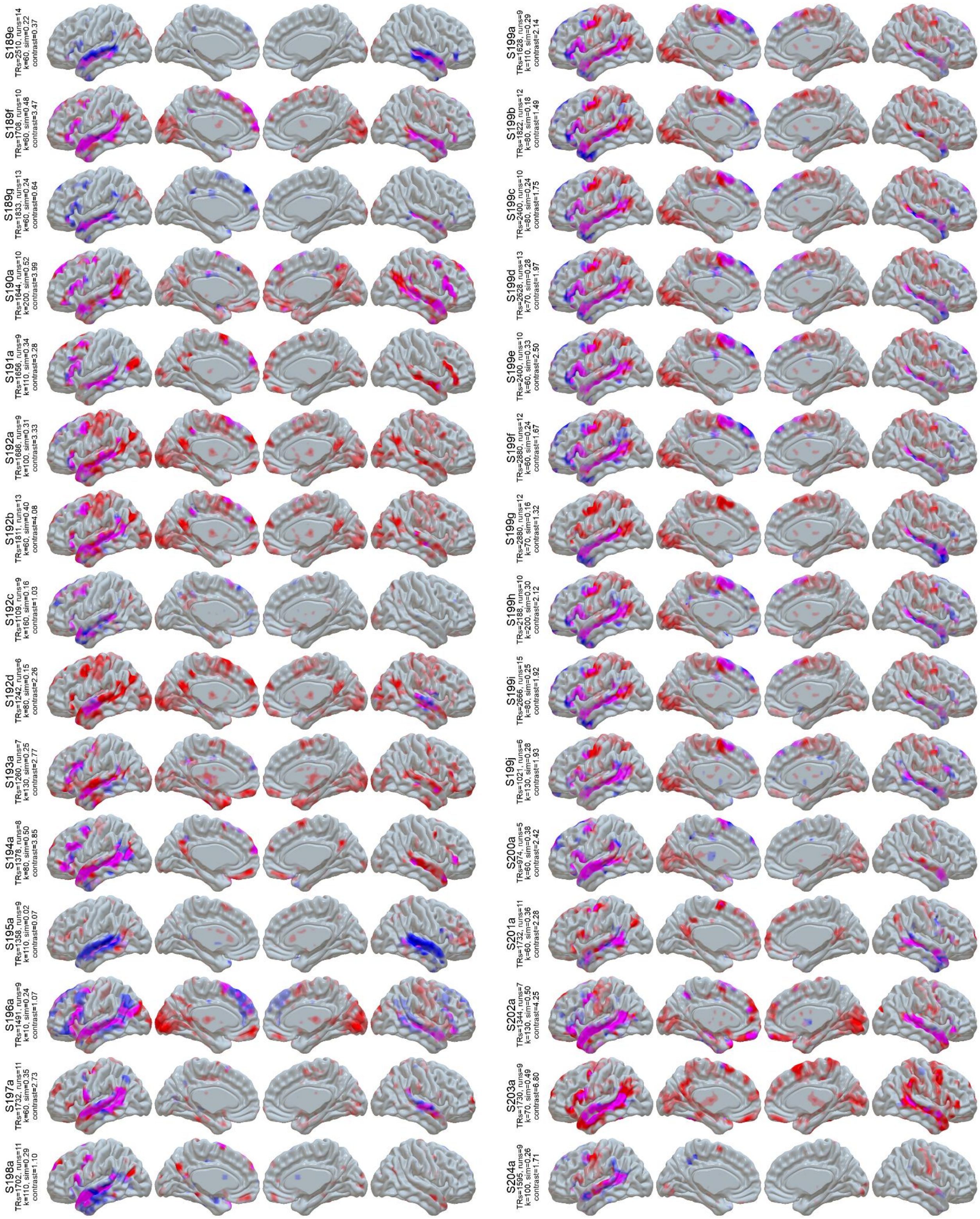


**Figure S1 (cont):** LangFC (blue) vs. task *t*-maps (sentences vs. nonword lists or *S-N*, red) in the 10 sessions with the highest S-N contrast stability between even and odd runs. Overlap is shown in magenta, and opacity reflects magnitude (0.2 < $p$ < 0.8 for LangFC, 1 < $t$ < 4 for S-N).

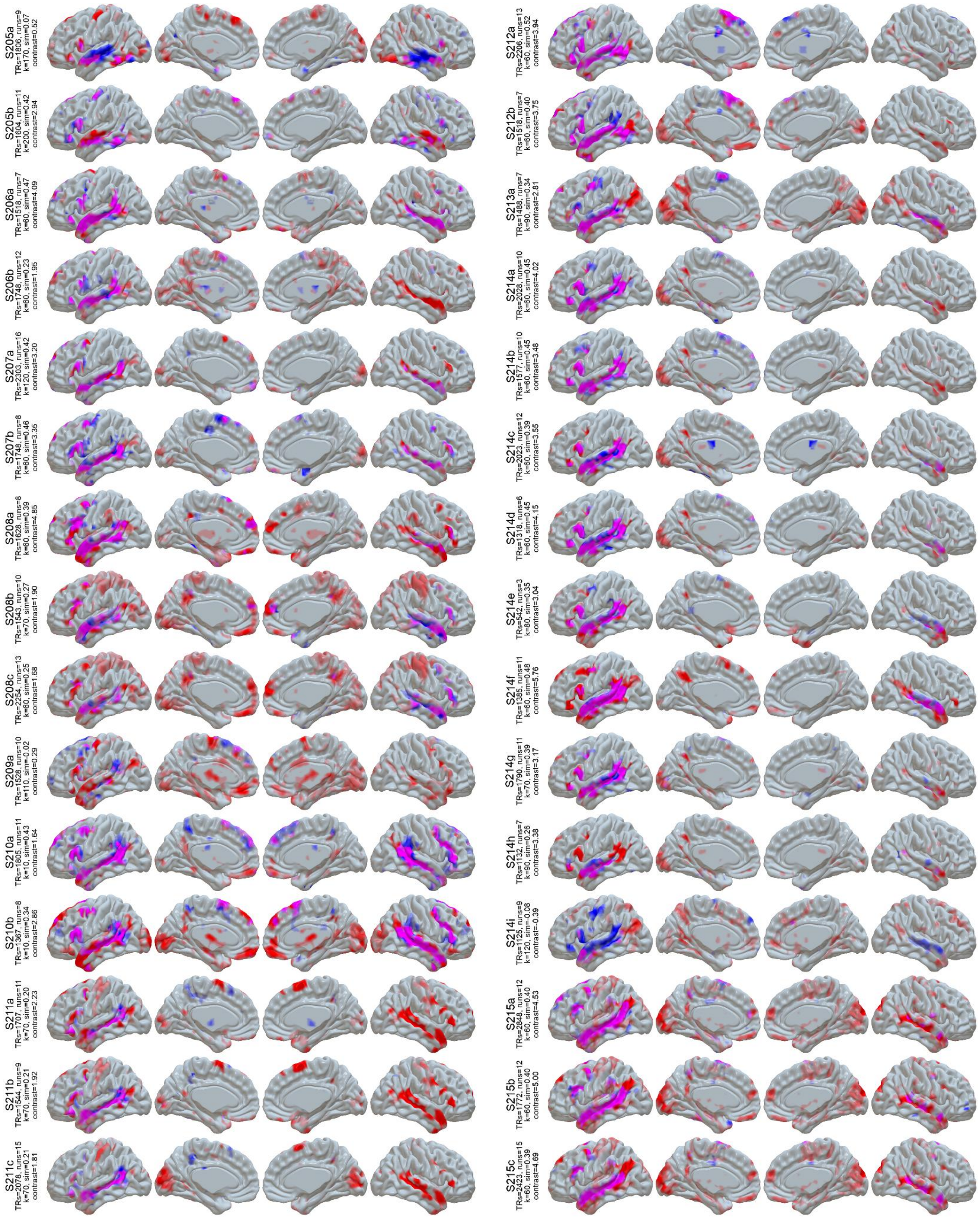


**Figure S1 (cont):** LangFC (blue) vs. task *t*-maps (sentences vs. nonword lists or *S-N*, red) in the 10 sessions with the highest S-N contrast stability between even and odd runs. Overlap is shown in magenta, and opacity reflects magnitude (0.2 < $p$ < 0.8 for LangFC, 1 < $t$ < 4 for S-N).

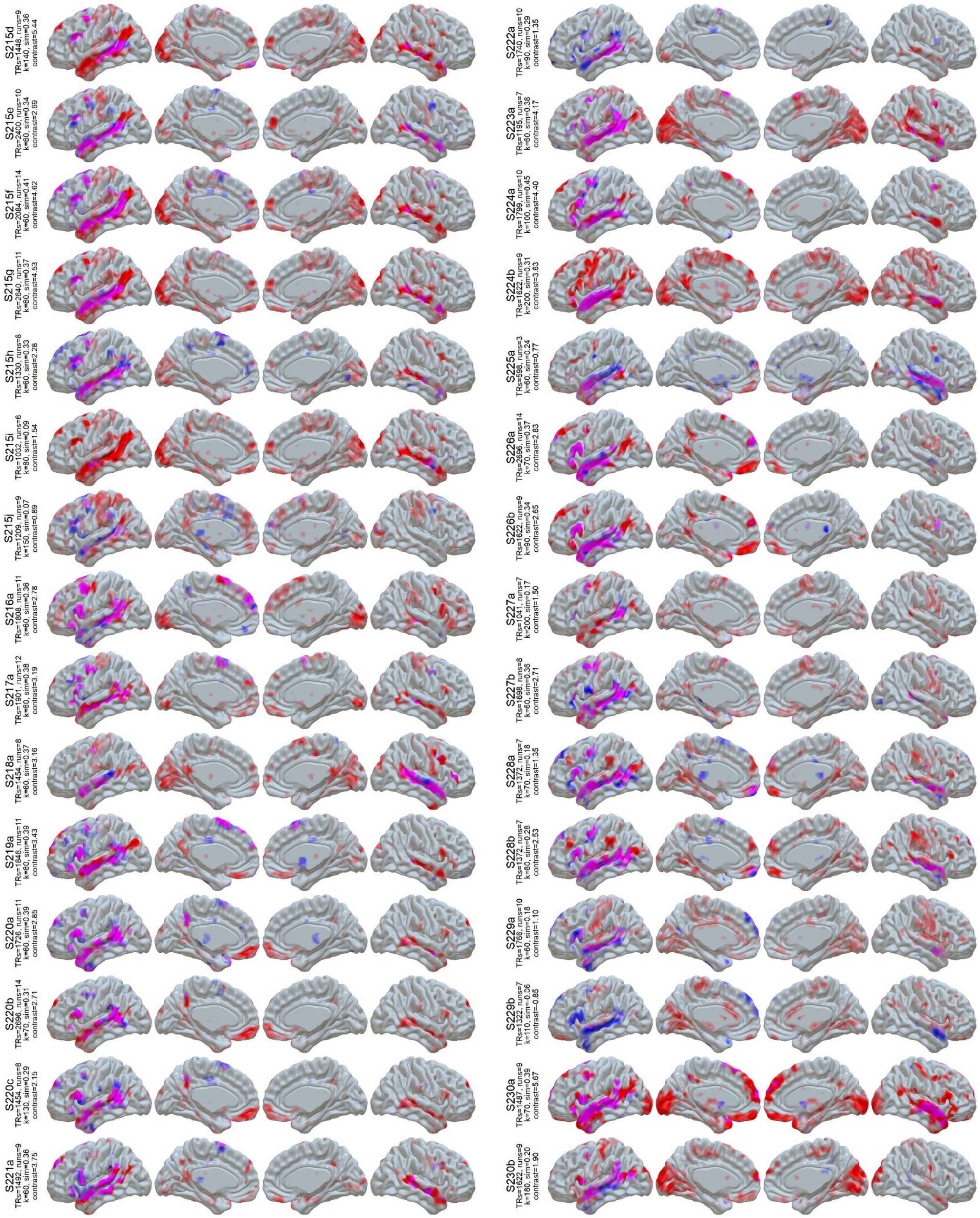

**Figure S1 (cont):** LangFC (blue) vs. task *t*-maps (sentences vs. nonword lists or *S-N*, red) in the 10 sessions with the highest S-N contrast stability between even and odd runs. Overlap is shown in magenta, and opacity reflects magnitude ($0.2 < p < 0.8$ for LangFC, $1 < t < 4$ for S-N).

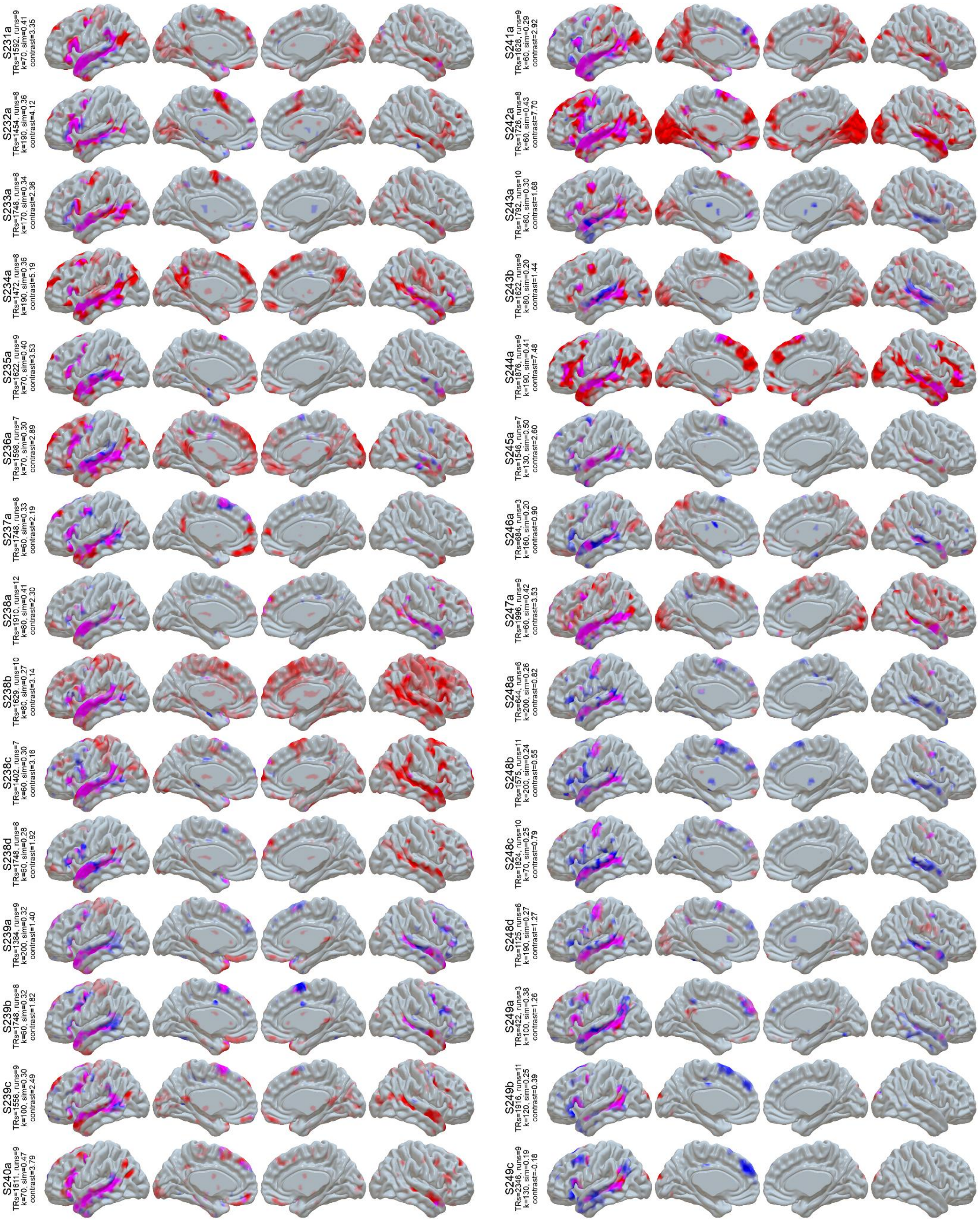


**Figure S1 (cont):** LangFC (blue) vs. task *t*-maps (sentences vs. nonword lists or *S-N*, red) in the 10 sessions with the highest S-N contrast stability between even and odd runs. Overlap is shown in magenta, and opacity reflects magnitude (0.2 < *p* < 0.8 for LangFC, 1 < *t* < 4 for S-N).

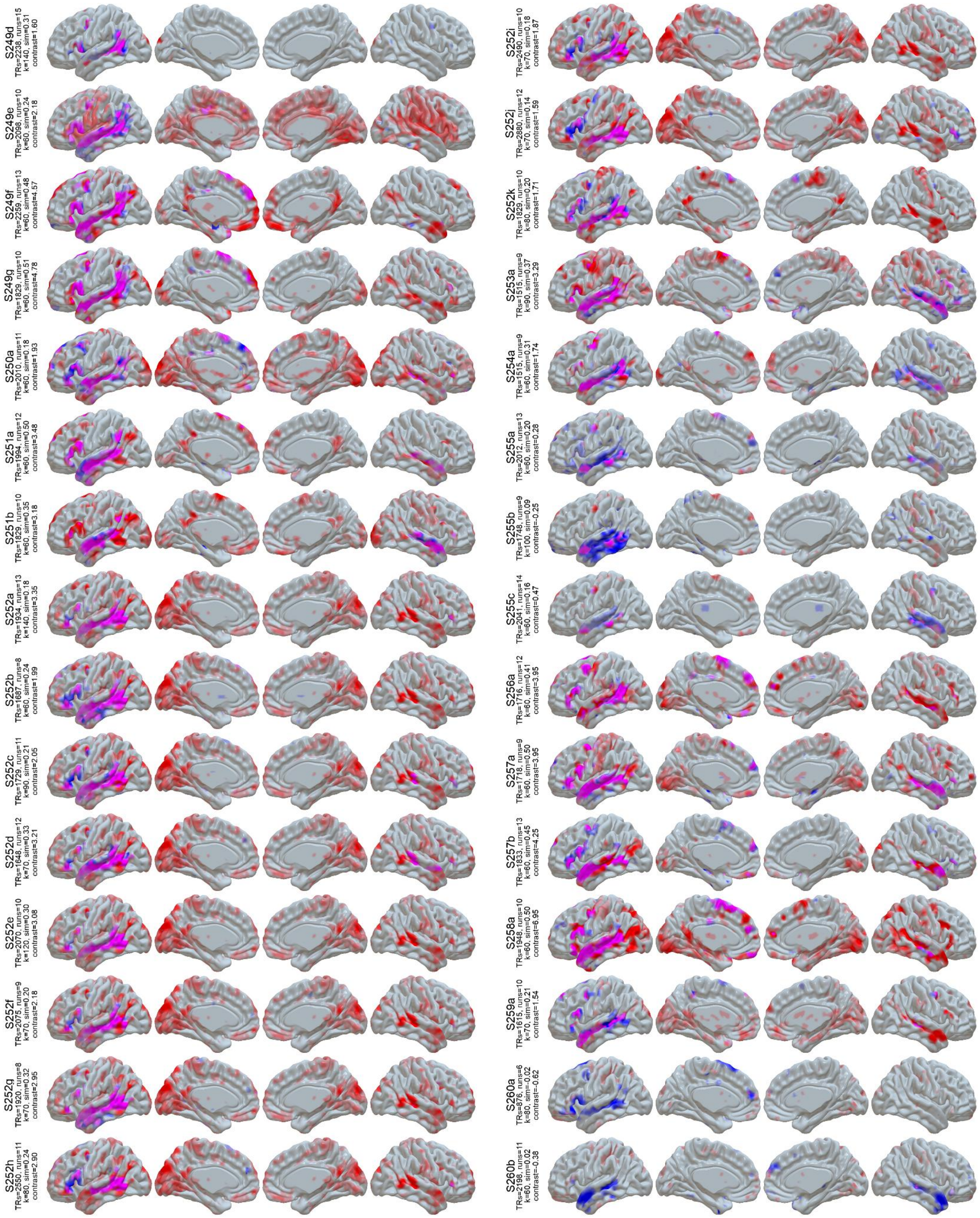


**Figure S1 (cont):** LangFC (blue) vs. task *t*-maps (sentences vs. nonword lists or *S-N*, red) in the 10 sessions with the highest S-N contrast stability between even and odd runs. Overlap is shown in magenta, and opacity reflects magnitude (0.2 < *p* < 0.8 for LangFC, 1 < *t* < 4 for S-N).

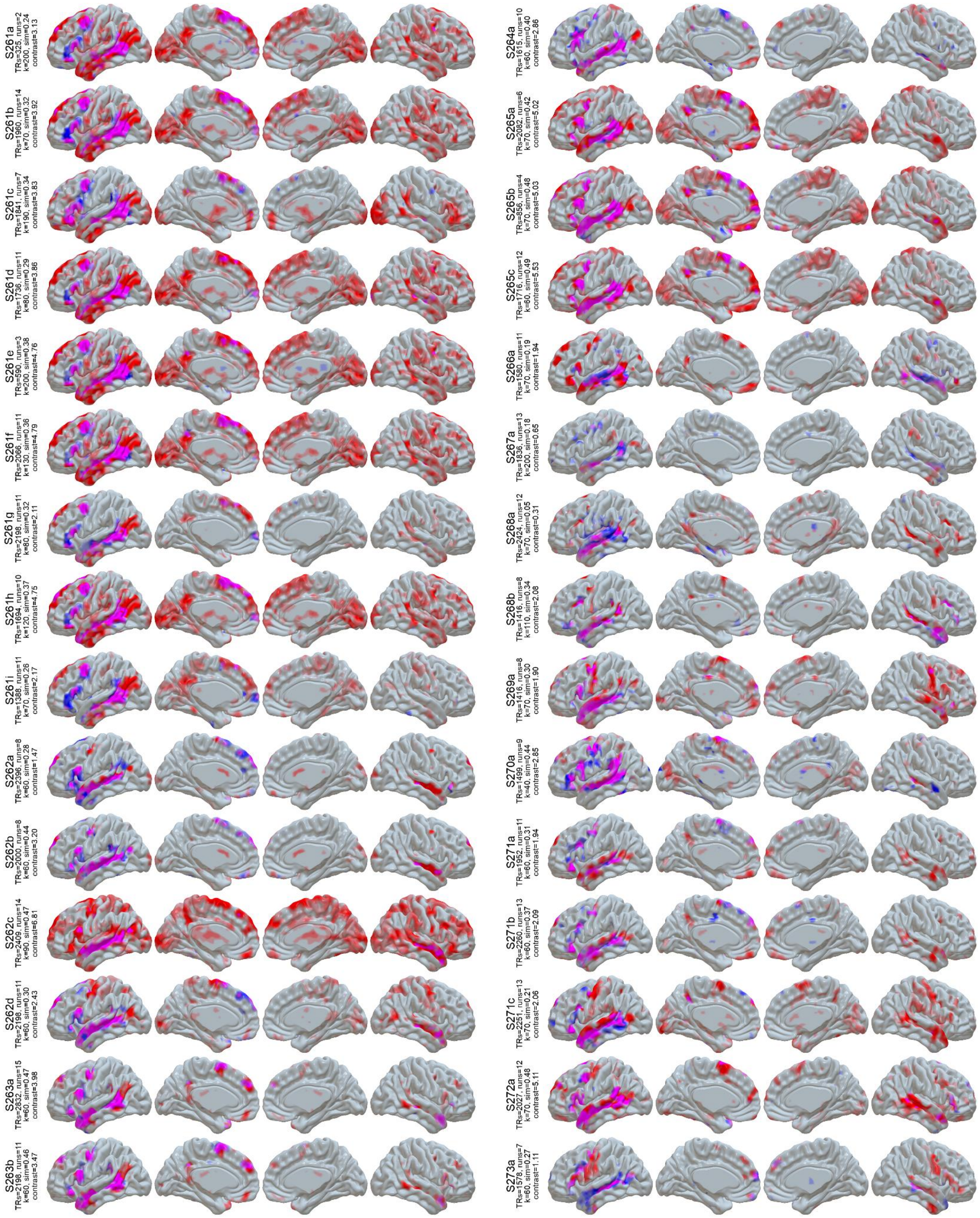


**Figure S1 (cont):** LangFC (blue) vs. task *t*-maps (sentences vs. nonword lists or *S-N*, red) in the 10 sessions with the highest S-N contrast stability between even and odd runs. Overlap is shown in magenta, and opacity reflects magnitude ($0.2 < p < 0.8$ for LangFC, $1 < t < 4$ for S-N).

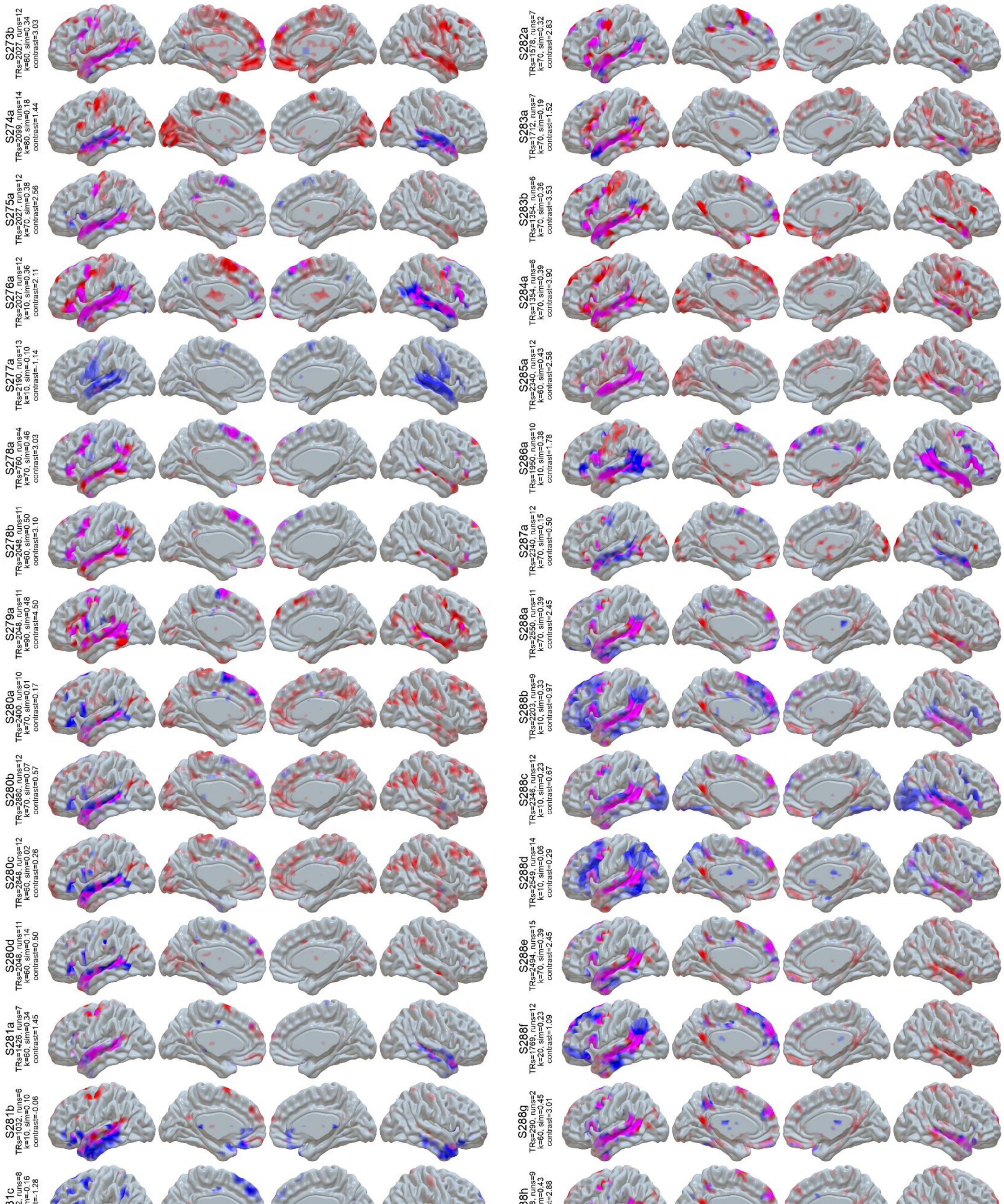


**Figure S1 (cont):** LangFC (blue) vs. task *t*-maps (sentences vs. nonword lists or *S-N*, red) in the 10 sessions with the highest S-N contrast stability between even and odd runs. Overlap is shown in magenta, and opacity reflects magnitude ($0.2 < p < 0.8$ for LangFC, $1 < t < 4$ for S-N).

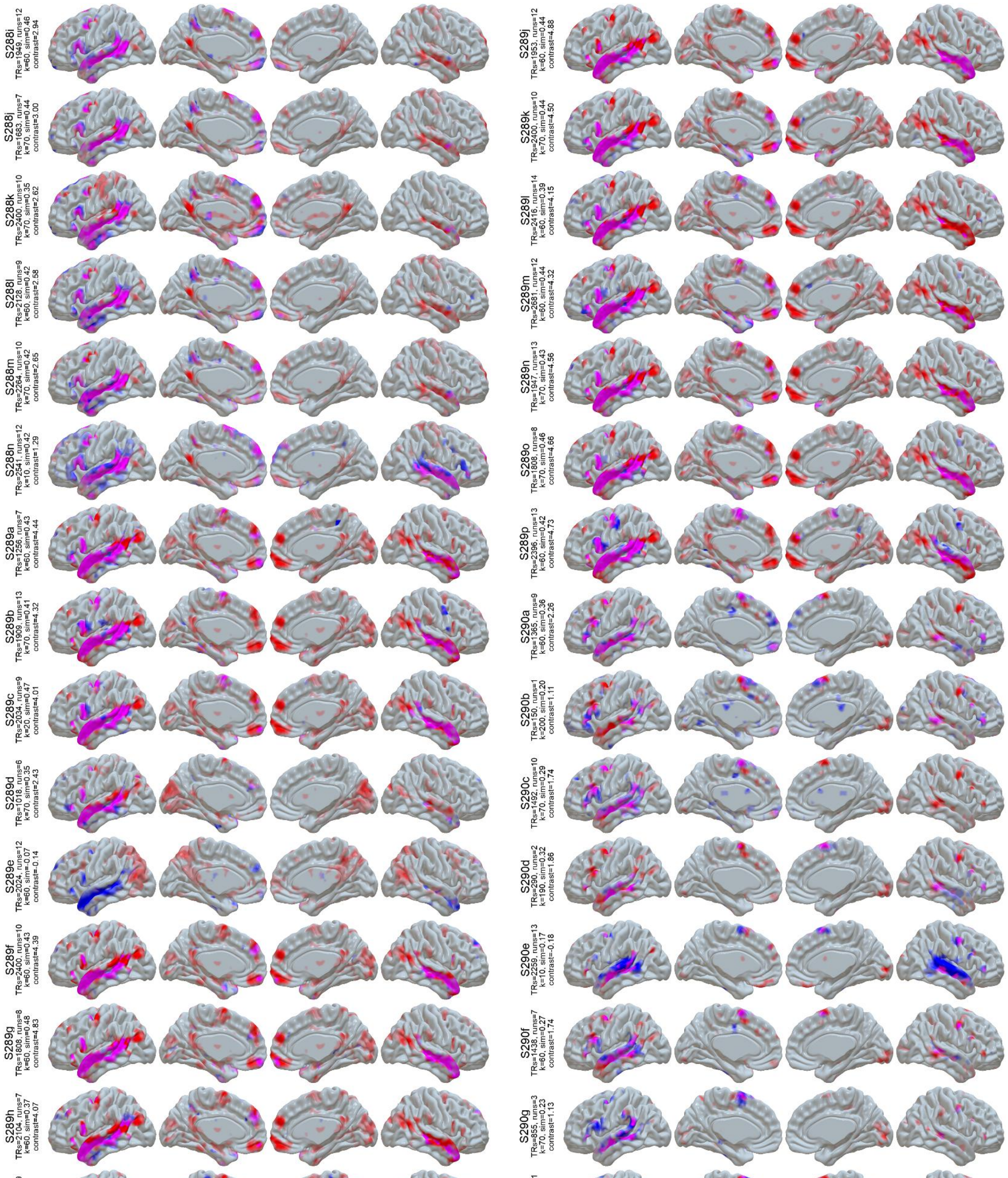


**Figure S1 (cont):** LangFC (blue) vs. task *t*-maps (sentences vs. nonword lists or *S-N*, red) in the 10 sessions with the highest S-N contrast stability between even and odd runs. Overlap is shown in magenta, and opacity reflects magnitude (0.2 < *p* < 0.8 for LangFC, 1 < *t* < 4 for S-N).

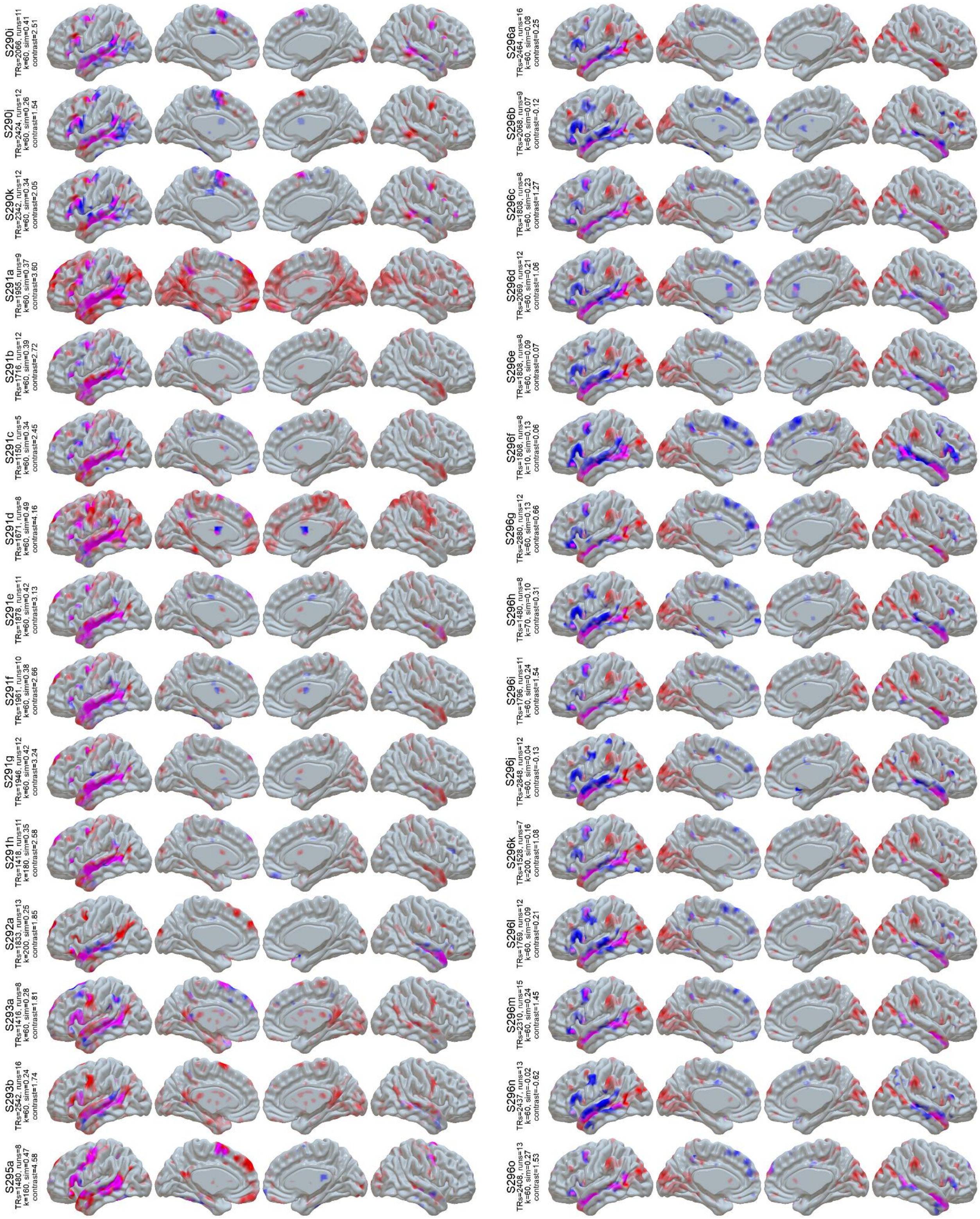


**Figure S1 (cont):** LangFC (blue) vs. task *t*-maps (sentences vs. nonword lists or *S-N*, red) in the 10 sessions with the highest S-N contrast stability between even and odd runs. Overlap is shown in magenta, and opacity reflects magnitude (0.2 < *p* < 0.8 for LangFC, 1 < *t* < 4 for S-N).

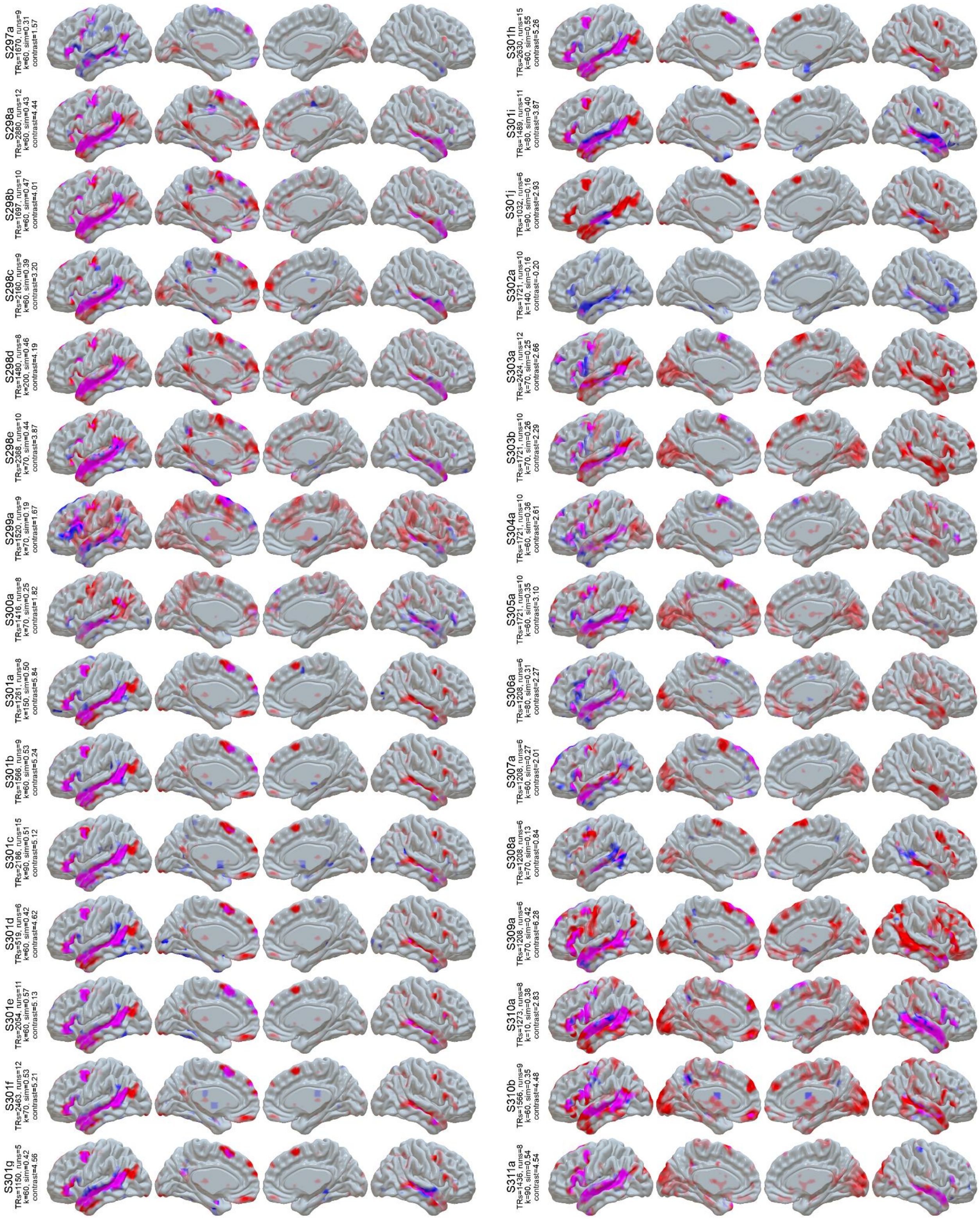


**Figure S1 (cont):** LangFC (blue) vs. task *t*-maps (sentences vs. nonword lists or *S-N*, red) in the 10 sessions with the highest S-N contrast stability between even and odd runs. Overlap is shown in magenta, and opacity reflects magnitude ($0.2 < p < 0.8$ for LangFC, $1 < t < 4$ for S-N).

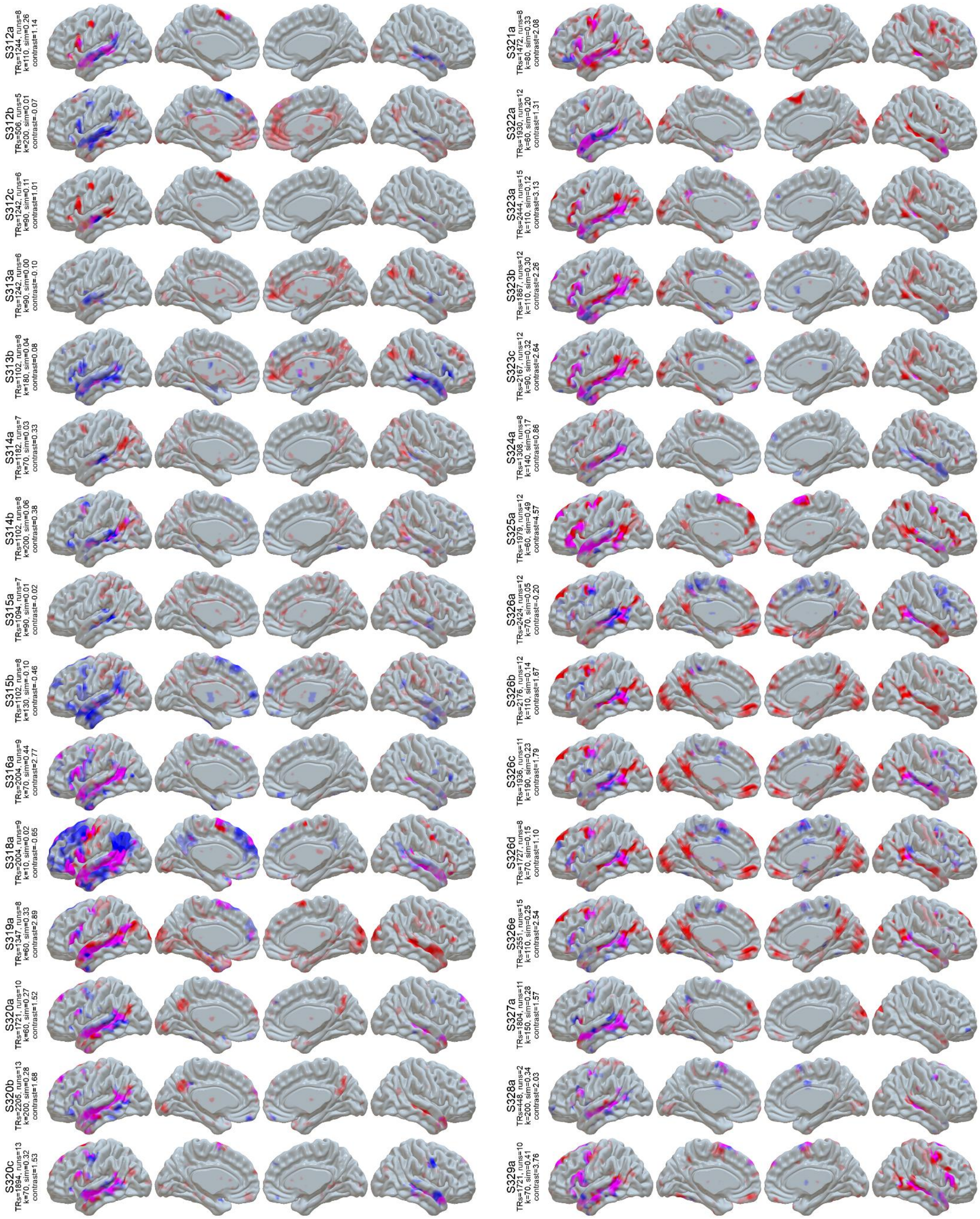


**Figure S1 (cont):** LangFC (blue) vs. task *t*-maps (sentences vs. nonword lists or *S-N*, red) in the 10 sessions with the highest S-N contrast stability between even and odd runs. Overlap is shown in magenta, and opacity reflects magnitude ($0.2 < p < 0.8$ for LangFC, $1 < t < 4$ for S-N).

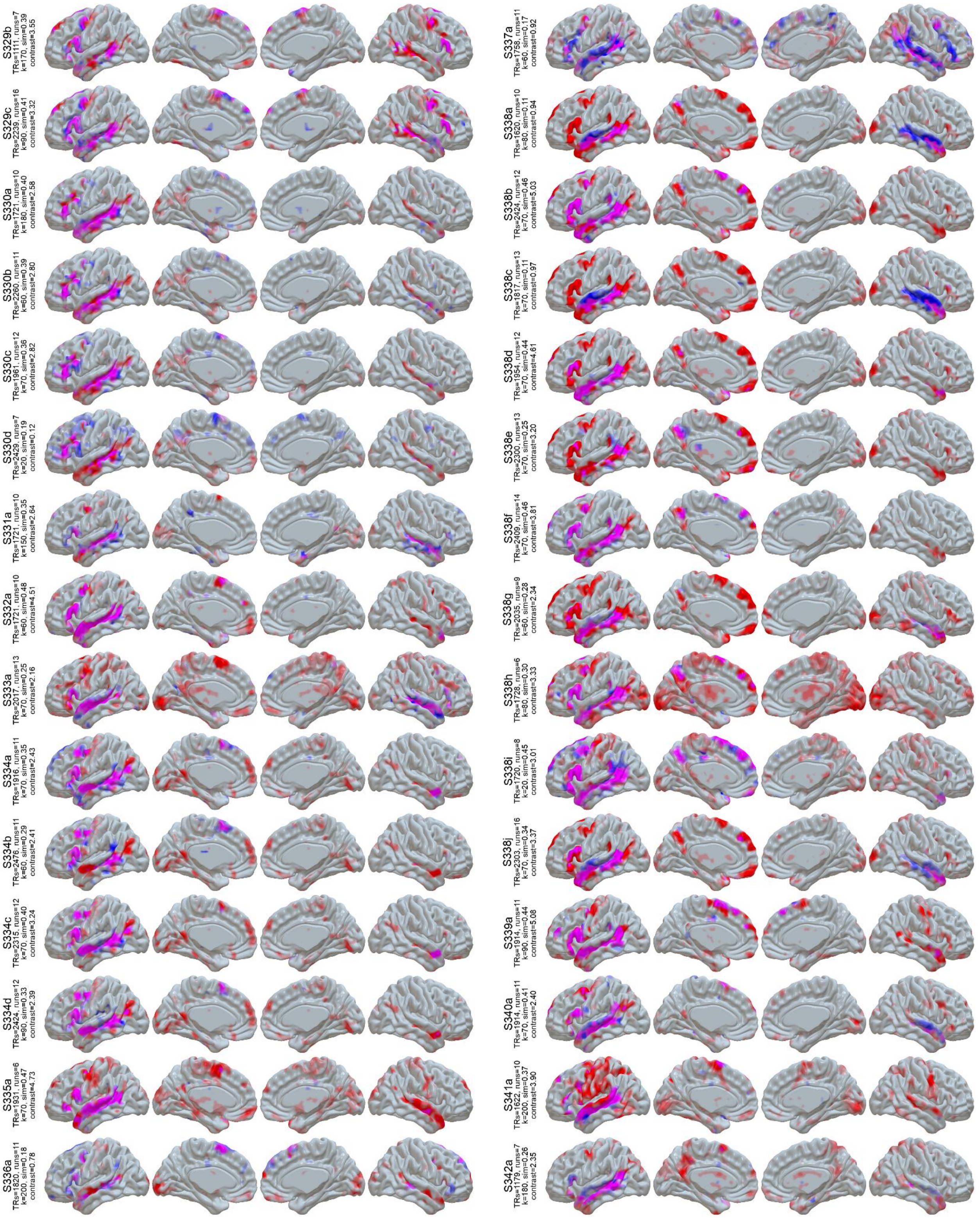


**Figure S1 (cont):** LangFC (blue) vs. task *t*-maps (sentences vs. nonword lists or *S-N*, red) in the 10 sessions with the highest S-N contrast stability between even and odd runs. Overlap is shown in magenta, and opacity reflects magnitude (0.2 < *p* < 0.8 for LangFC, 1 < *t* < 4 for S-N).

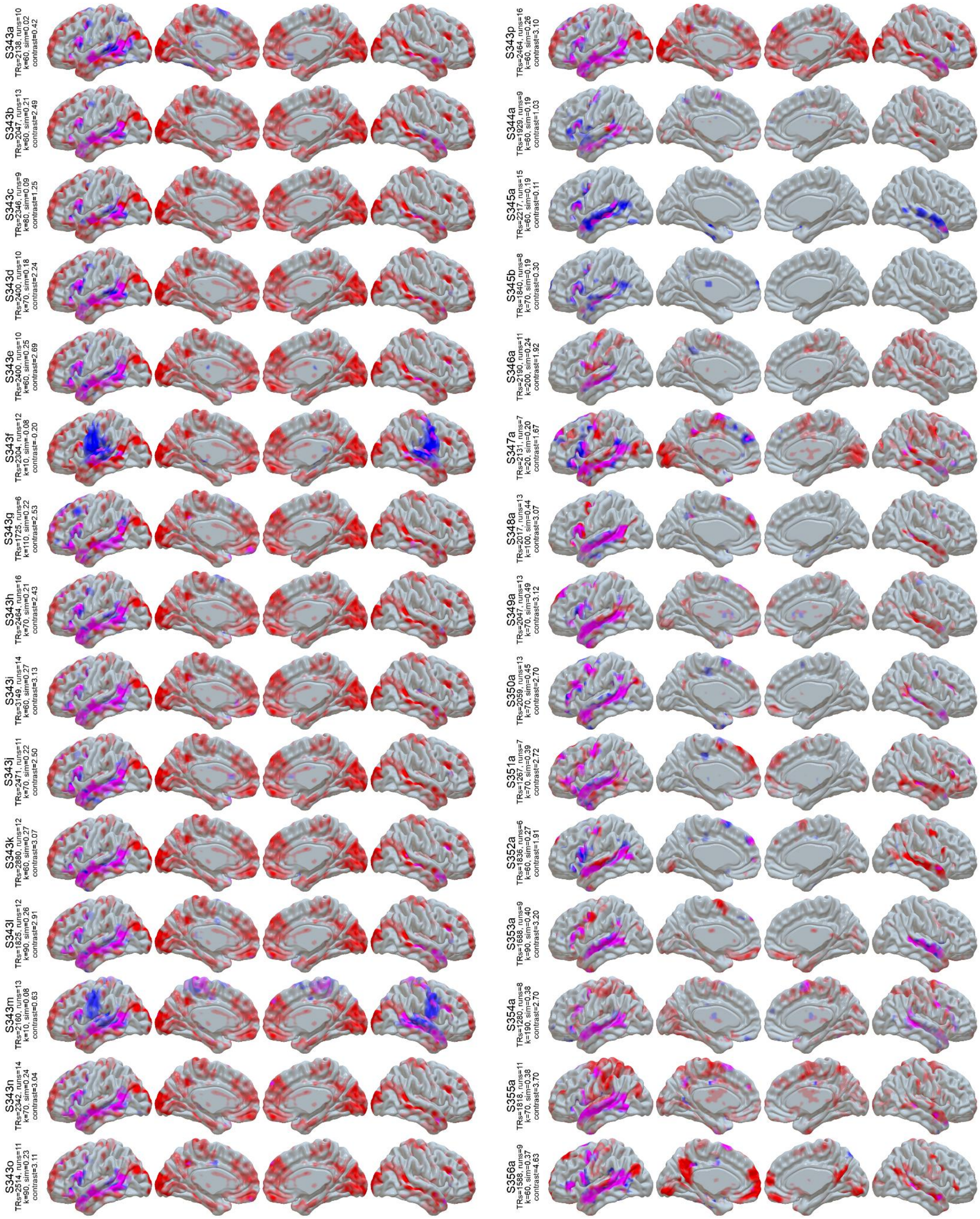


**Figure S1 (cont):** LangFC (blue) vs. task *t*-maps (sentences vs. nonword lists or *S-N*, red) in the 10 sessions with the highest S-N contrast stability between even and odd runs. Overlap is shown in magenta, and opacity reflects magnitude (0.2 < *p* < 0.8 for LangFC, 1 < *t* < 4 for S-N).

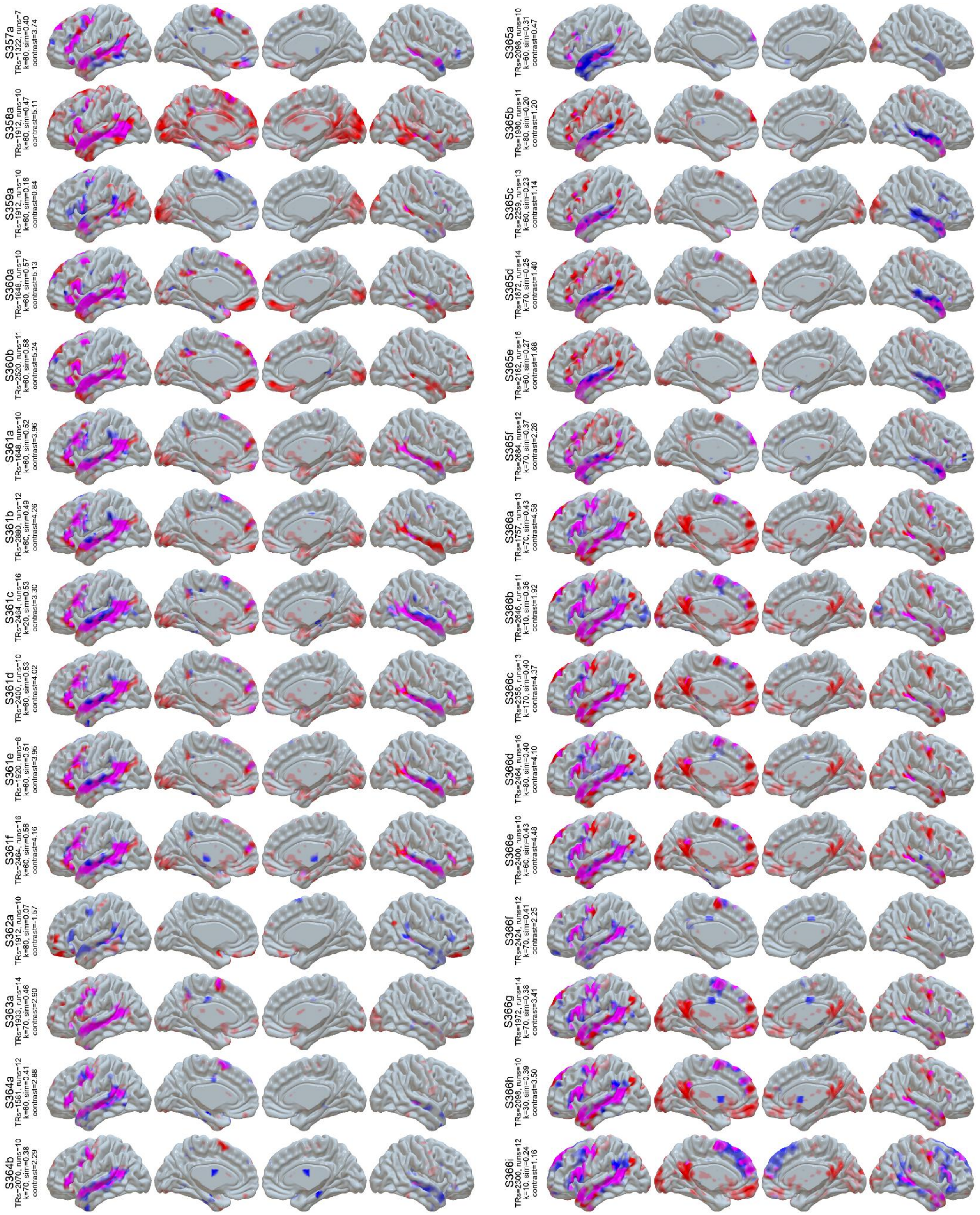


**Figure S1 (cont):** LangFC (blue) vs. task *t*-maps (sentences vs. nonword lists or *S-N*, red) in the 10 sessions with the highest S-N contrast stability between even and odd runs. Overlap is shown in magenta, and opacity reflects magnitude (0.2 < *p* < 0.8 for LangFC, 1 < *t* < 4 for S-N).

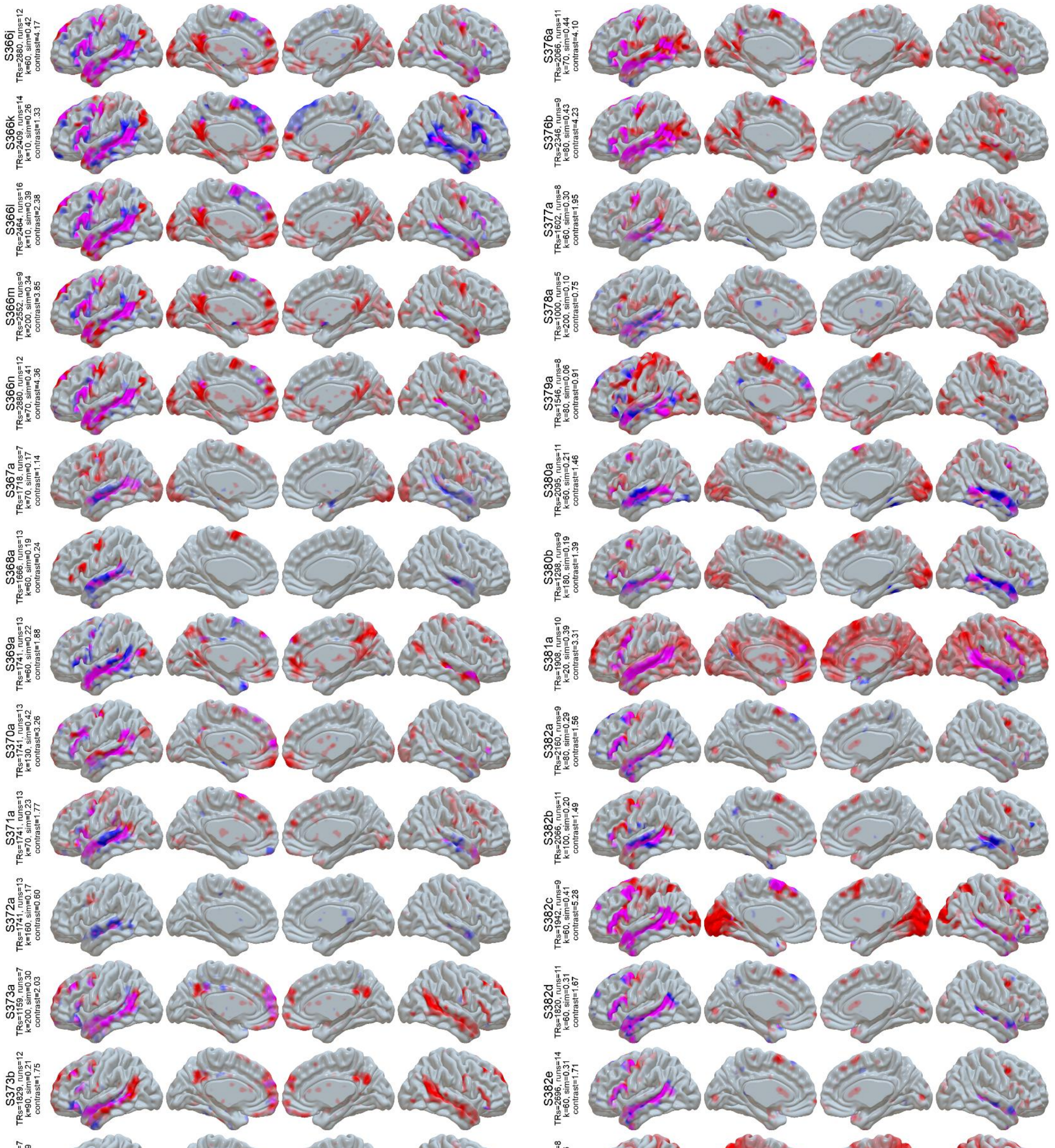


**Figure S1 (cont):** LangFC (blue) vs. task *t*-maps (sentences vs. nonword lists or *S-N*, red) in the 10 sessions with the highest S-N contrast stability between even and odd runs. Overlap is shown in magenta, and opacity reflects magnitude ($0.2 < p < 0.8$ for LangFC, $1 < t < 4$ for S-N).

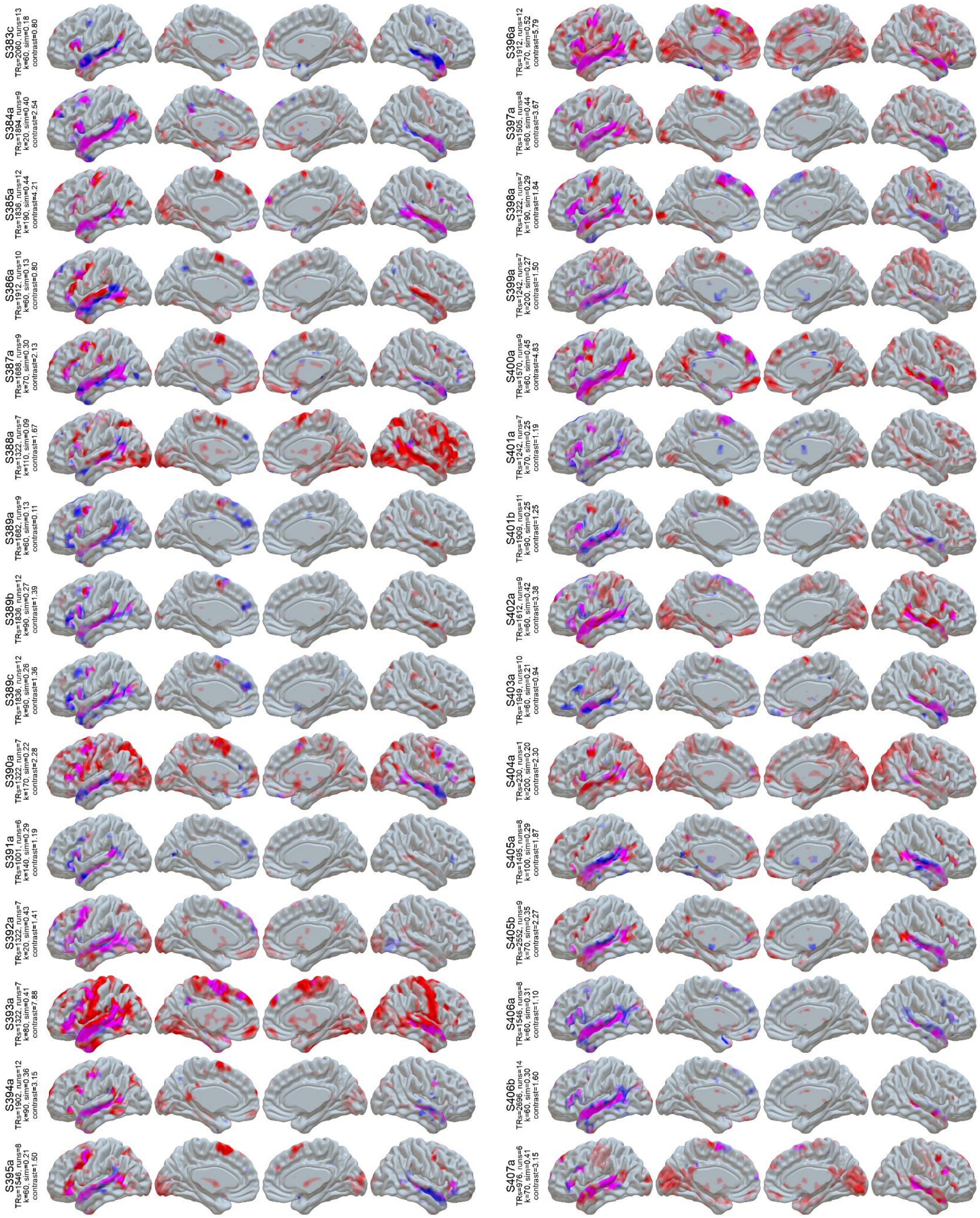

**Figure S1 (cont):** LangFC (blue) vs. task *t*-maps (sentences vs. nonword lists or *S-N*, red) in the 10 sessions with the highest S-N contrast stability between even and odd runs. Overlap is shown in magenta, and opacity reflects magnitude ($0.2 < p < 0.8$ for LangFC, $1 < t < 4$ for S-N).

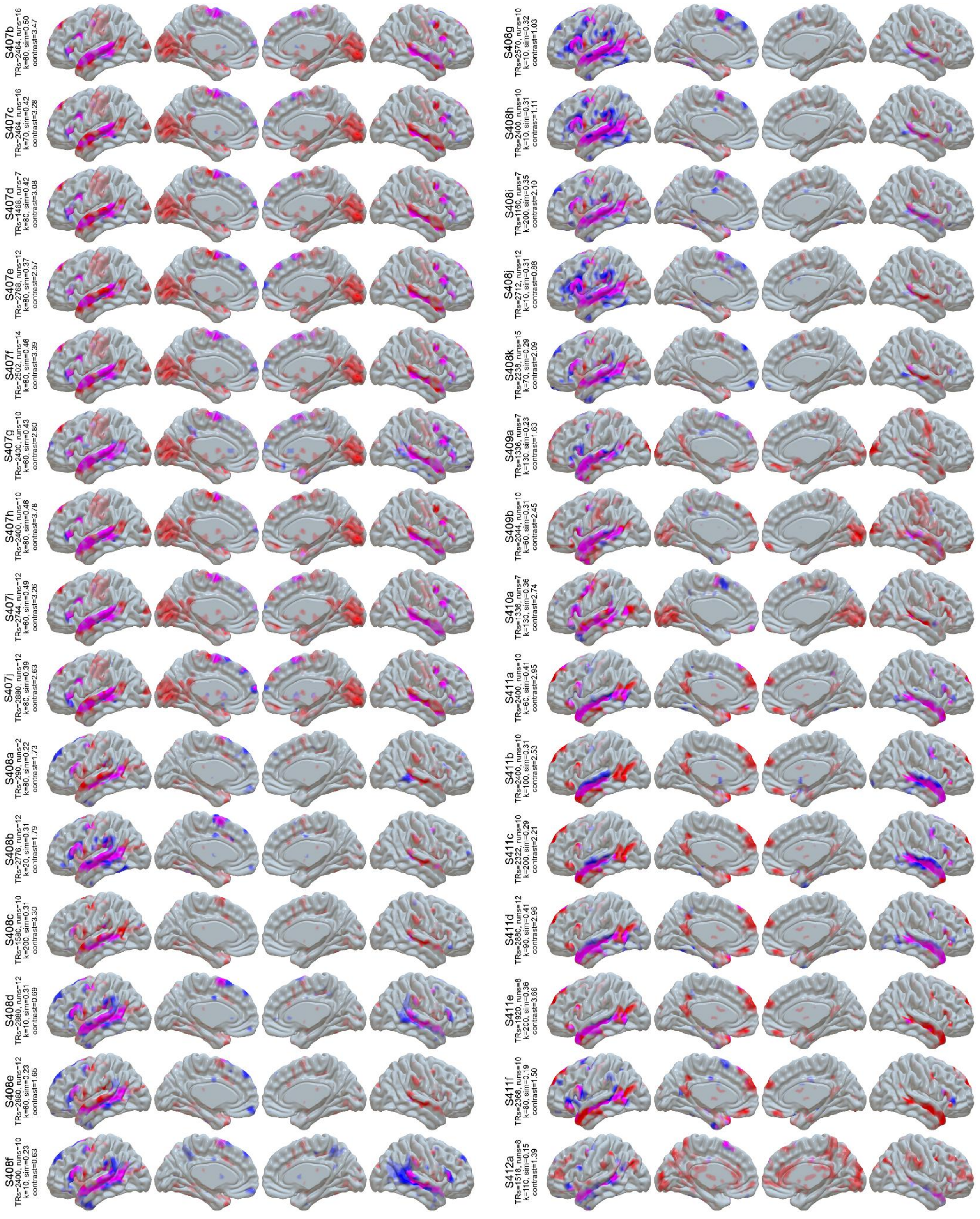


**Figure S1 (cont):** LangFC (blue) vs. task *t*-maps (sentences vs. nonword lists or *S-N*, red) in the 10 sessions with the highest S-N contrast stability between even and odd runs. Overlap is shown in magenta, and opacity reflects magnitude (0.2 < *p* < 0.8 for LangFC, 1 < *t* < 4 for S-N).

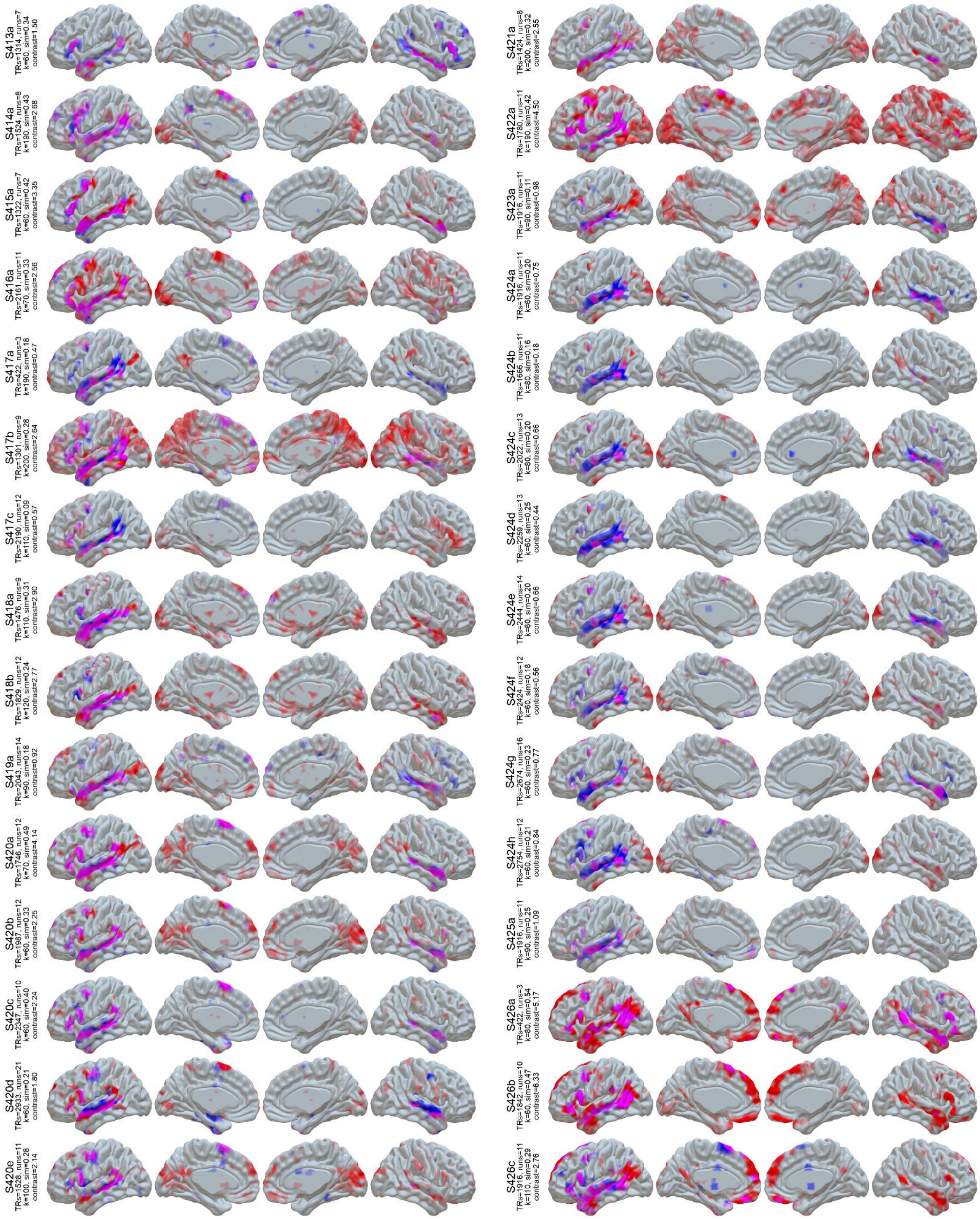


**Figure S1 (cont):** LangFC (blue) vs. task *t*-maps (sentences vs. nonword lists or *S-N*, red) in the 10 sessions with the highest S-N contrast stability between even and odd runs. Overlap is shown in magenta, and opacity reflects magnitude (0.2 < $p$ < 0.8 for LangFC, 1 < $t$ < 4 for S-N).

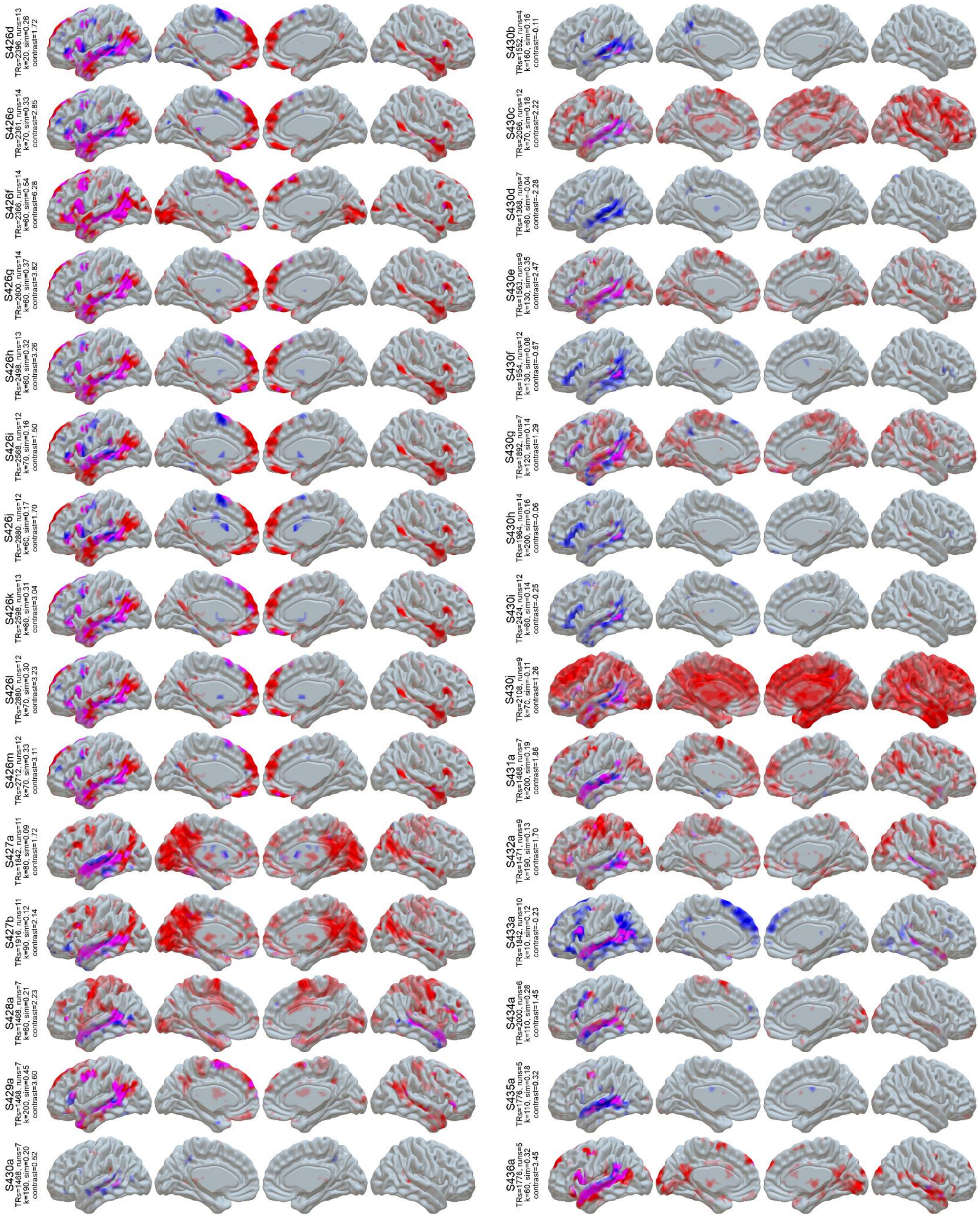


**Figure S1 (cont):** LangFC (blue) vs. task *t*-maps (sentences vs. nonword lists or *S-N*, red) in the 10 sessions with the highest S-N contrast stability between even and odd runs. Overlap is shown in magenta, and opacity reflects magnitude (0.2 < $p$ < 0.8 for LangFC, 1 < $t$ < 4 for S-N).

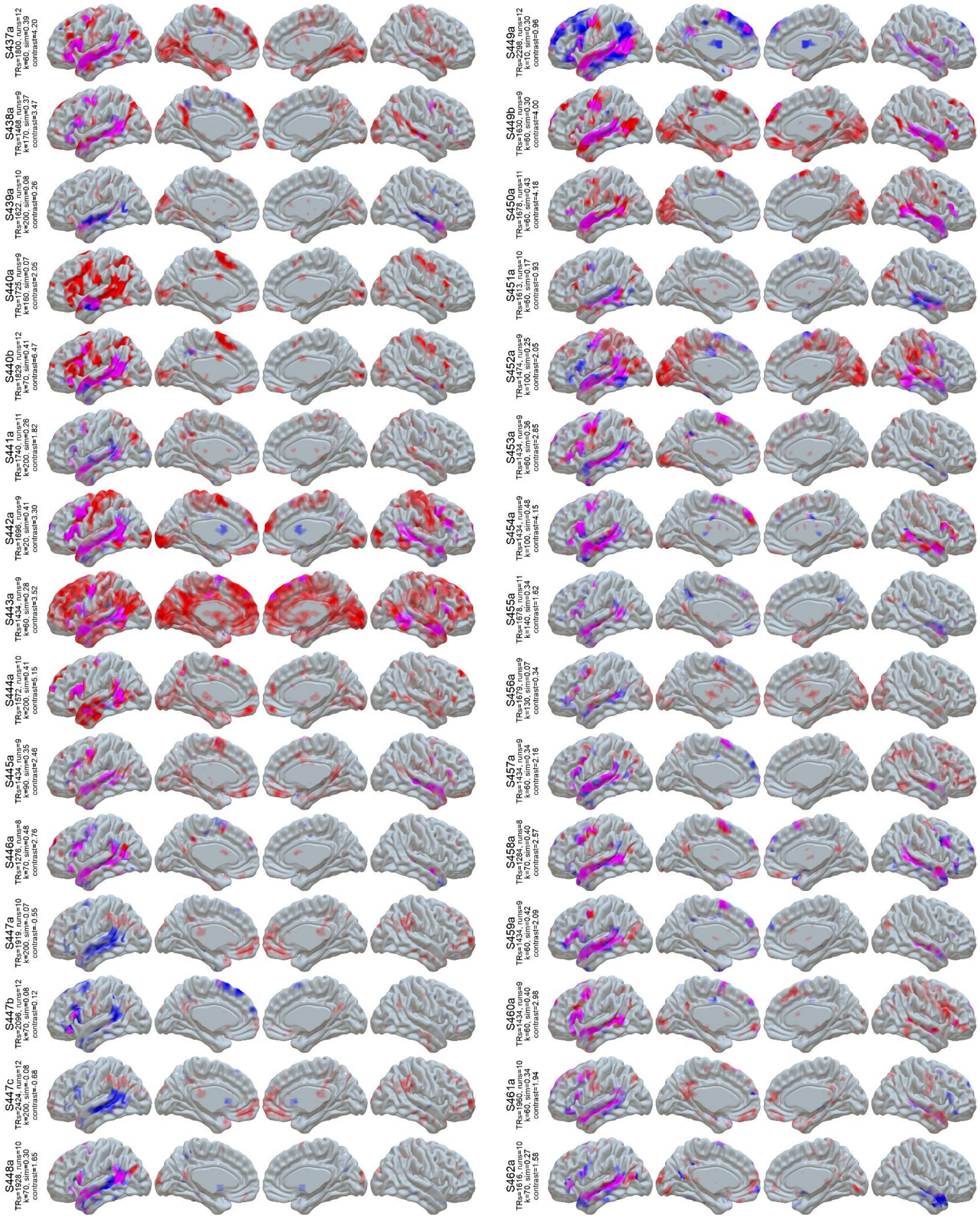

**Figure S1 (cont):** LangFC (blue) vs. task *t*-maps (sentences vs. nonword lists or *S-N*, red) in the 10 sessions with the highest S-N contrast stability between even and odd runs. Overlap is shown in magenta, and opacity reflects magnitude ($0.2 < p < 0.8$ for LangFC, $1 < t < 4$ for S-N).

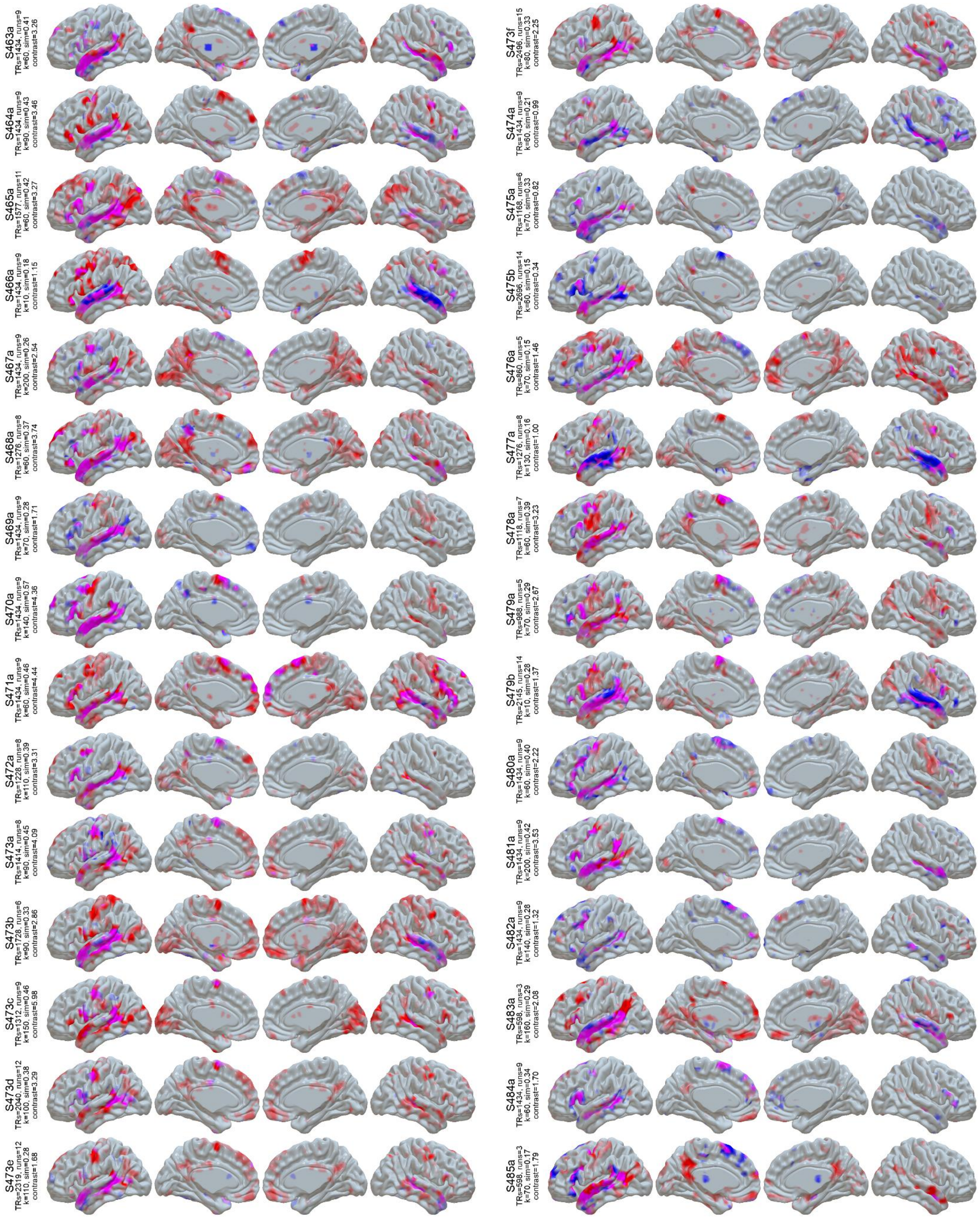


**Figure S1 (cont):** LangFC (blue) vs. task *t*-maps (sentences vs. nonword lists or *S-N*, red) in the 10 sessions with the highest S-N contrast stability between even and odd runs. Overlap is shown in magenta, and opacity reflects magnitude (0.2 < *p* < 0.8 for LangFC, 1 < *t* < 4 for S-N).

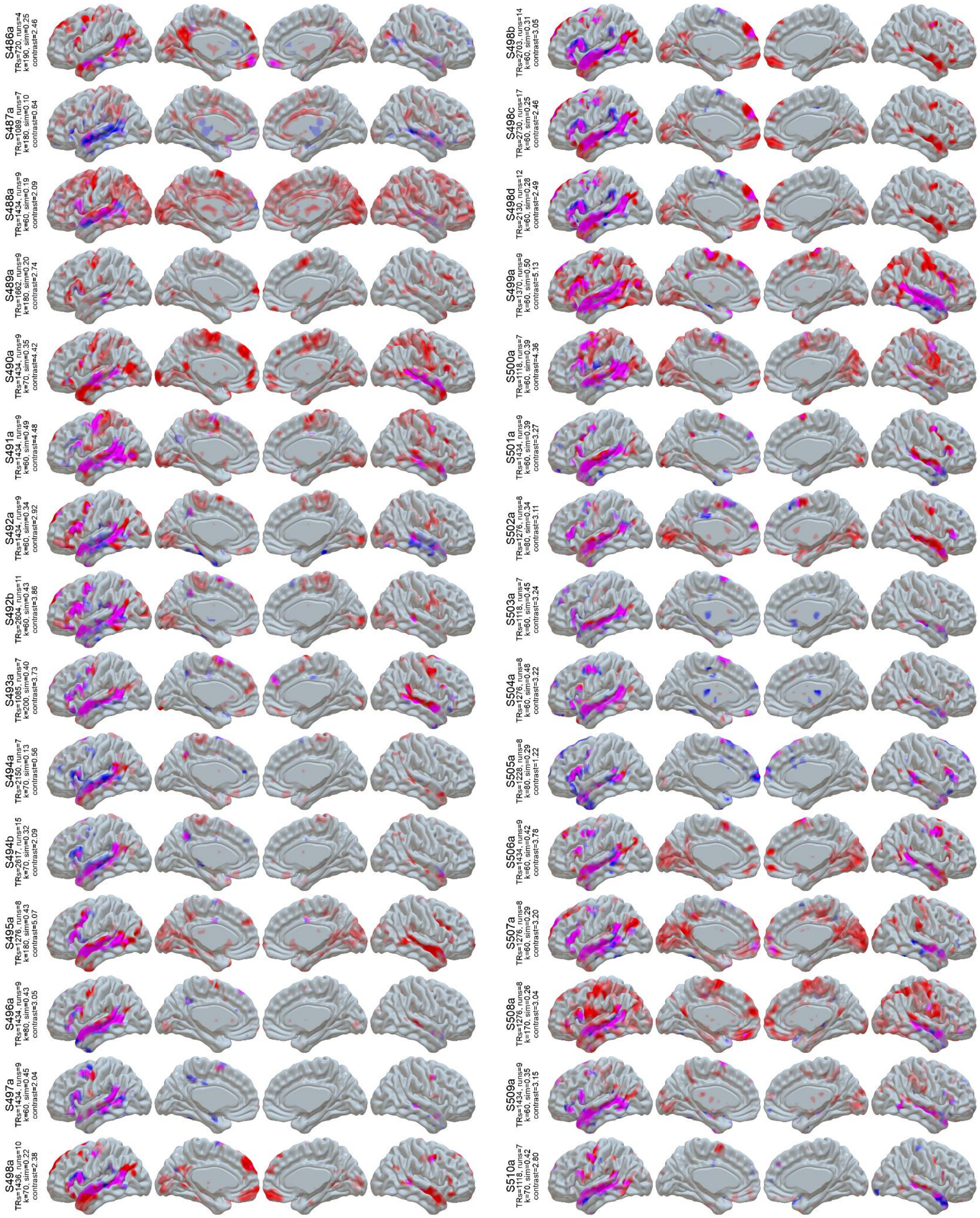

**Figure S1 (cont):** LangFC (blue) vs. task *t*-maps (sentences vs. nonword lists or *S-N*, red) in the 10 sessions with the highest S-N contrast stability between even and odd runs. Overlap is shown in magenta, and opacity reflects magnitude (0.2 < $p$ < 0.8 for LangFC, 1 < $t$ < 4 for S-N).

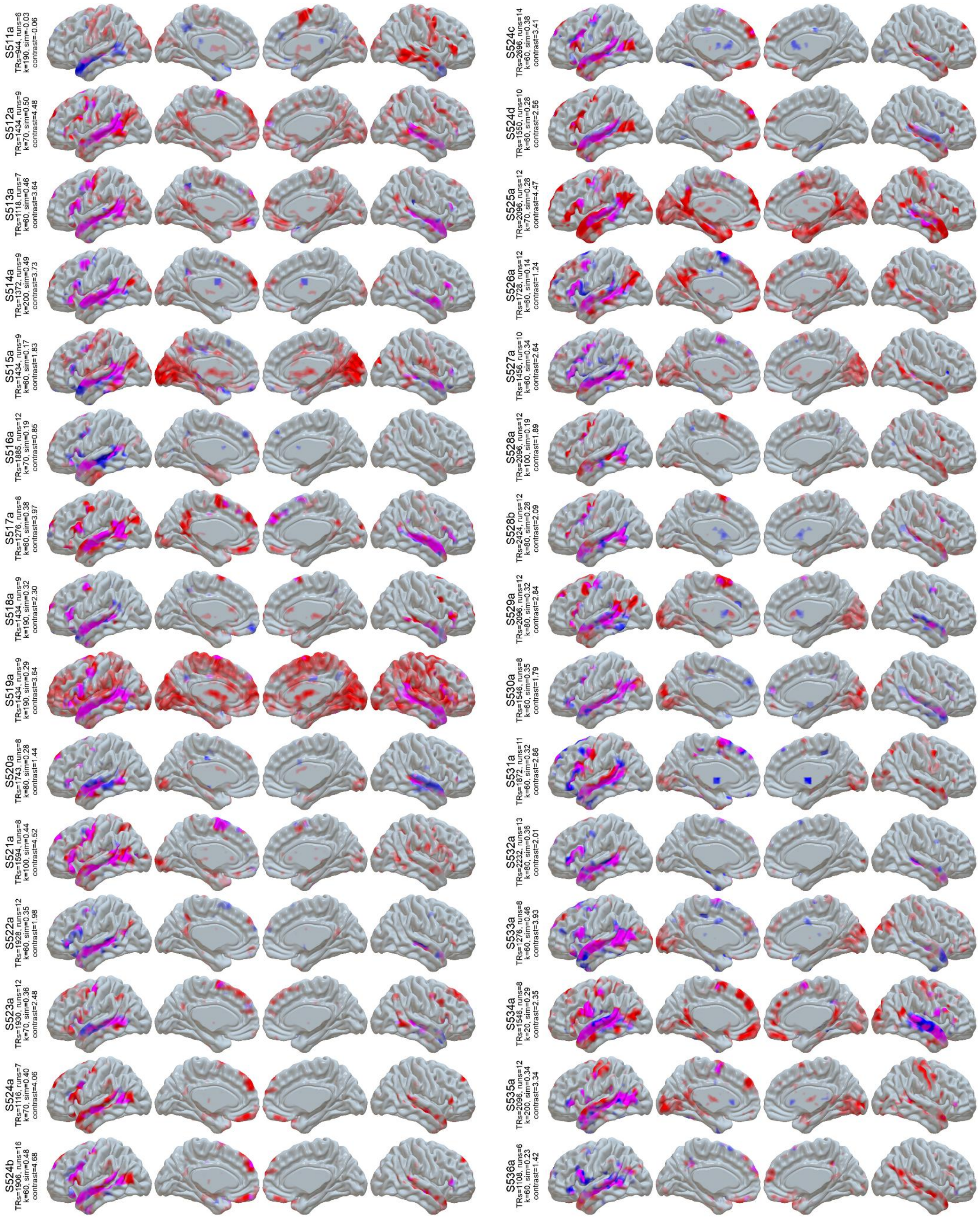


**Figure S1 (cont):** LangFC (blue) vs. task *t*-maps (sentences vs. nonword lists or *S-N*, red) in the 10 sessions with the highest S-N contrast stability between even and odd runs. Overlap is shown in magenta, and opacity reflects magnitude (0.2 < *p* < 0.8 for LangFC, 1 < *t* < 4 for S-N).

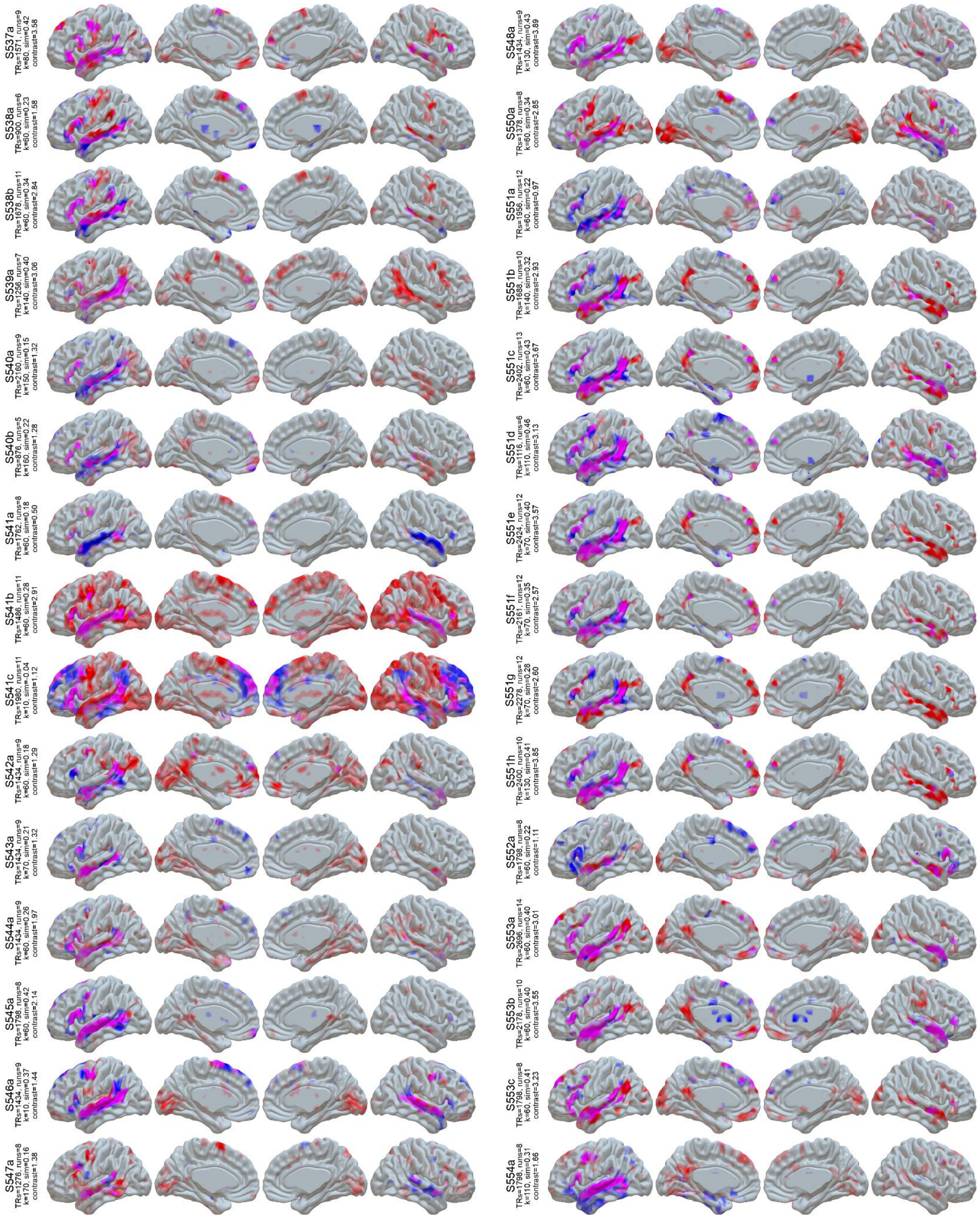


**Figure S1 (cont):** LangFC (blue) vs. task *t*-maps (sentences vs. nonword lists or *S-N*, red) in the 10 sessions with the highest S-N contrast stability between even and odd runs. Overlap is shown in magenta, and opacity reflects magnitude ($0.2 < p < 0.8$ for LangFC, $1 < t < 4$ for S-N).

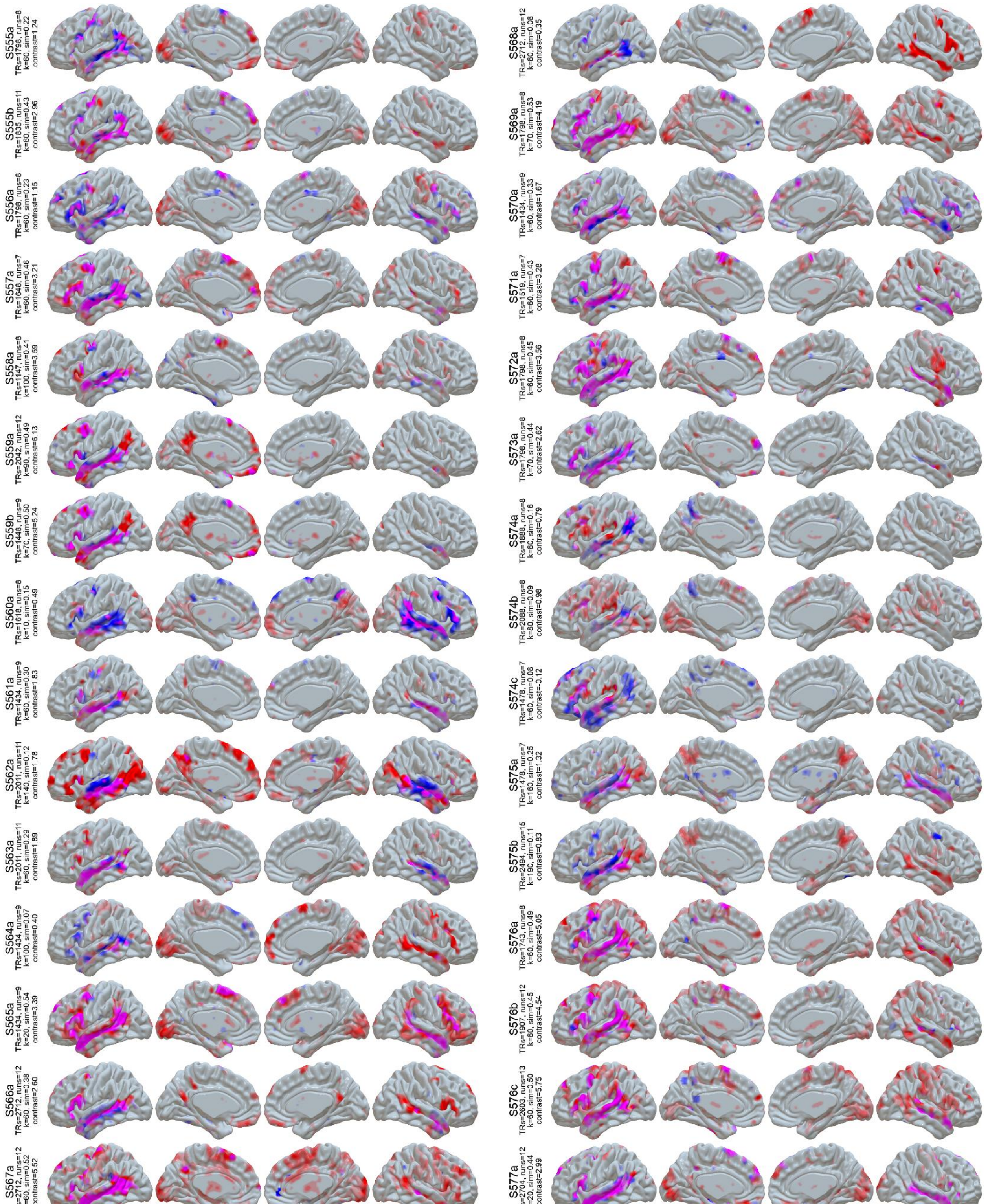


**Figure S1 (cont):** LangFC (blue) vs. task *t*-maps (sentences vs. nonword lists or *S-N*, red) in the 10 sessions with the highest S-N contrast stability between even and odd runs. Overlap is shown in magenta, and opacity reflects magnitude (0.2 < *p* < 0.8 for LangFC, 1 < *t* < 4 for S-N).

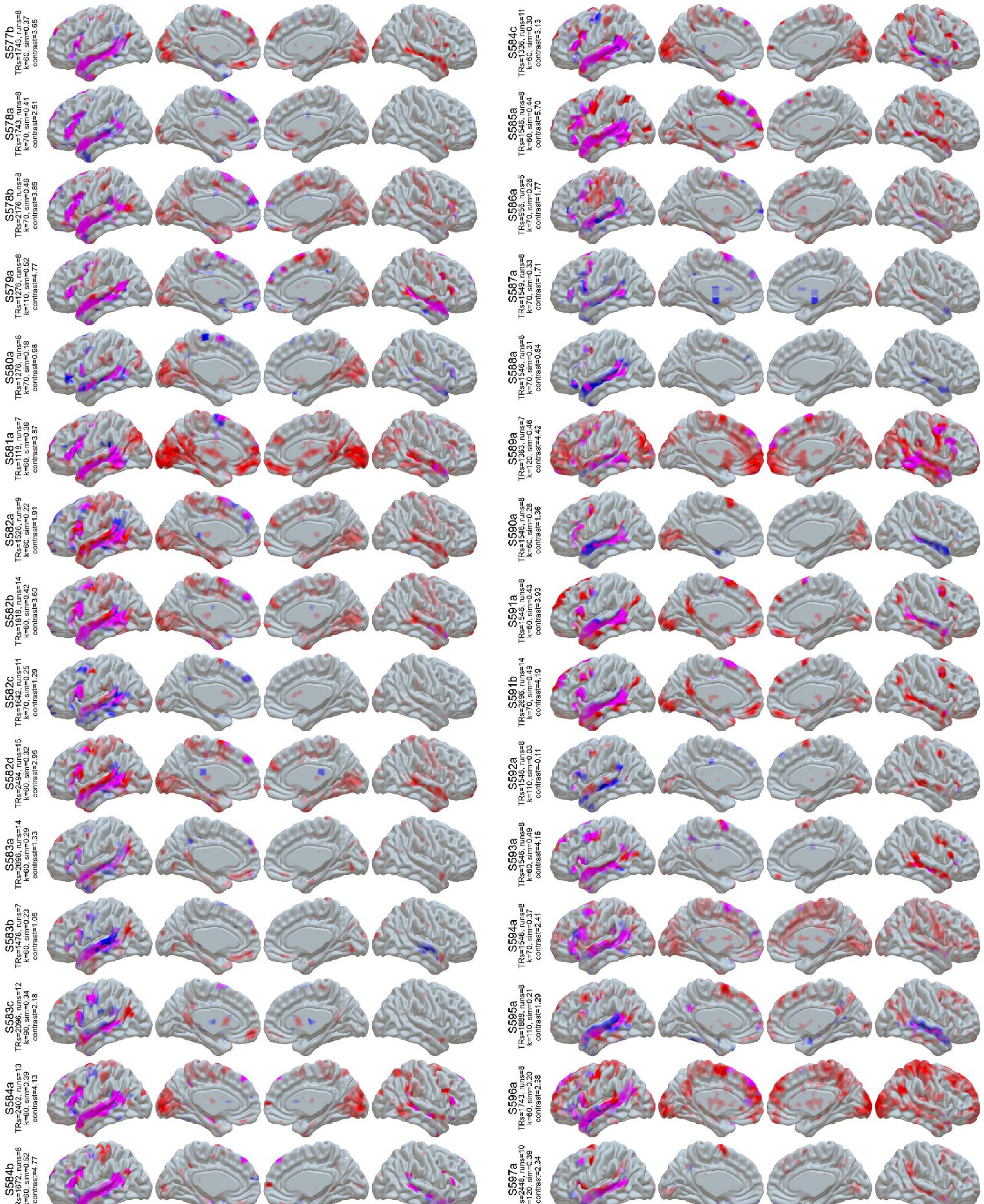


**Figure S1 (cont):** LangFC (blue) vs. task *t*-maps (sentences vs. nonword lists or *S-N*, red) in the 10 sessions with the highest S-N contrast stability between even and odd runs. Overlap is shown in magenta, and opacity reflects magnitude (0.2 < *p* < 0.8 for LangFC, 1 < *t* < 4 for S-N).

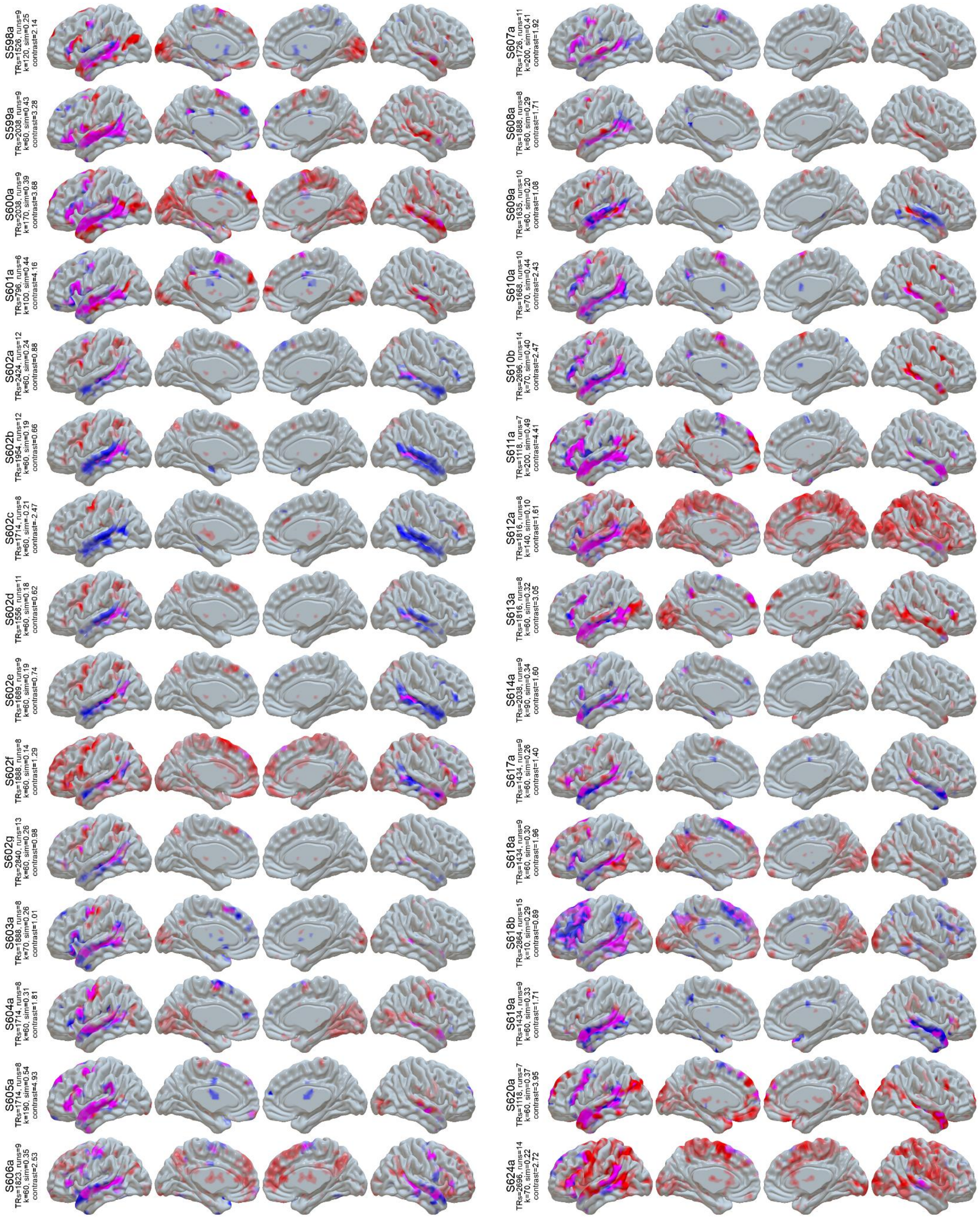


**Figure S1 (cont):** LangFC (blue) vs. task *t*-maps (sentences vs. nonword lists or *S-N*, red) in the 10 sessions with the highest S-N contrast stability between even and odd runs. Overlap is shown in magenta, and opacity reflects magnitude (0.2 < *p* < 0.8 for LangFC, 1 < *t* < 4 for S-N).

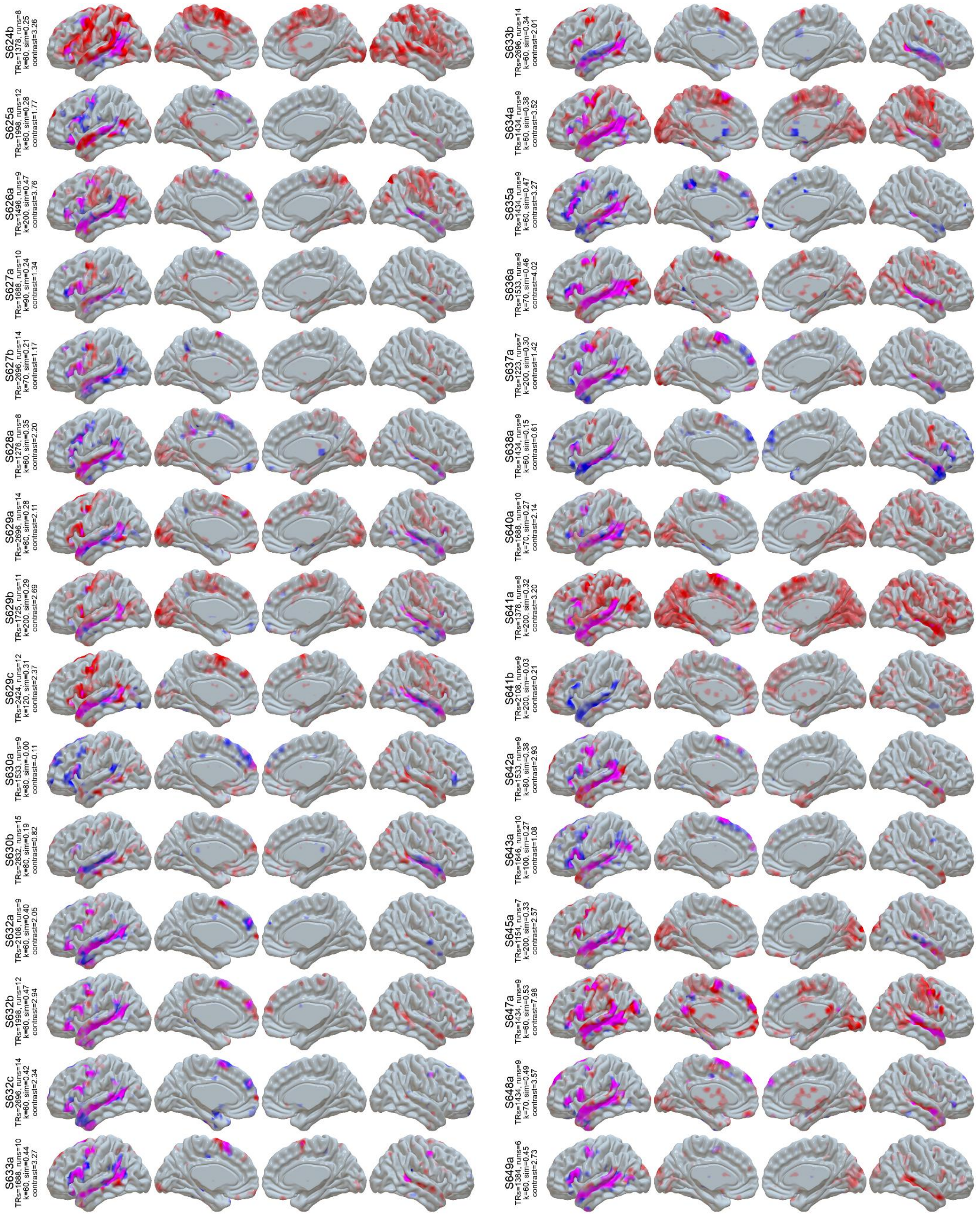


**Figure S1 (cont):** LangFC (blue) vs. task *t*-maps (sentences vs. nonword lists or *S-N*, red) in the 10 sessions with the highest S-N contrast stability between even and odd runs. Overlap is shown in magenta, and opacity reflects magnitude (0.2 < *p* < 0.8 for LangFC, 1 < *t* < 4 for S-N).

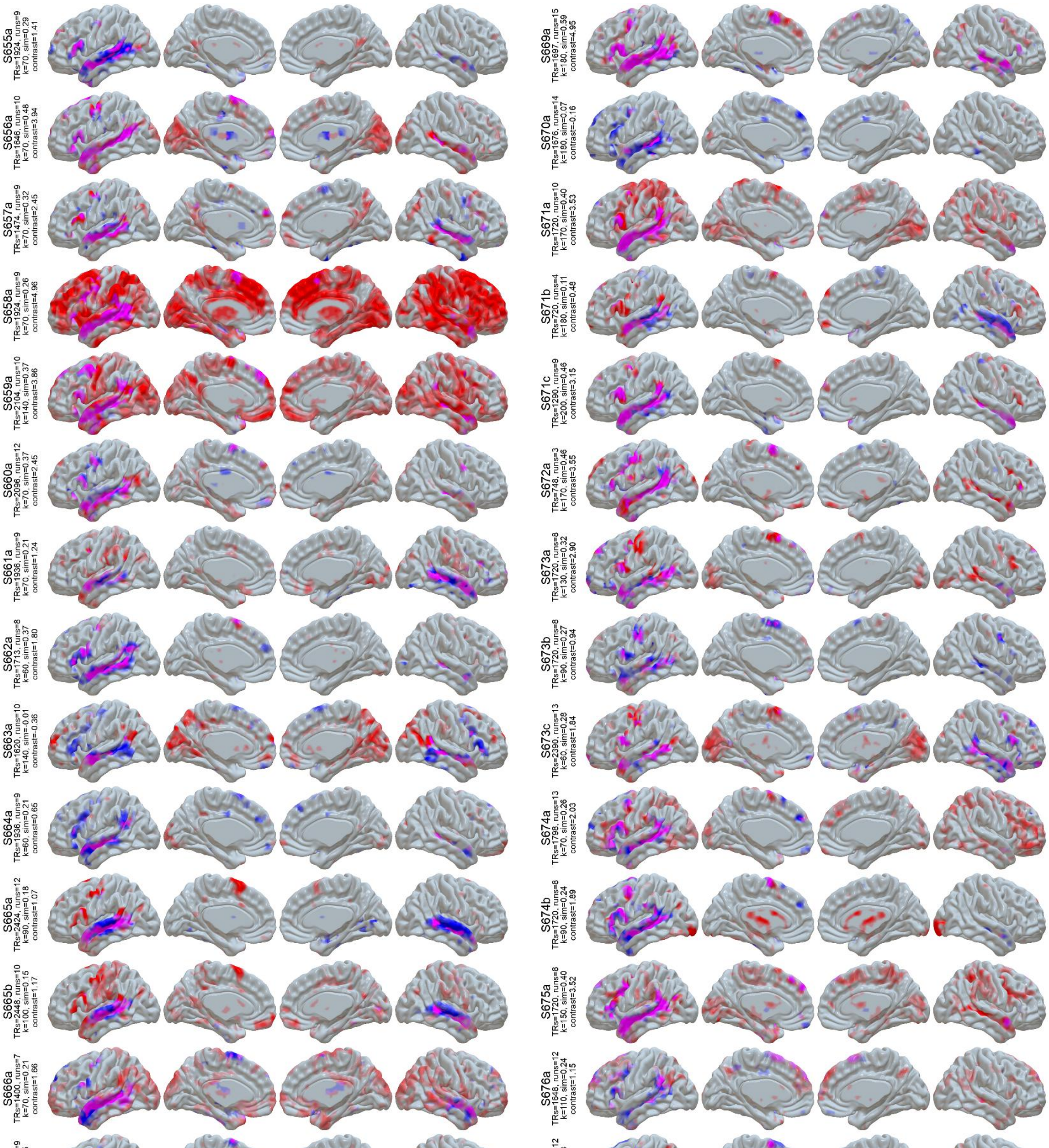


**Figure S1 (cont):** LangFC (blue) vs. task *t*-maps (sentences vs. nonword lists or *S-N*, red) in the 10 sessions with the highest S-N contrast stability between even and odd runs. Overlap is shown in magenta, and opacity reflects magnitude (0.2 < *p* < 0.8 for LangFC, 1 < *t* < 4 for S-N).

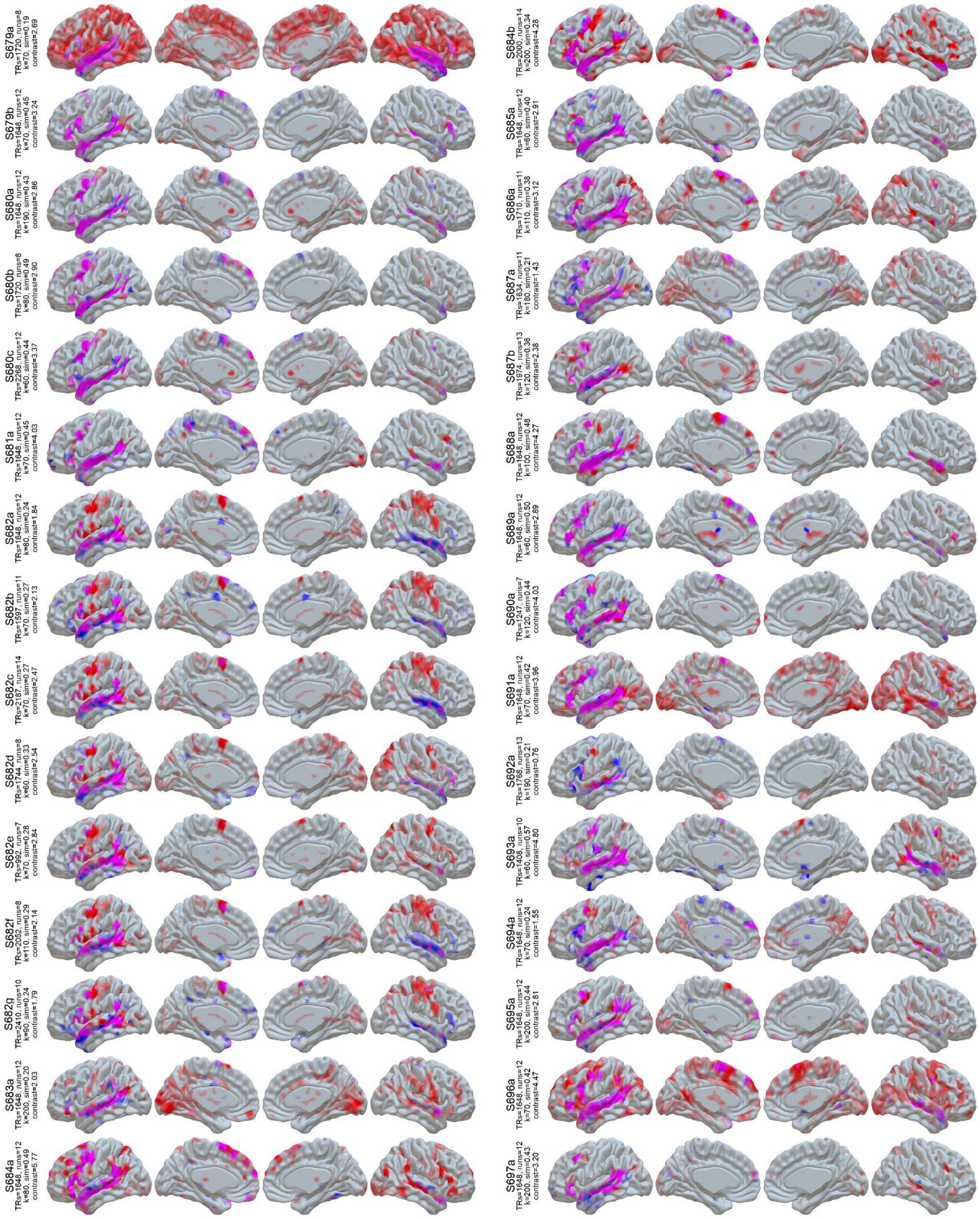

**Figure S1 (cont):** LangFC (blue) vs. task *t*-maps (sentences vs. nonword lists or *S-N*, red) in the 10 sessions with the highest S-N contrast stability between even and odd runs. Overlap is shown in magenta, and opacity reflects magnitude (0.2 < *p* < 0.8 for LangFC, 1 < *t* < 4 for S-N).

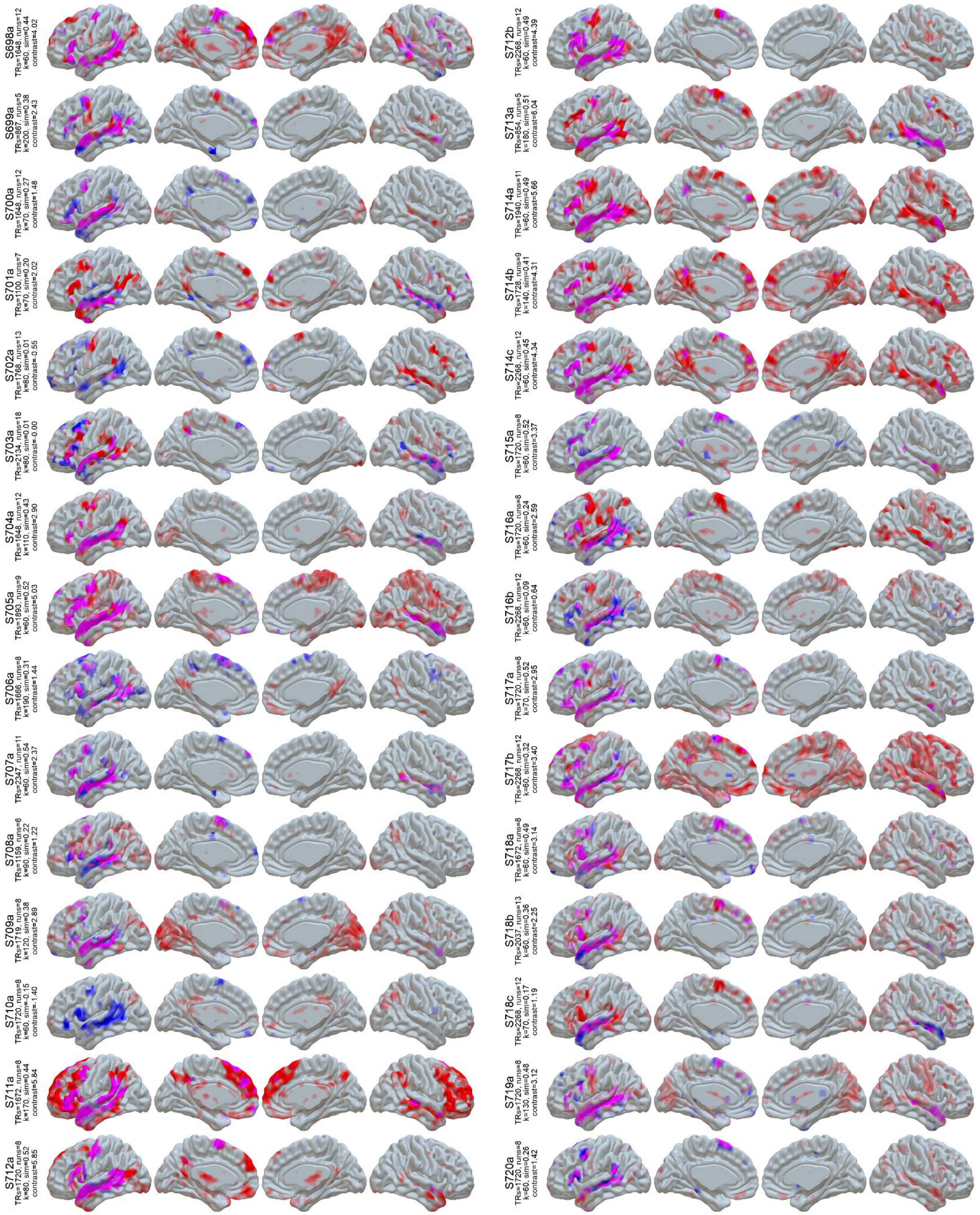


**Figure S1 (cont):** LangFC (blue) vs. task *t*-maps (sentences vs. nonword lists or *S-N*, red) in the 10 sessions with the highest S-N contrast stability between even and odd runs. Overlap is shown in magenta, and opacity reflects magnitude (0.2 < *p* < 0.8 for LangFC, 1 < *t* < 4 for S-N).

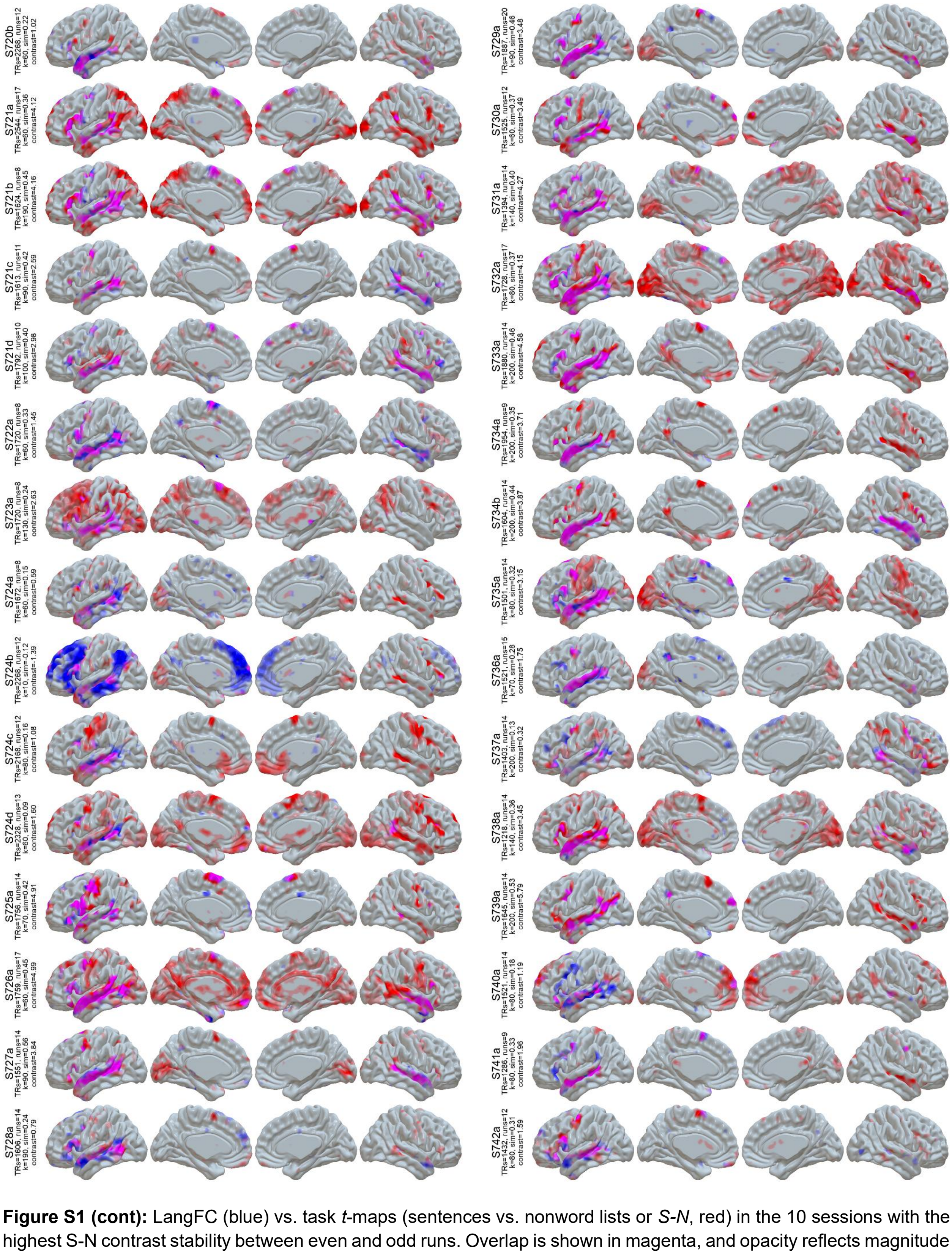


**Figure S1 (cont):** LangFC (blue) vs. task *t*-maps (sentences vs. nonword lists or *S-N*, red) in the 10 sessions with the highest S-N contrast stability between even and odd runs. Overlap is shown in magenta, and opacity reflects magnitude (0.2 < *p* < 0.8 for LangFC, 1 < *t* < 4 for S-N).

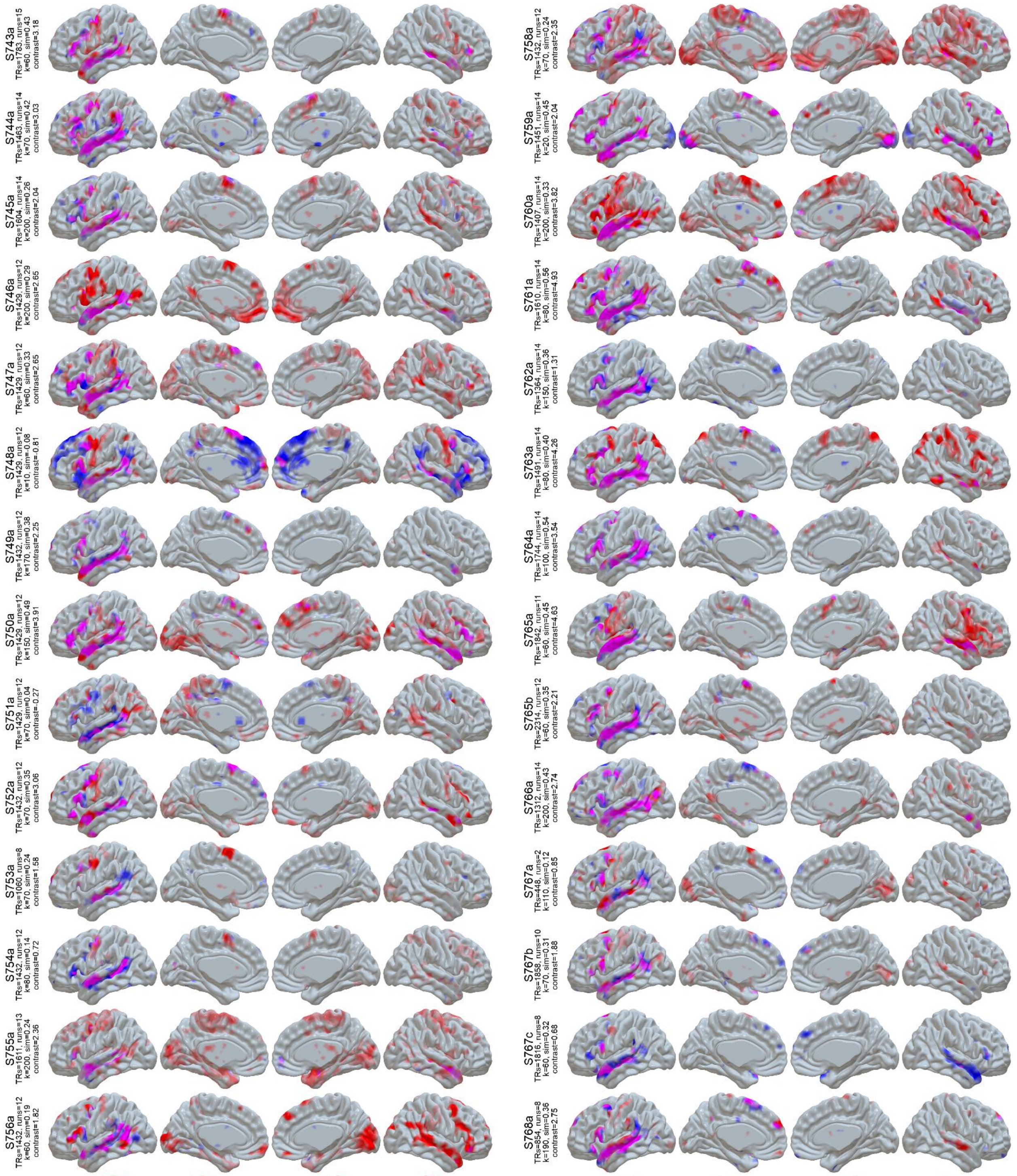


**Figure S1 (cont):** LangFC (blue) vs. task *t*-maps (sentences vs. nonword lists or *S-N*, red) in the 10 sessions with the highest S-N contrast stability between even and odd runs. Overlap is shown in magenta, and opacity reflects magnitude (0.2 < *p* < 0.8 for LangFC, 1 < *t* < 4 for S-N).

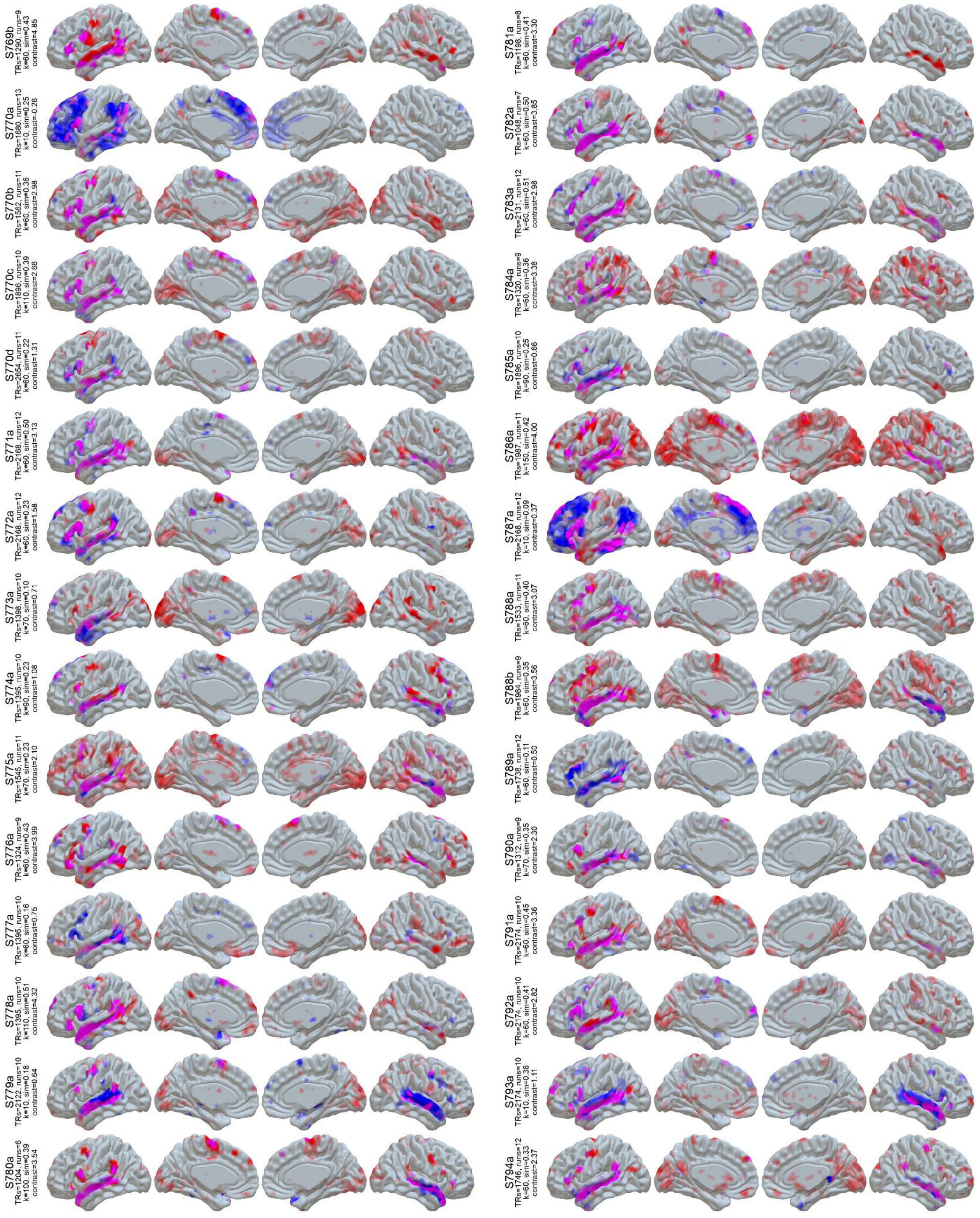

**Figure S1 (cont):** LangFC (blue) vs. task *t*-maps (sentences vs. nonword lists or *S-N*, red) in the 10 sessions with the highest S-N contrast stability between even and odd runs. Overlap is shown in magenta, and opacity reflects magnitude (0.2 < *p* < 0.8 for LangFC, 1 < *t* < 4 for S-N).

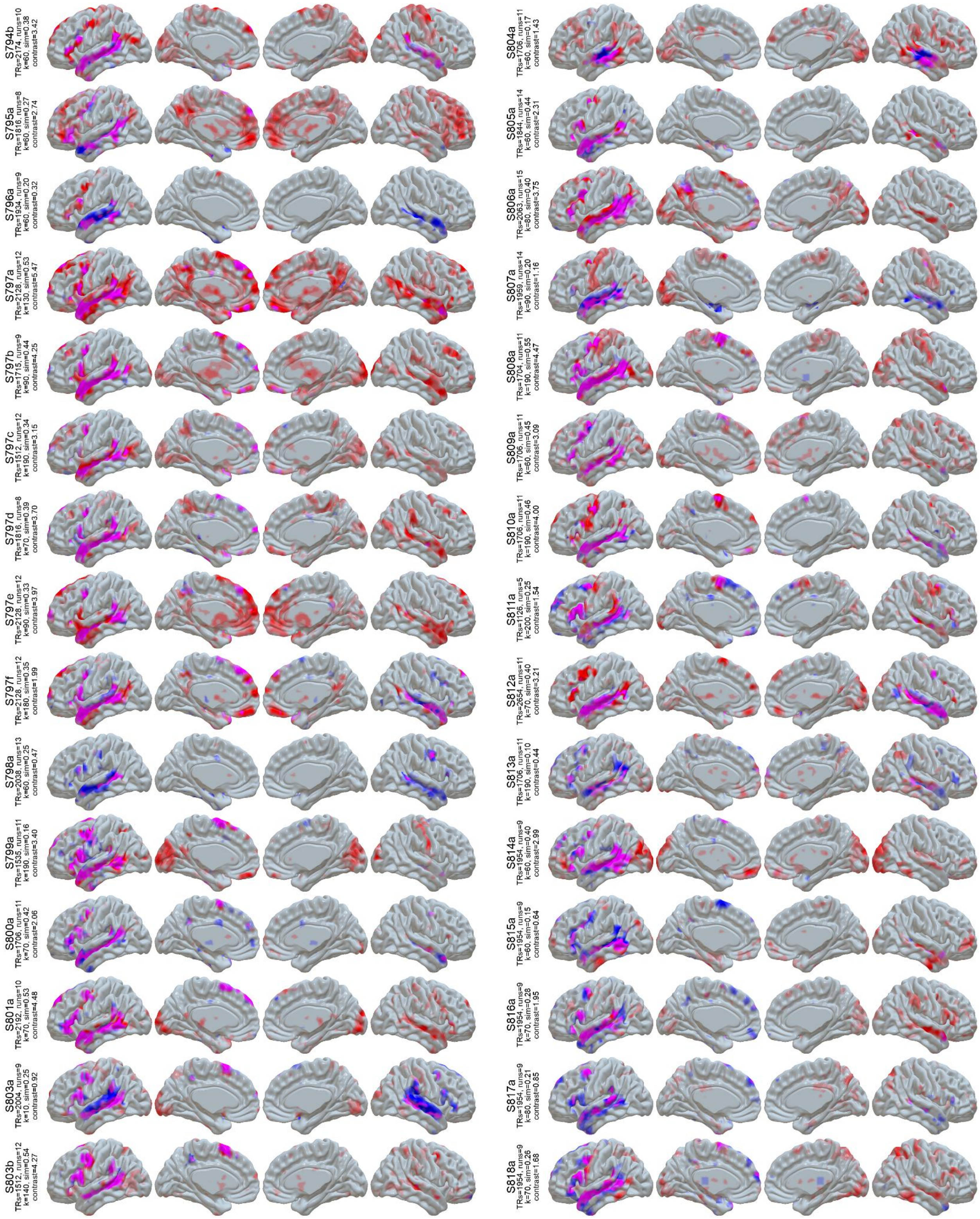


**Figure S1 (cont):** LangFC (blue) vs. task *t*-maps (sentences vs. nonword lists or *S-N*, red) in the 10 sessions with the highest S-N contrast stability between even and odd runs. Overlap is shown in magenta, and opacity reflects magnitude ($0.2 < p < 0.8$ for LangFC, $1 < t < 4$ for S-N).

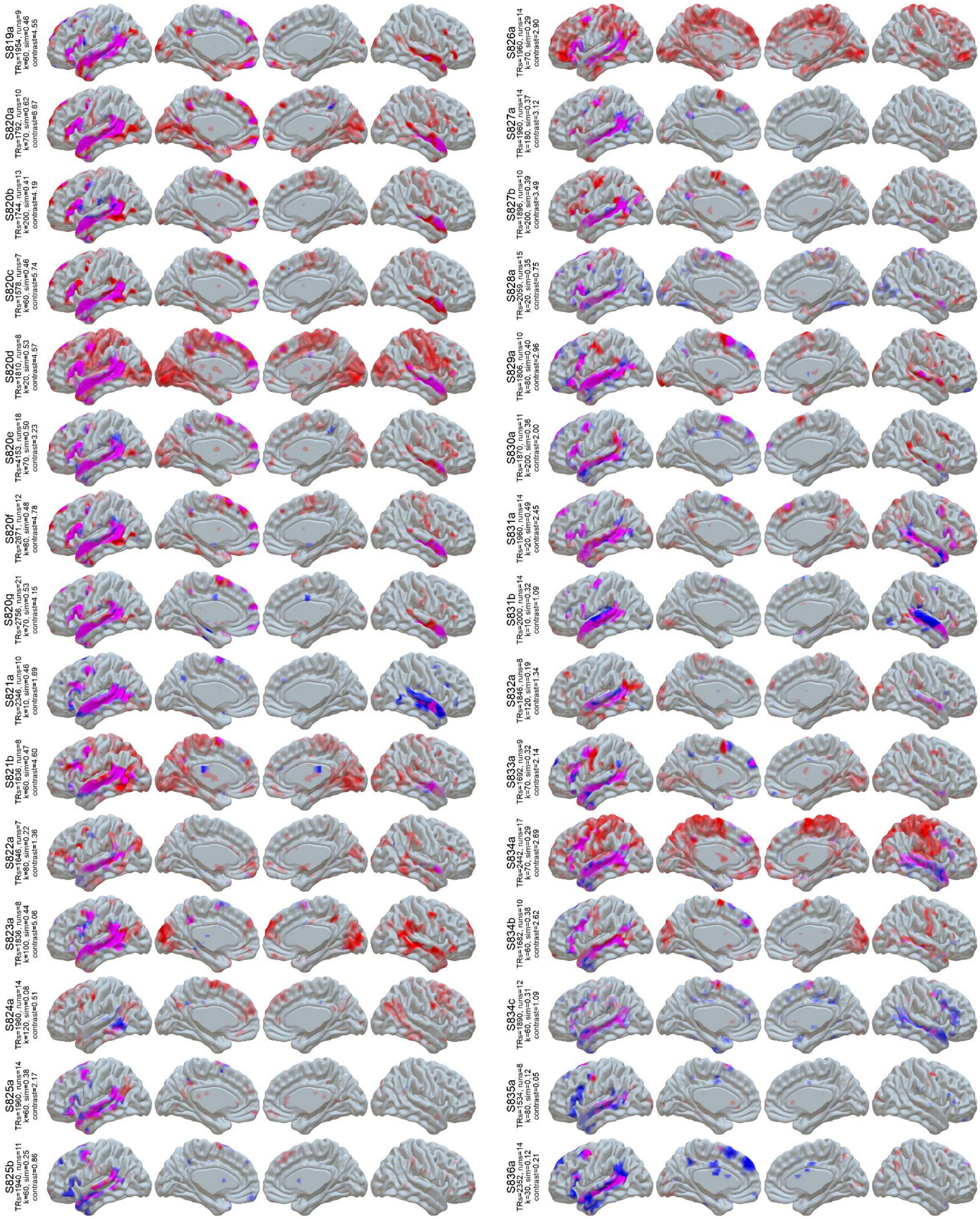


**Figure S1 (cont):** LangFC (blue) vs. task *t*-maps (sentences vs. nonword lists or *S-N*, red) in the 10 sessions with the highest S-N contrast stability between even and odd runs. Overlap is shown in magenta, and opacity reflects magnitude (0.2 < *p* < 0.8 for LangFC, 1 < *t* < 4 for S-N).

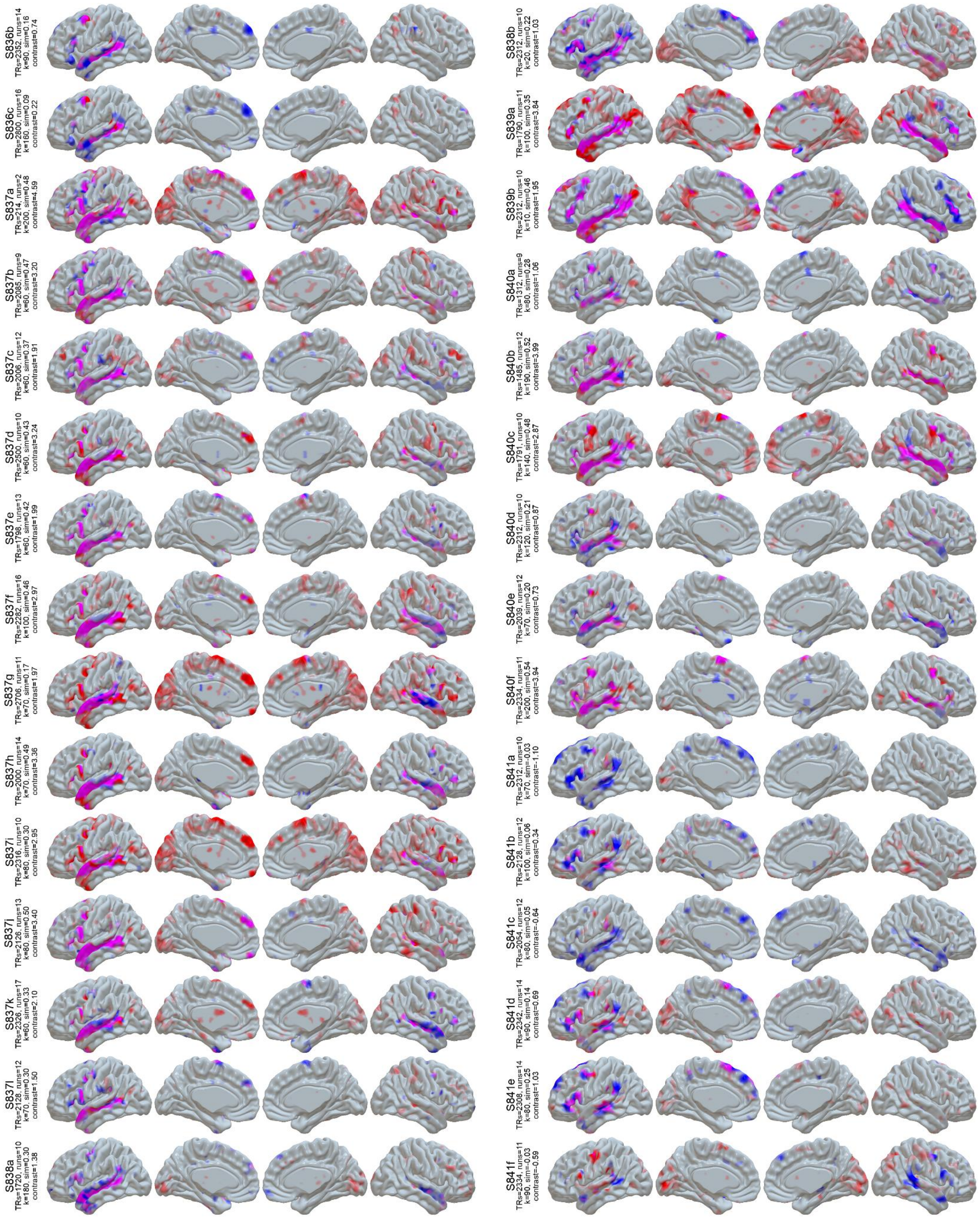

**Figure S1 (cont):** LangFC (blue) vs. task *t*-maps (sentences vs. nonword lists or *S-N*, red) in the 10 sessions with the highest S-N contrast stability between even and odd runs. Overlap is shown in magenta, and opacity reflects magnitude ($0.2 < p < 0.8$ for LangFC, $1 < t < 4$ for S-N).

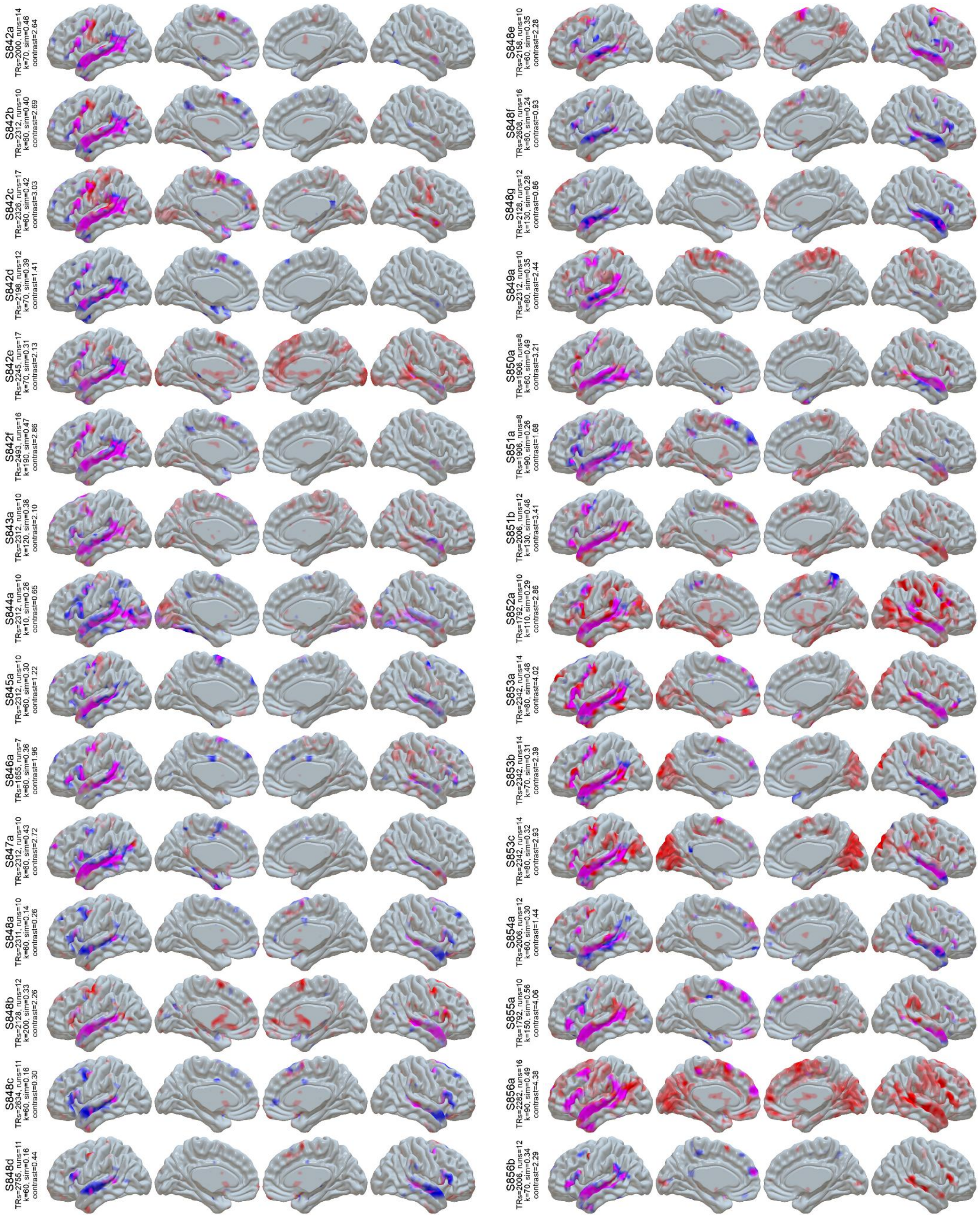


**Figure S1 (cont):** LangFC (blue) vs. task *t*-maps (sentences vs. nonword lists or *S-N*, red) in the 10 sessions with the highest S-N contrast stability between even and odd runs. Overlap is shown in magenta, and opacity reflects magnitude (0.2 < *p* < 0.8 for LangFC, 1 < *t* < 4 for S-N).

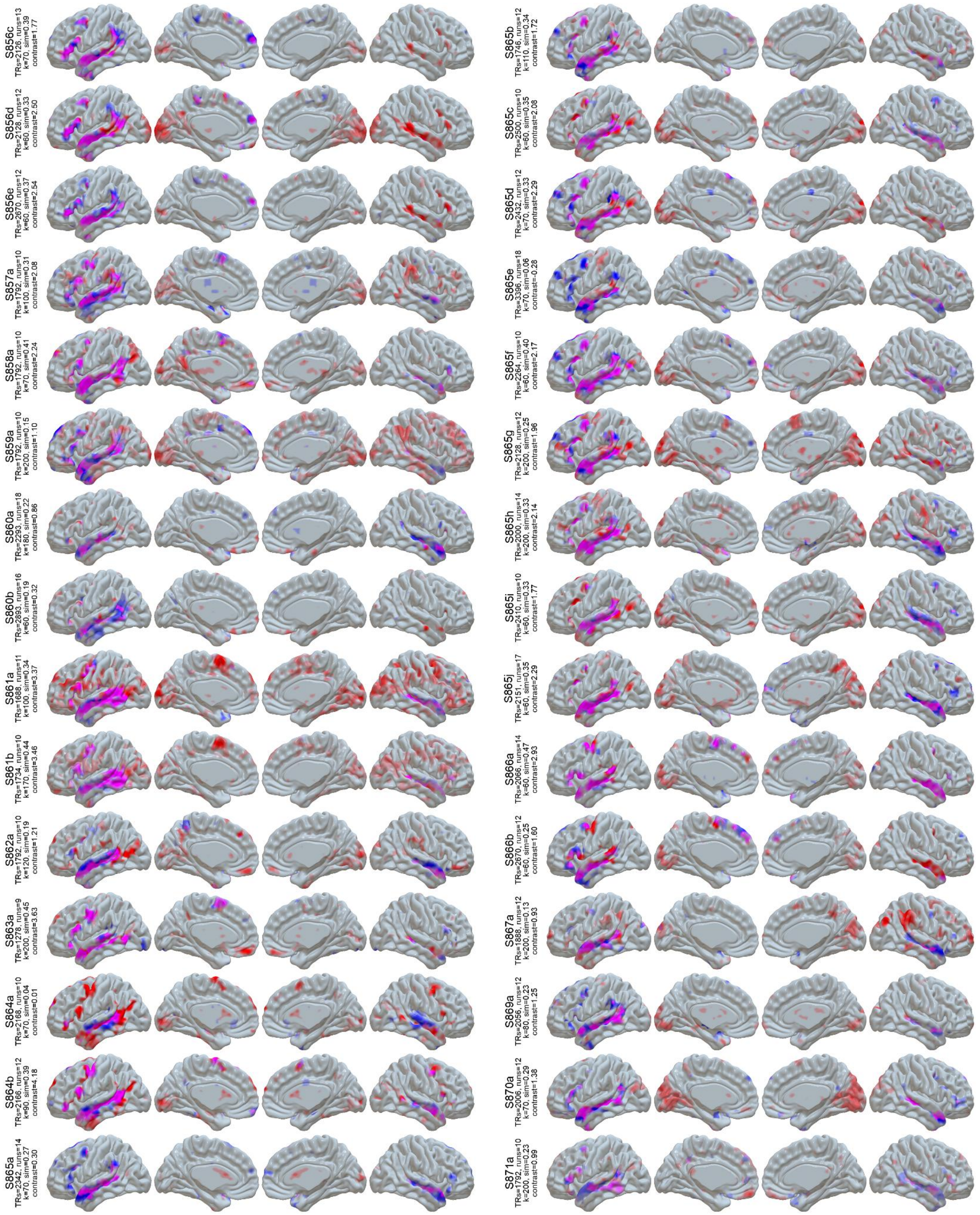


**Figure S1 (cont):** LangFC (blue) vs. task *t*-maps (sentences vs. nonword lists or *S-N*, red) in the 10 sessions with the highest S-N contrast stability between even and odd runs. Overlap is shown in magenta, and opacity reflects magnitude (0.2 < $p$ < 0.8 for LangFC, 1 < $t$ < 4 for S-N).

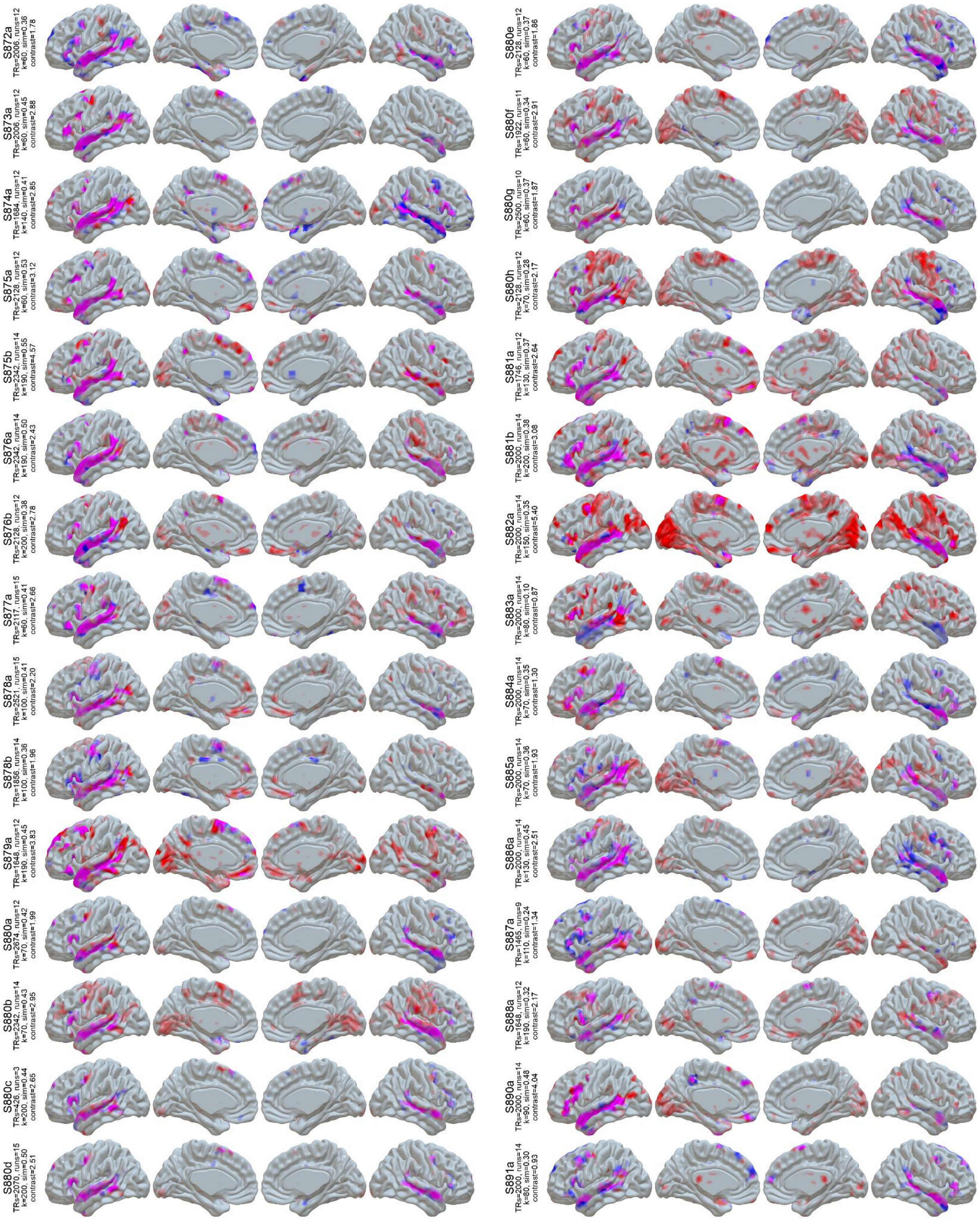


**Figure S1 (cont):** LangFC (blue) vs. task *t*-maps (sentences vs. nonword lists or *S-N*, red) in the 10 sessions with the highest S-N contrast stability between even and odd runs. Overlap is shown in magenta, and opacity reflects magnitude ($0.2 < p < 0.8$ for LangFC, $1 < t < 4$ for S-N).

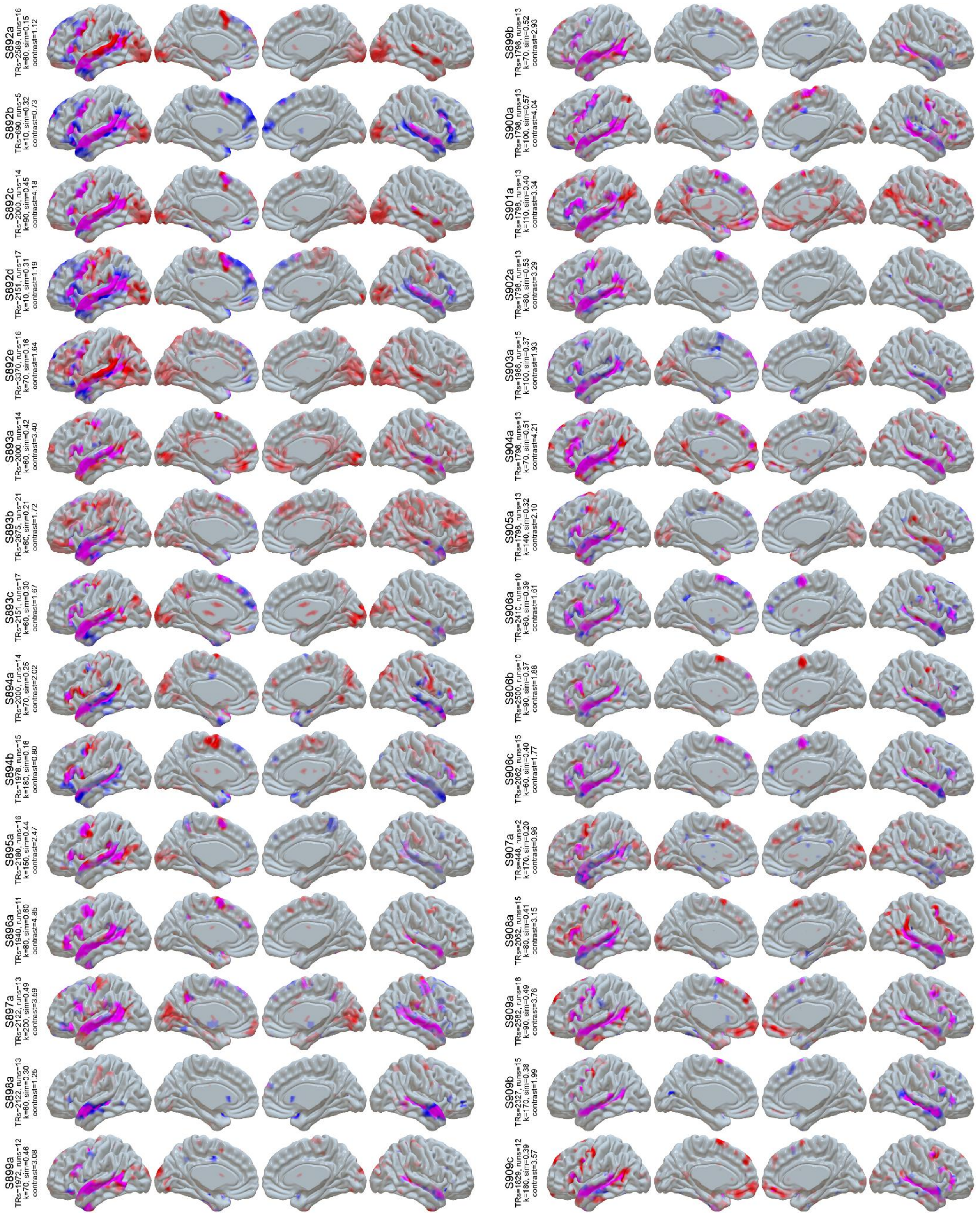


**Figure S1 (cont):** LangFC (blue) vs. task *t*-maps (sentences vs. nonword lists or *S-N*, red) in the 10 sessions with the highest S-N contrast stability between even and odd runs. Overlap is shown in magenta, and opacity reflects magnitude (0.2 < *p* < 0.8 for LangFC, 1 < *t* < 4 for S-N).

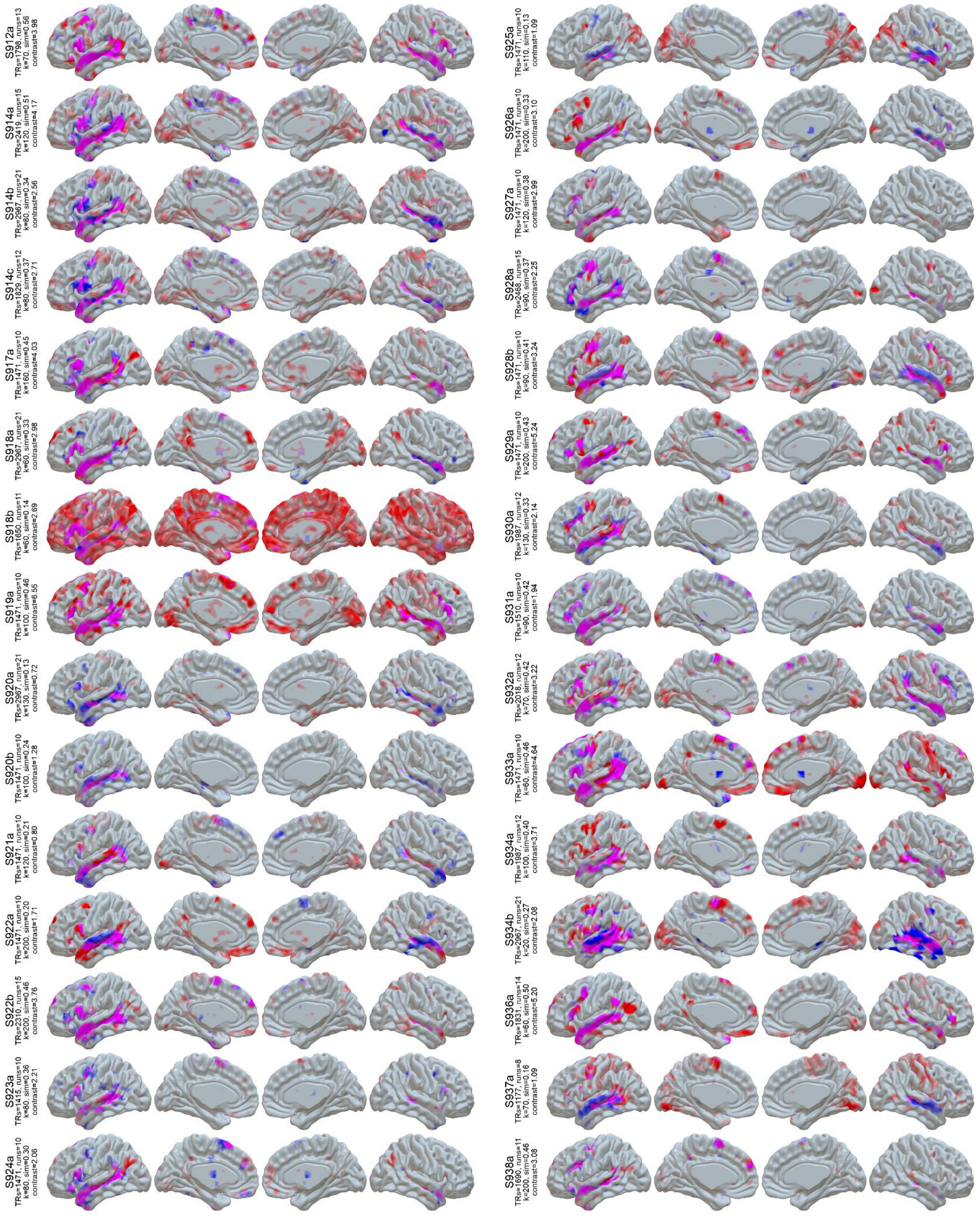

**Figure S1 (cont):** LangFC (blue) vs. task *t*-maps (sentences vs. nonword lists or *S-N*, red) in the 10 sessions with the highest S-N contrast stability between even and odd runs. Overlap is shown in magenta, and opacity reflects magnitude (0.2 < *p* < 0.8 for LangFC, 1 < *t* < 4 for S-N).

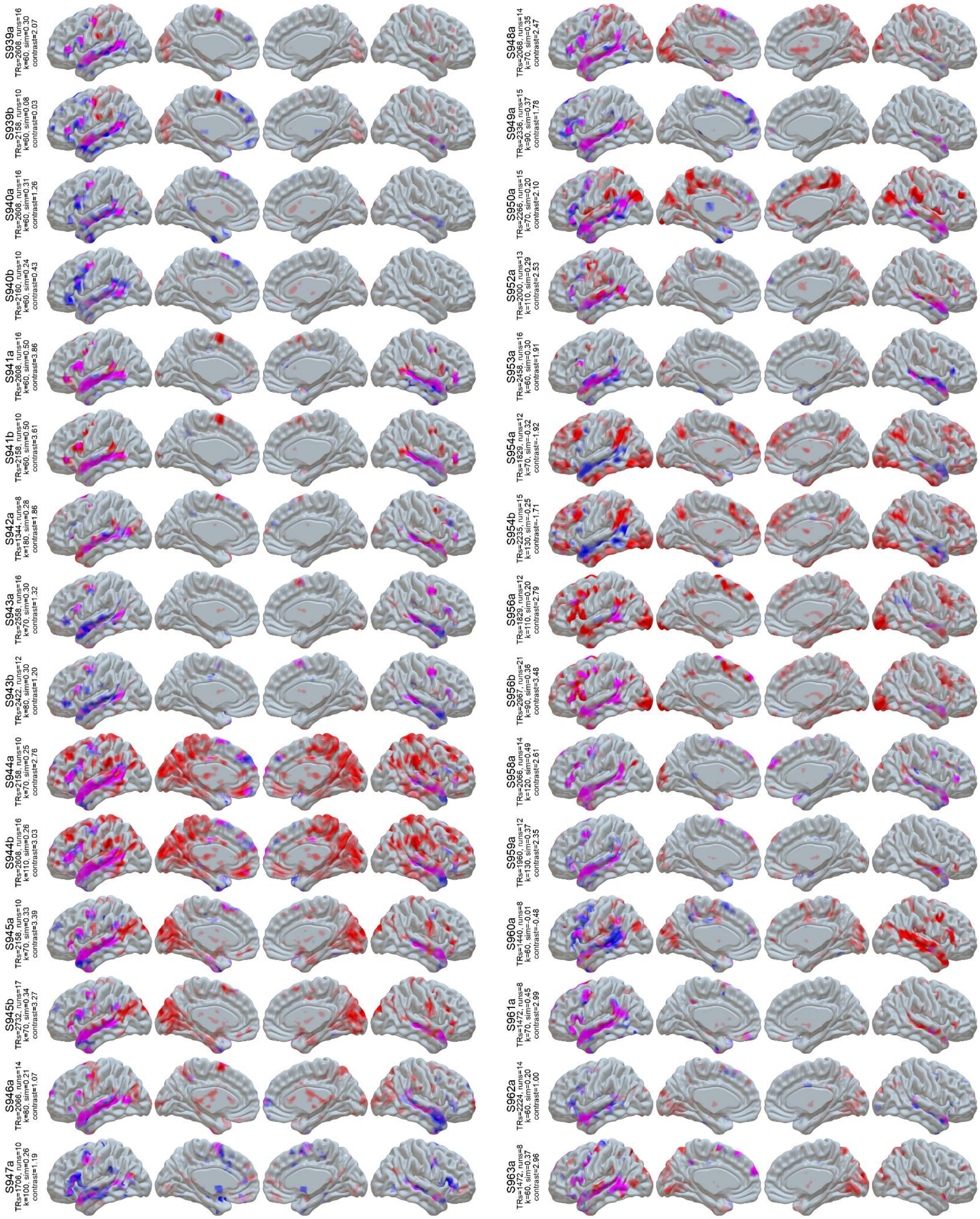


**Figure S1 (cont):** LangFC (blue) vs. task *t*-maps (sentences vs. nonword lists or *S-N*, red) in the 10 sessions with the highest S-N contrast stability between even and odd runs. Overlap is shown in magenta, and opacity reflects magnitude (0.2 < *p* < 0.8 for LangFC, 1 < *t* < 4 for S-N).

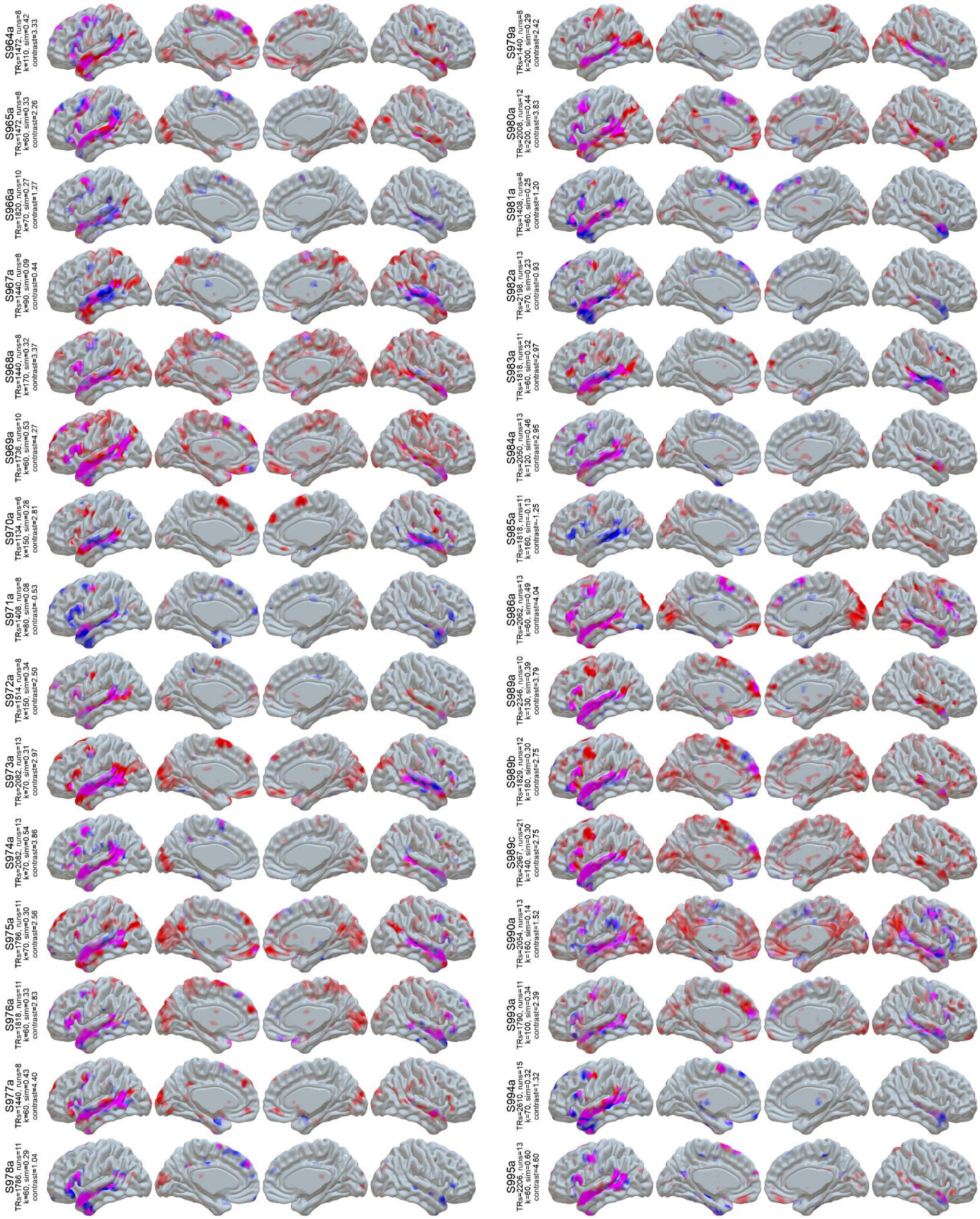


**Figure S1 (cont):** LangFC (blue) vs. task *t*-maps (sentences vs. nonword lists or *S-N*, red) in the 10 sessions with the highest S-N contrast stability between even and odd runs. Overlap is shown in magenta, and opacity reflects magnitude (0.2 < *p* < 0.8 for LangFC, 1 < *t* < 4 for S-N).

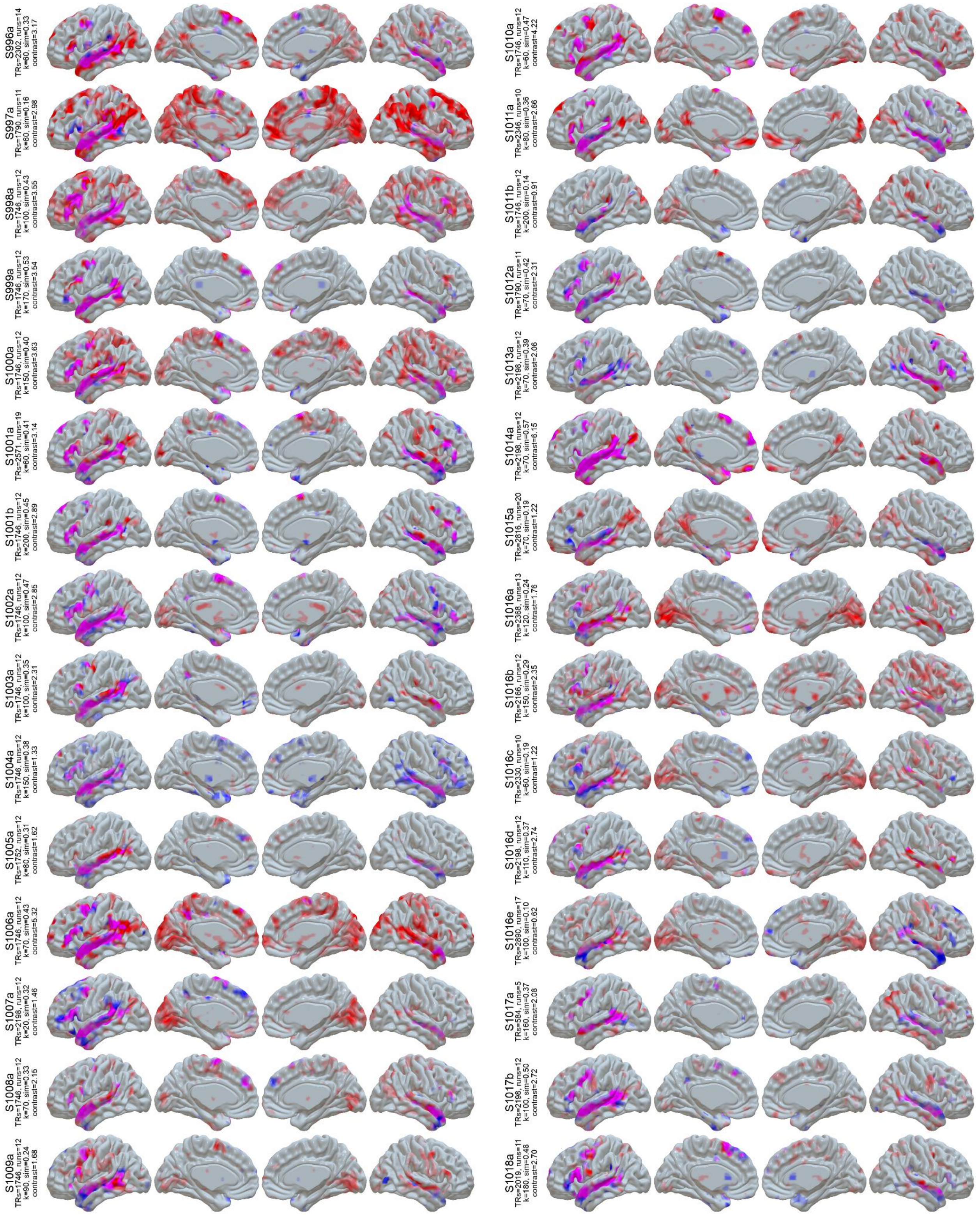


**Figure S1 (cont):** LangFC (blue) vs. task *t*-maps (sentences vs. nonword lists or *S-N*, red) in the 10 sessions with the highest S-N contrast stability between even and odd runs. Overlap is shown in magenta, and opacity reflects magnitude ($0.2 < p < 0.8$ for LangFC, $1 < t < 4$ for S-N).

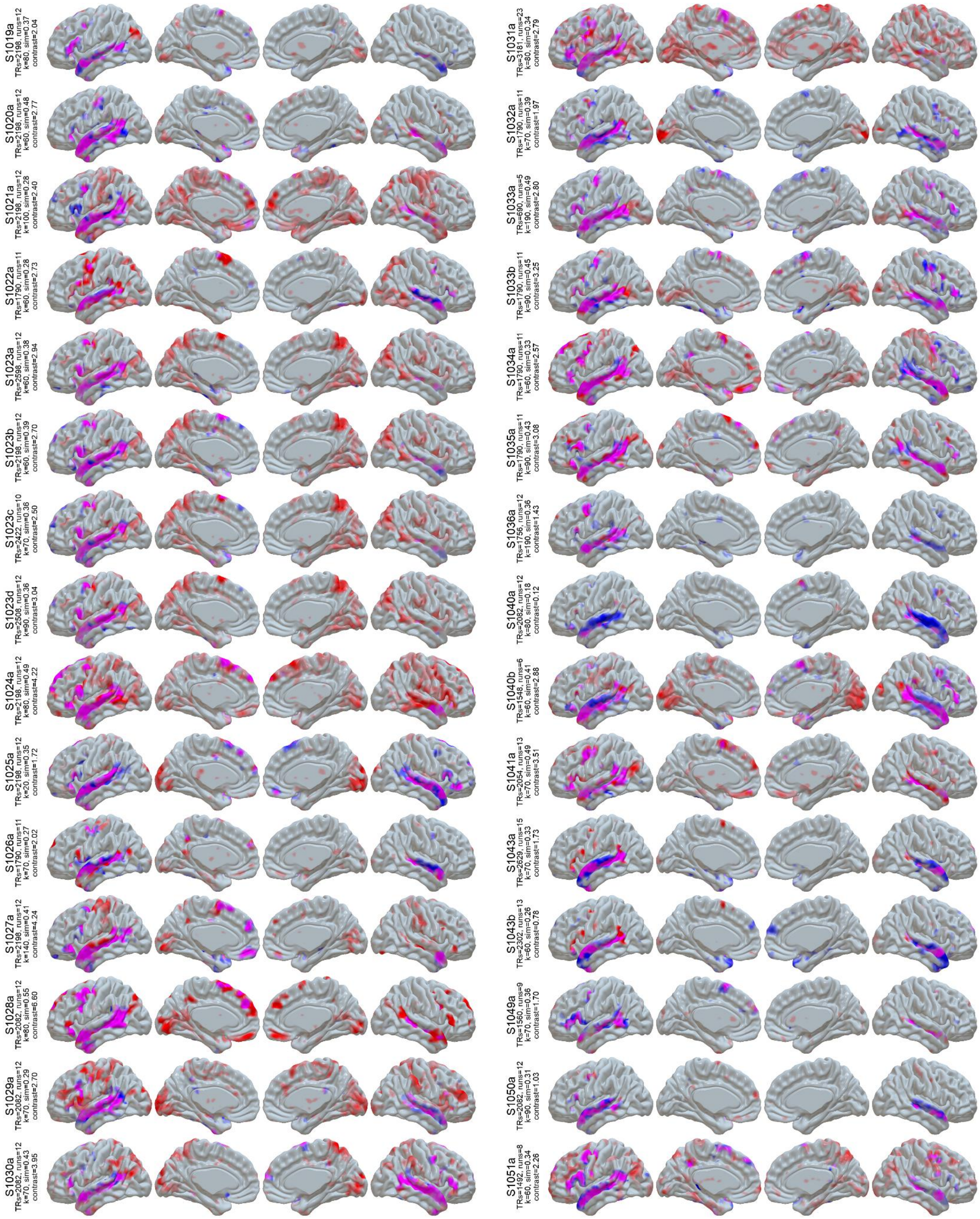


**Figure S1 (cont):** LangFC (blue) vs. task *t*-maps (sentences vs. nonword lists or *S-N*, red) in the 10 sessions with the highest S-N contrast stability between even and odd runs. Overlap is shown in magenta, and opacity reflects magnitude (0.2 < *p* < 0.8 for LangFC, 1 < *t* < 4 for S-N).

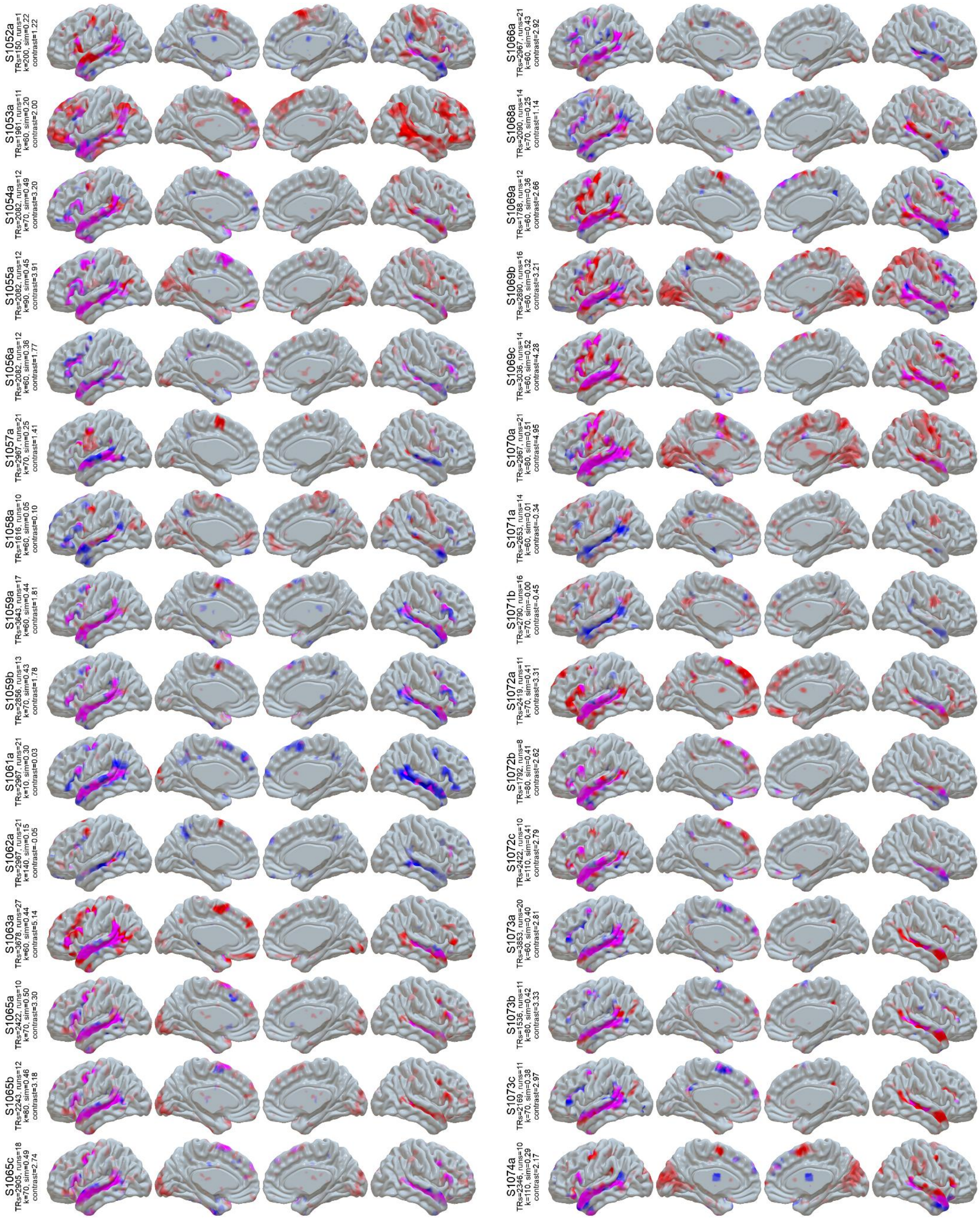


**Figure S1 (cont):** LangFC (blue) vs. task *t*-maps (sentences vs. nonword lists or *S-N*, red) in the 10 sessions with the highest S-N contrast stability between even and odd runs. Overlap is shown in magenta, and opacity reflects magnitude (0.2 < *p* < 0.8 for LangFC, 1 < *t* < 4 for S-N).

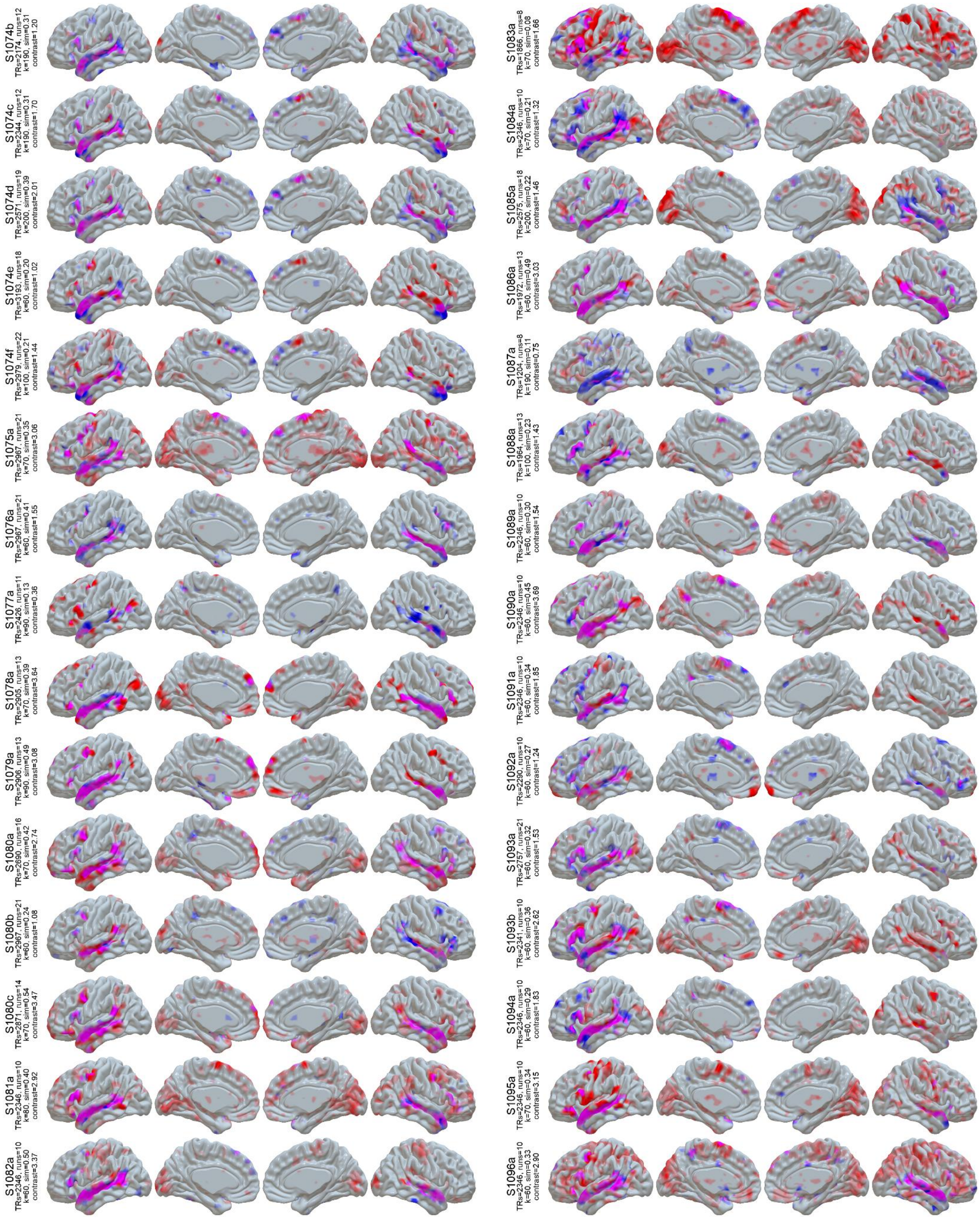


**Figure S1 (cont):** LangFC (blue) vs. task *t*-maps (sentences vs. nonword lists or *S-N*, red) in the 10 sessions with the highest S-N contrast stability between even and odd runs. Overlap is shown in magenta, and opacity reflects magnitude (0.2 < $p$ < 0.8 for LangFC, 1 < $t$ < 4 for S-N).

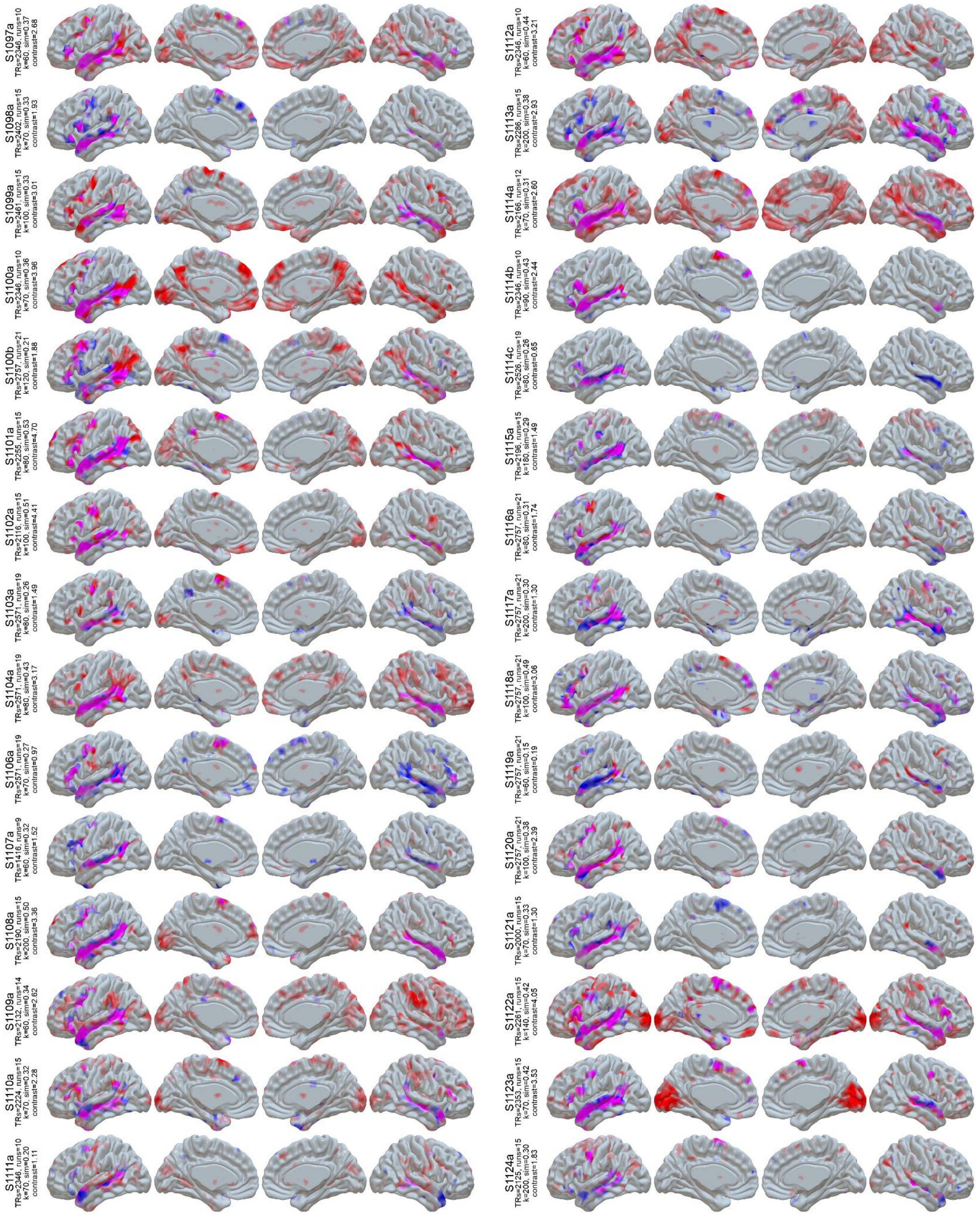


**Figure S1 (cont):** LangFC (blue) vs. task *t*-maps (sentences vs. nonword lists or *S-N*, red) in the 10 sessions with the highest S-N contrast stability between even and odd runs. Overlap is shown in magenta, and opacity reflects magnitude (0.2 < *p* < 0.8 for LangFC, 1 < *t* < 4 for S-N).

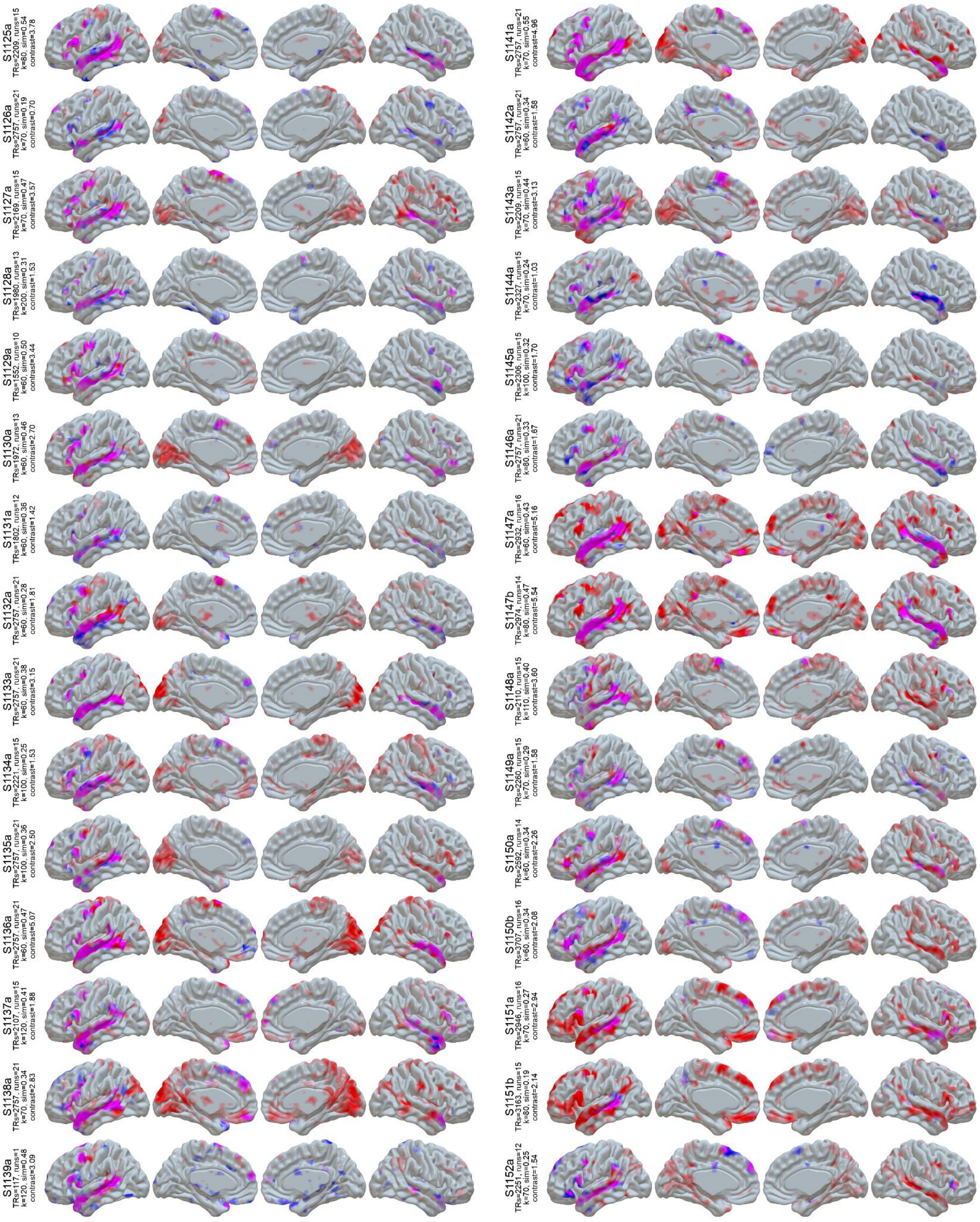


**Figure S1 (cont):** LangFC (blue) vs. task *t*-maps (sentences vs. nonword lists or *S-N*, red) in the 10 sessions with the highest S-N contrast stability between even and odd runs. Overlap is shown in magenta, and opacity reflects magnitude (0.2 < *p* < 0.8 for LangFC, 1 < *t* < 4 for S-N).

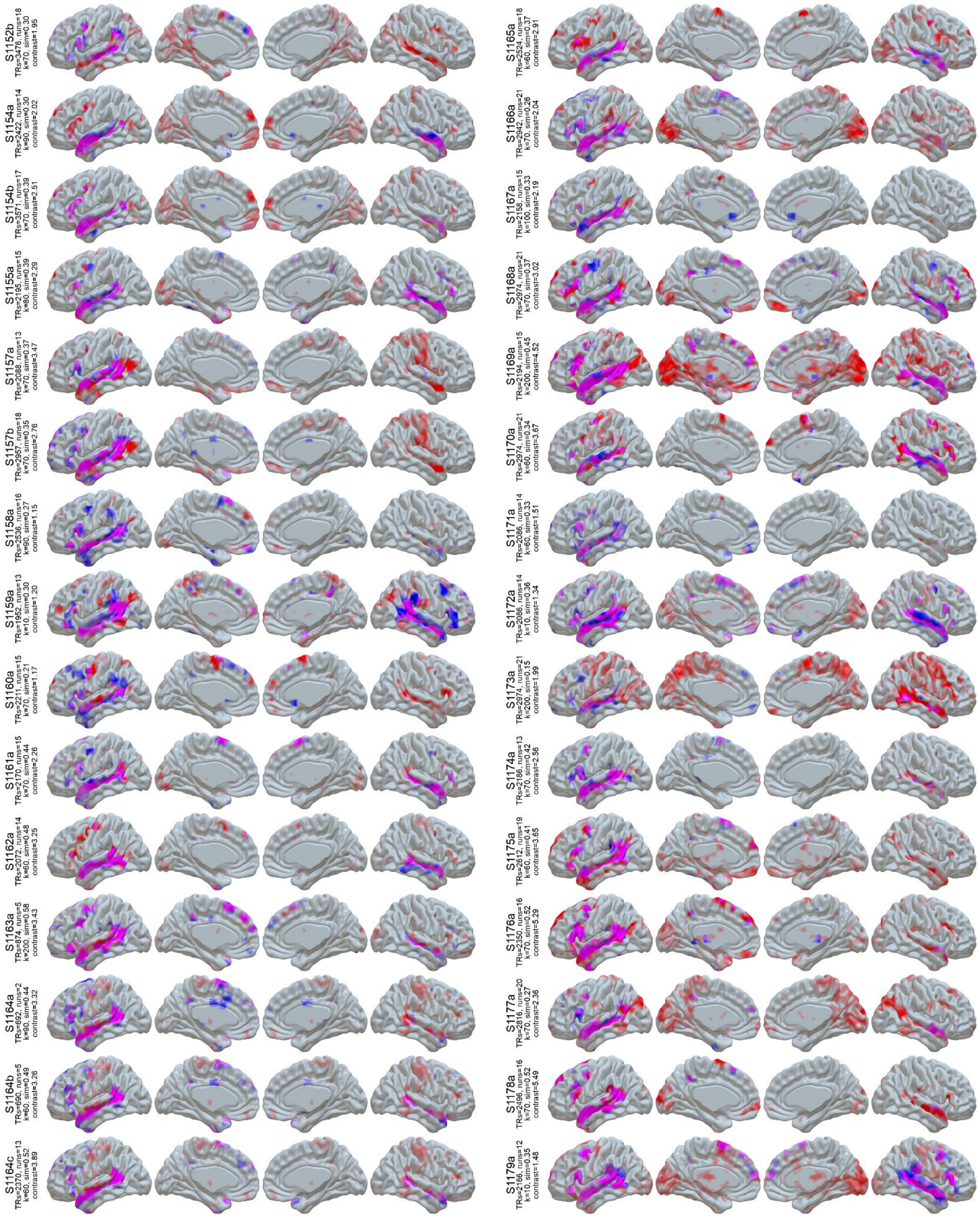


**Figure S1 (cont):** LangFC (blue) vs. task *t*-maps (sentences vs. nonword lists or *S-N*, red) in the 10 sessions with the highest S-N contrast stability between even and odd runs. Overlap is shown in magenta, and opacity reflects magnitude (0.2 < *p* < 0.8 for LangFC, 1 < *t* < 4 for S-N).

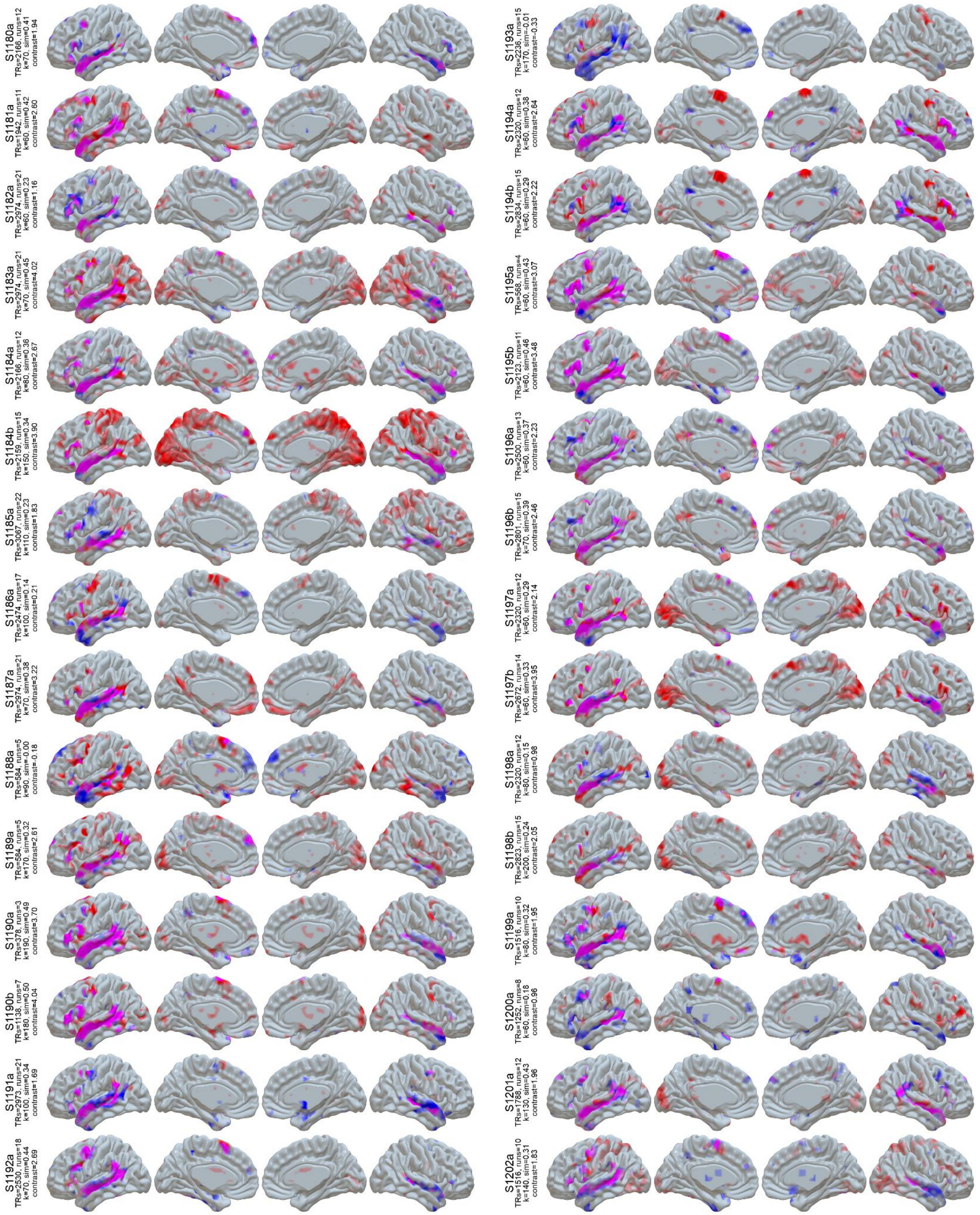


**Figure S1 (cont):** LangFC (blue) vs. task *t*-maps (sentences vs. nonword lists or *S-N*, red) in the 10 sessions with the highest S-N contrast stability between even and odd runs. Overlap is shown in magenta, and opacity reflects magnitude (0.2 < *p* < 0.8 for LangFC, 1 < *t* < 4 for S-N).

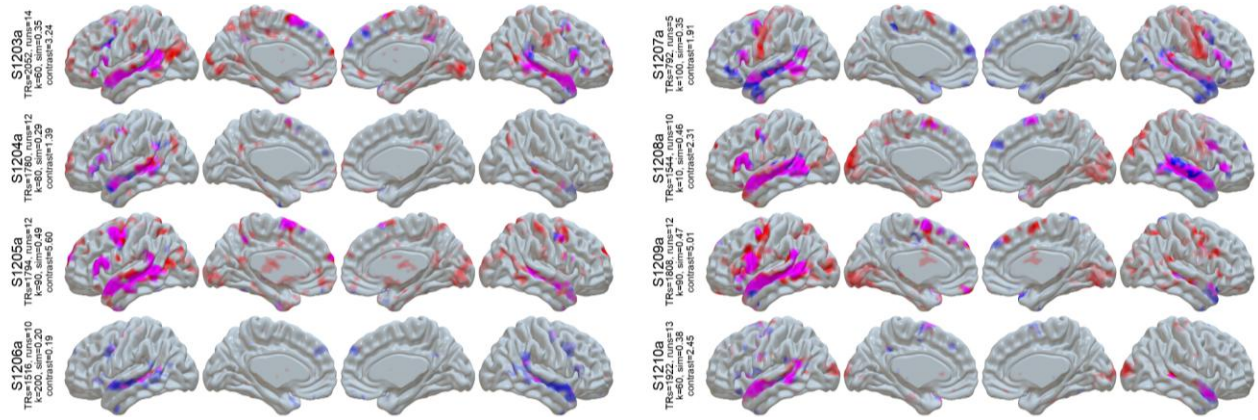


**Figure S1 (cont):** LangFC (blue) vs. task *t*-maps (sentences vs. nonword lists or *S-N*, red) in the 10 sessions with the highest S-N contrast stability between even and odd runs. Overlap is shown in magenta, and opacity reflects magnitude (0.2 < *p* < 0.8 for LangFC, 1 < *t* < 4 for S-N).

### SI B: LangFC Approaches Oracle Performance

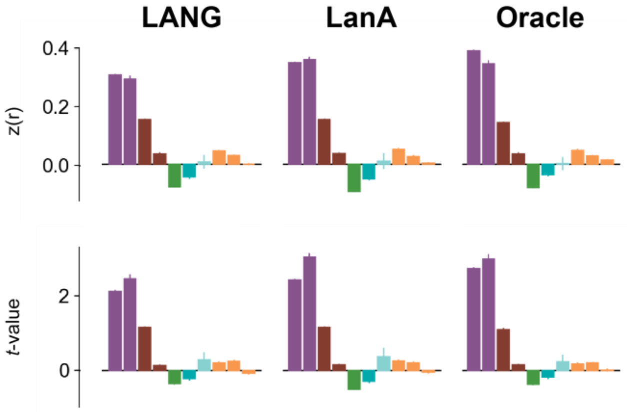


**Figure S2:** Functional tuning of LangFC when labeled by similarity to the LANG (left) or LanA (center) reference atlases, relative to an “oracle” consisting of the reading localizer task contrast map for the individual (right). Task colors parallel those in **Fig 4C** of the main article. Labeling relative to (participant-agnostic) reference atlases yields comparable degree and kind of LangFC tuning relative to the oracle labeling strategy (i.e., directly finding the network that is most spatially similar to the reading localizer task contrast).

We analyzed the performance of our approach to labeling LangFC (i.e., via comparison to reference atlases derived from prior work) by comparing it to an “oracle” setting in which we directly choose the network with the most similar topography to the participant’s own contrast map from the reading-based language localizer (**Fig S2**). We found that the oracle offered little benefit, relative to reference-based labeling, for recovering either the topography (z(r), top) or the task selectivity (*t*-value, bottom) of the localizer contrast maps themselves. In other words, in aggregate, our data-driven labeling method approaches the participant-specific optimum for this dataset.

### SI C: LangFC Is Functionally Similar With and Without Task Regression

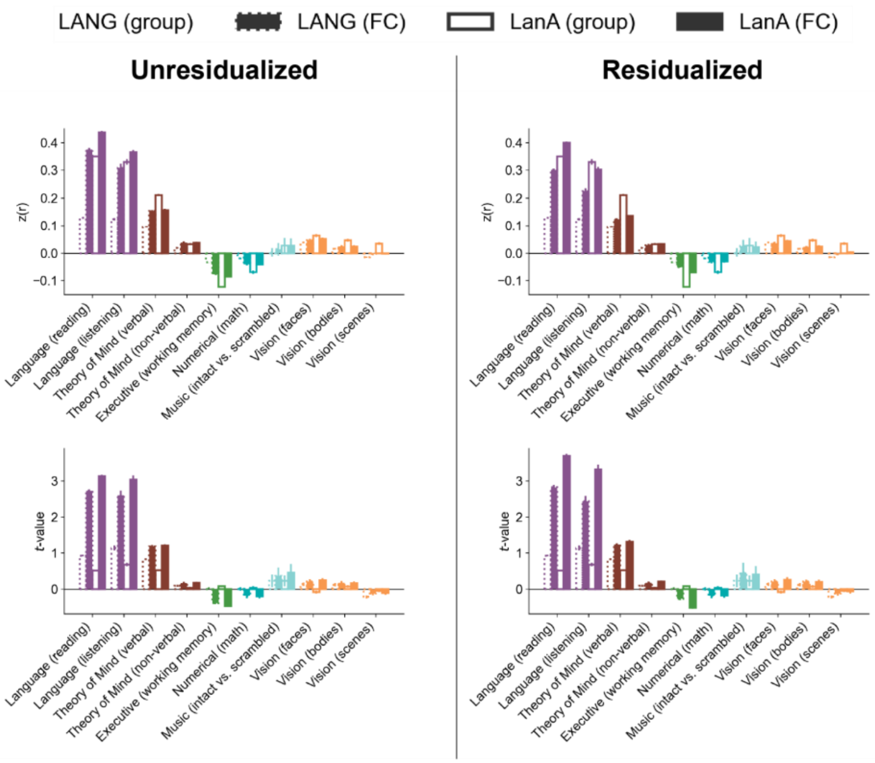


**Figure S3:** Functional tuning of LangFC estimated from reading language localizer task timecourses with (right) and without (left) task regression (76), parallel to **Fig 3A** of the main article. Tuning is largely preserved under task residualization.

We examined how strongly LangFC connectivity could be driven by tasks in an extreme case: the S-N localizer itself, which is known to elicit large responses in language areas (31). This analysis is motivated by two concerns: *first*, FC studies of task data typically regress out the task structure in order to better simulate rest-like connectivity (76), and *second*, it is possible that LangFC's emergence from task data is partially driven by indirectly capturing task contrasts. For example, if regions respond strongly to a task manipulation (like the reading language localizer), they may have inflated functional connectivity due to task-induced correlations. This could limit the practical utility of iFC for task-agnostic language localization. We reasoned that if LangFC is largely conserved with and without task regression from reading localizer runs, then this would mitigate both of these concerns.

To investigate this, in each unique reading localizer session used to estimate S-N contrasts in our study (n=1,479), we fitted a voxelwise generalized linear model (GLM) using a design matrix created from the experiment's task structure. This resulted in two versions of the reading language localizer timecourses in a given session: the inputs to the GLM ("unresidualized") and the residuals of the GLM, i.e., task-regressed timecourses ("residualized"). We then estimated LangFC for each localizer session, with and without residualization. For efficiency reasons, we did not optimize the parcellation granularity by participant (as in the main study) but simply fixed it at $k$=100, one of many values that gave reasonable performance in our main analysis (**Fig 4C**). We found that, within a session, LangFC estimated from unresidualized timecourses was spatially similar to LangFC estimated from residualized timecourses (mean z(r) = 0.75 ± 0.005 SEM). Moreover, in both conditions, we found a strongly language-selective profile for LangFC (**Fig S3**), as in the main study. LangFC's spatial similarity to S-N task contrasts (z(r), top of panel) decreases slightly under residualization, but its S-N response strength ($t$-value, bottom of panel) remains high under task residualization (if anything, $t$ increases under residualization for LangFC labeled against the LanA atlas). Thus, the core topography and functional tuning of the network is preserved across residualization, suggesting that a language network is robustly present in the individualized functional connectome, with or without task regression.

### SI D: LangFC Results Are Robust to Parcellation Design

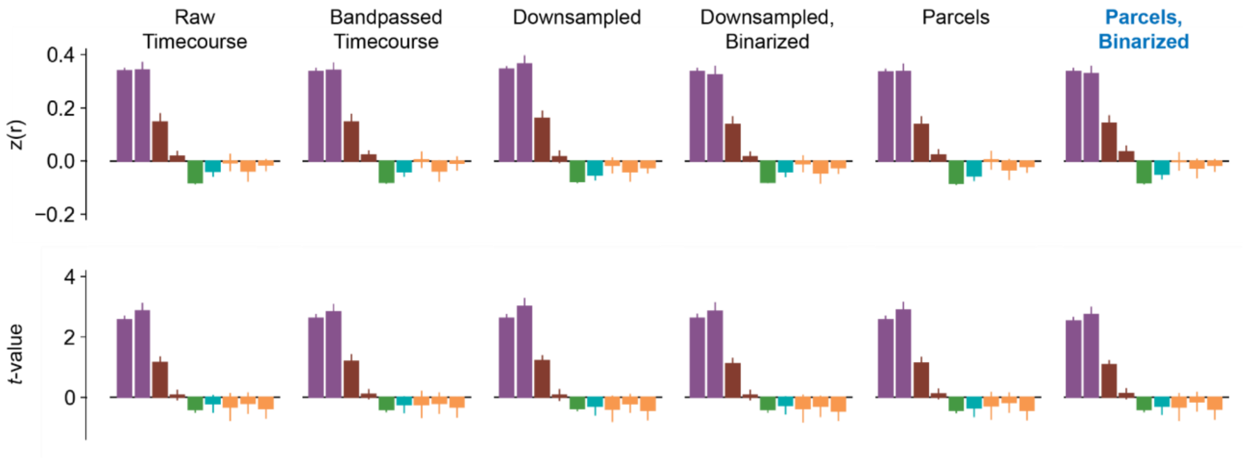


**Figure S4:** Functional tuning of LangFC within a subset of 100 of our fMRI sessions as estimated by six variants of the parcellation approach, parallel to **Fig 4C** of the main article. The rightmost variant (Parcels, Binarized) was the one we used throughout this study. The music task is absent from these plots because (by chance) none of the participants in this subset completed it.

Our method for parcellating individual brains into networks (**Parcellation and Evaluation Procedure**) involved many design choices, some of which were motivated primarily by precedent established by related studies (e.g., parcellating based on binarized connectivity to a regional atlas, (65)). Here we show evidence (**Fig S4**) that results are largely robust to these decisions. To do so, we explored six variants of the parcellation procedure in a subset of 100 of our fMRI sessions spaced roughly evenly across the lab's acquisition timeline. For efficiency reasons, as in **SI C**, we did not optimize the parcellation granularity but simply fixed it at $k$=100. The variants were as follows:

- **Raw Timecourse:** The raw (preprocessed) BOLD timecourses were directly parcellated using $k$-means, with no bandpassing or connectivity computation.

- **Bandpassed Timecourse:** The bandpassed (0.01-0.1 Hz) BOLD timecourses were directly parcellated using *k*-means, with no connectivity computation
- **Downsampled:** The bandpassed BOLD timecourses were first spatially downsampled to 4mm isotropic voxel resolution and the pairwise connectivity between these larger voxels was parcellated using *k*-means.
- **Downsampled, Binarized:** Same as Downsampled, except that the connectivity was first binarized by thresholding at the 90th percentile before being parcellated using *k*-means.
- **Parcels:** Connectivity was computed between the (134,713 gray matter voxel) BOLD timescales and 1,000 larger-scale regions from an existing atlas (53), resulting in a functional connectome with dimension 134,713 × 1,000, which was then parcellated using *k*-means.
- **Parcels, Binarized:** Same as Parcels, except that the voxel-by-region connectome was first binarized by thresholding at the 90th percentile before being parcellated using *k*-means.

The final configuration (Parcels, Binarized) corresponds to the design used throughout this study. The rest are defensible alternatives. Our key finding (**Fig S4**) is that none of these choices matters much to the aggregate picture of LangFC's functional tuning in this sample of participants, which is stable across all six variants. This outcome indicates that iFC provides a reliable window onto language function in the brain across multiple choice points in the analysis design.

### SI E: Results from Our Approach Converge with Template Matching

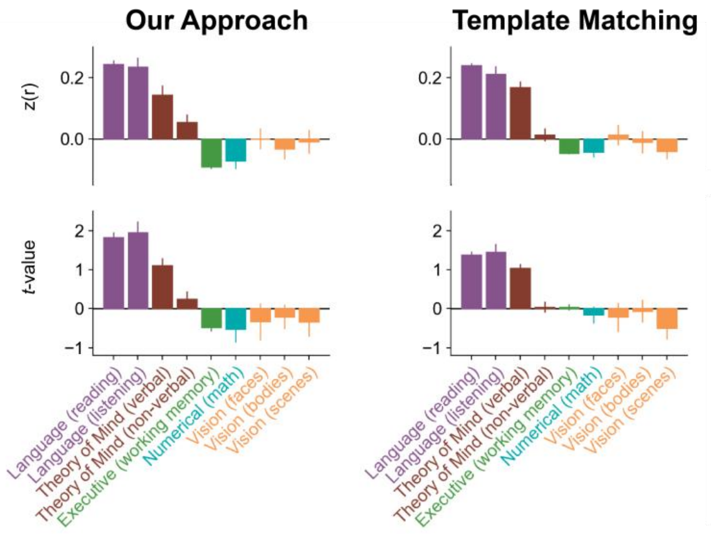


**Figure S5:** Functional tuning of LangFC as estimated by the approach in the main article ("Our Approach") relative to template matching against the Du et al. (2024) group atlases. As in e.g., **Fig 3**, plots show spatial similarity z(r) (above) and weighted-average *t*-value (below) between LangFC and diverse task activation maps in the grey matter volume.

The central contribution of this study is to clarify the nature and function of the brain's language network as revealed by functional connectivity, rather than to advocate a particular algorithm for parcellating the brain into networks. Our method was designed for simplicity and scale, and, as stressed in the main article and supported by **SI D**, we expect our findings to generalize across a range of parcellation methods. In this section, we extend the findings in **SI D** to a substantively different alternative approach known as *template matching* (59, 169–171). In brief, template matching assigns voxels to *a priori* networks by finding the network with the most similar topography to the voxel's connectivity map according to some similarity score. Template matching is not an appropriate method for our core goal of testing the language network hypothesis from functional connectivity, due to potential circularity: template-matching parcellations cannot be used to validate the existence of the networks that were assumed by the parcellation procedure. The sampling approach we used to derive our main results differs critically in this respect: our parcellations are computed strictly bottom-up from connectivity, and *a priori* networks are only used to assign labels to these parcellations *post hoc*.

Nonetheless, the degree to which our parcellations may be shaped by this deliberate *neglect* of prior expectations about functional brain networks is unknown, and we therefore conducted a direct comparison to template matching. To ensure maximal comparability to our main parcellation method, we used the same approach as in the main study to compute a binarized connectivity map in the Ref. (53) 1,000-region atlas for each grey matter voxel. We then assigned each voxel to the *a priori* network from Ref. (65) that had the largest spatial similarity (Pearson correlation in the grey matter volume) to the voxel's connectivity map. We then analyzed the correspondence of the resulting networks to diverse task data, as in the main study. For computational efficiency, we restricted this analysis to the 100 sessions analyzed in **SI D**. Results are presented in **Fig S5** ("Template Matching") as compared to the results from our chosen "Parcels, Binarized" method ("Our Approach") from **SI D**. As shown, both methods produce LangFC estimates with qualitatively similar functional tuning (and strong language selectivity). Template matching is if anything slightly worse than our method, which achieves slightly higher spatial similarity z(r) and effect size *t* for language tasks. Although we leave detailed follow-up to future work, this outcome is plausibly because template matching must by definition assign all voxels to one of the (fifteen) networks in the a priori parcellation and therefore (unlike our approach) cannot consider either graded strength of evidence for network membership between voxels or the possibility of additional networks not anticipated by the prior atlas. Again, our purpose in drawing this comparison is not to adjudicate between our approach and template matching in a general sense, but merely to stress that our method is not materially distorting the conclusions we have drawn or the claims we have advocated, relative to an established alternative.

### SI F: iFC Parcellations Are Similar Across Task and Rest

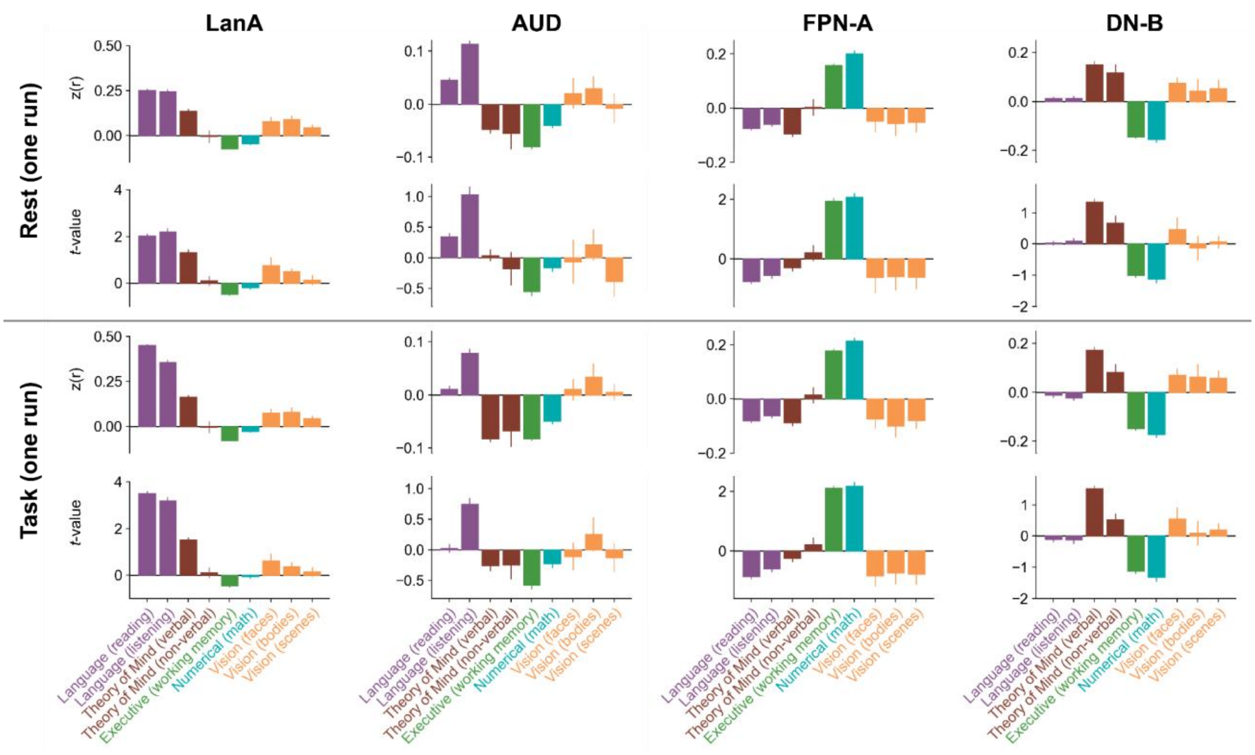


**Figure S6:** Functional tuning of LangFC as estimated from a single resting-state run (top-half) vs. a single run of the reading language task (sentences vs. nonwords, bottom half), for four different a priori networks: language (LanA), auditory (AUD), multiple-demand (FPN-A) and theory-of-mind (DN-B). As in e.g., **Fig 3**, plots show spatial similarity z(r) (above) and weighted-average *t*-value (bottom) between LangFC and diverse task activation maps in the grey matter volume.

A central argument of this study is that iFC can be applied retrospectively to identify the language network from arbitrary task data. As stressed in the main article, this finding unlocks new avenues for precision neuroimaging of language function, extending to both existing fMRI datasets in which resting state or language localizer task data were not collected, as well as future fMRI data collection, especially in populations for which it may be infeasible to collect additional rest or task data for localization purposes. Importantly, we are not taking a position in ongoing debates about whether task or rest data are better for connectivity analysis in general (68–70, 172). We are simply highlighting the feasibility of iFC-based language network identification from arbitrary fMRI data, in line with recent arguments by others (66).

Nonetheless, given the fundamental role played by resting state data in what is currently known about functional connectivity, we sought to study this relationship directly. Our dataset is not optimally suited to this: it contains comparatively little resting data, and no more than one resting state run per scanning session. To address this, we conducted the closest approximation we could design to a minimal comparison between rest and task. The most frequently-used resting state configuration in our dataset was 300s of resting data at a TR of 2s, or 150 TRs. This configuration is reasonably similar in duration and acquisition parameters to the most frequently-used task state configuration in our dataset: a single 358s run of the reading localizer task (sentences vs. nonword lists), also at a TR of 2s, or 179 TRs. Because this task is one of the tasks used to evaluate the language selectivity of the parcellations, it can be viewed as a rough upper bound on the amount of signal about the language network that can be derived from connectivity estimated from 6-7 minutes of task data. We identified all sessions in the main study in which participants completed both of these configurations, resulting in a total of 189 sessions. We then parcellated these sessions using the same technique as in the main article and compared the resulting networks to diverse task maps in order to study functional tuning. The one deviation from our standard approach that was required for this analysis was to reduce the number of spatial principal components from 200 to 100 prior to computing connectivity, since neither of these tasks contain the 200 TRs needed to compute 200 components. We also removed the music task from the evaluation, since it was only used in two of the 189 sessions.

Results are given in **Fig S6**. In order to assess the impact of the rest vs. task distinction on network estimation in general (not just estimation of the language network), we also included the three other *a priori* networks from **Fig 3** of the main article whose activations are plausibly modulated by our task set: the auditory network (AUD), the frontoparietal/multiple-demand network (FPN-A), and the default/theory-of-mind network (DN-B). As shown, the functional tuning of the resulting networks is highly similar whether they are estimated from rest (top half) or task (bottom half), suggesting that connectivity patterns in both rest and task states paint a similar picture of these four large-scale brain networks. The principal exception is that LangFC has substantially higher spatial similarity z(r) and task response *t* in language tasks (purple) when estimated from task as opposed to rest, especially in the reading language task (but also to some extent in the listening language task). This outcome for the reading task is unsurprising given that (unlike in our main study) the same data was used in the task (but not the rest) condition to estimate both reading task responses and connectivity. Nonetheless, although the results in Fig S6 reveal some effect of task, they largely align with prior claims that the functional connectome is highly conserved across task states (72), enough to warrant the use of functional network identification from task-based connectivity in a wide range of cases.
